# Multi-omics profiling of Earth’s biomes reveals patterns of diversity and co-occurrence in microbial and metabolite composition across environments

**DOI:** 10.1101/2021.06.04.446988

**Authors:** Justin P. Shaffer, Louis-Félix Nothias, Luke R. Thompson, Jon G. Sanders, Rodolfo A. Salido, Sneha P. Couvillion, Asker D. Brejnrod, Franck Lejzerowicz, Niina Haiminen, Shi Huang, Holly L. Lutz, Qiyun Zhu, Cameron Martino, James T. Morton, Smruthi Karthikeyan, Mélissa Nothias-Esposito, Kai Dührkop, Sebastian Böcker, Hyun Woo Kim, Alexander A. Aksenov, Wout Bittremieux, Jeremiah J. Minich, Clarisse Marotz, MacKenzie M. Bryant, Karenina Sanders, Tara Schwartz, Greg Humphrey, Yoshiki Vásquez-Baeza, Anupriya Tripathi, Laxmi Parida, Anna Paola Carrieri, Kristen L. Beck, Promi Das, Antonio González, Daniel McDonald, Søren M. Karst, Mads Albertsen, Gail Ackermann, Jeff DeReus, Torsten Thomas, Daniel Petras, Ashley Shade, James Stegen, Se Jin Song, Thomas O. Metz, Austin D. Swafford, Pieter C. Dorrestein, Janet K. Jansson, Jack A. Gilbert, Rob Knight, the Earth Microbiome Project 500 (EMP500) Consortium

## Abstract

As our understanding of the structure and diversity of the microbial world grows, interpreting its function is of critical interest for understanding and managing the many systems microbes influence. Despite advances in sequencing, lack of standardization challenges comparisons among studies that could provide insight into the structure and function of microbial communities across multiple habitats on a planetary scale. Technical variation among distinct studies without proper standardization of approaches prevents robust meta-analysis. Here, we present a multi-omics, meta-analysis of a novel, diverse set of microbial community samples collected for the Earth Microbiome Project. We include amplicon (16S, 18S, ITS) and shotgun metagenomic sequence data, and untargeted metabolomics data (liquid chromatography-tandem mass spectrometry and gas chromatography mass spectrometry), centering our description on relationships and co-occurrences of microbially-related metabolites and microbial taxa across environments. Standardized protocols and analytical methods for characterizing microbial communities, including assessment of molecular diversity using untargeted metabolomics, facilitate identification of shared microbial and metabolite features, permitting us to explore diversity at extraordinary scale. In addition to a reference database for metagenomic and metabolomic data, we provide a framework for incorporating additional studies, enabling the expansion of existing knowledge in the form of a community resource that will become more valuable with time. To provide examples of applying this database, we outline important ecological questions that can be addressed, and test the hypotheses that every microbe and metabolite is everywhere, but the environment selects. Our results show that metabolite diversity exhibits turnover and nestedness related to both microbial communities and the environment. The relative abundances of microbially-related metabolites vary and co-occur with specific microbial consortia in a habitat-specific manner, and highlight the power of certain chemistry – in particular terpenoids – in distinguishing Earth’s environments.

---

A major goal in microbial ecology is to understand dynamics surrounding the structure and function of microbial communities, how this relates to their taxonomic and phylogenetic composition, and how those relationships vary across space and time. As any single study is not able to sample all environments repeatedly over time to allow for such inferences, fostering the use of standardized methods for studying microbial communities that permit meta-analysis across distinct sampling efforts and studies is of utmost importance ^1–4^. Initial efforts focused on standardized protocols for 16S ribosomal RNA (rRNA) sequencing of bacterial/archaeal communities provided insight into the how communities structure in the environment, supporting strong axes of separation of microbes along gradients of host-association and salinity ^1, 5^. More recent efforts focused on shotgun metagenomics data ^6–9^ have begun to provide additional insight regarding functional potential across environments ^10–14^, and the current state-of-the-art methods employ multi-omics approaches including metagenomics, transcriptomics, proteomics, and/or metabolomics ^15–24^.

Microbes produce diverse secondary metabolites that perform vital functions from communication to defense ^25–27^ and can benefit human health and environmental sustainability ^28–34^. Whereas metagenome mining and transcriptomics are powerful ways to characterize function in microbial communities ^10, 14, 24^, a more powerful approach to understanding functional diversity is to generate chemical evidence that confirms the presence of metabolites ^19–21^ and accurately describe their distribution across the Earth. Here, we present an approach that directly assesses the presence and relative abundance of metabolites, and that provides an accurate description of metabolite profiles in microbial communities across the Earth’s environments. Although several studies have previously employed tandem metagenomics and metabolomics ^22, 23, 35–40^, many existing studies employed relatively limited technical methods or profiled a relatively small number of classes of metabolites ^23, 35, 40^, preventing comparison across studies that could expand our understanding. Further, several previous studies are limited in scope to a single environment or habitat ^20, 23, 24, 35–39^. Our work goes substantially beyond what has been reported previously regarding multi-omics analysis of microbial communities using metagenomics and metabolomics, by including multiple ecosystems. The approach we apply complements metagenomics with a direct survey of secondary metabolites using untargeted metabolomics.

Liquid chromatography with untargeted tandem mass spectrometry (LC-MS/MS) is a versatile method that detects tens-of-thousands of metabolites in biological samples ^19^. Although LC-MS/MS metabolomics has historically suffered from low metabolite annotation rates when applied to non-model organisms, recent computational advances can systematically assign chemical classes to metabolites using their fragmentation spectra ^45^. Untargeted mass-spectrometry-based metabolomics provides the relative abundance (i.e., intensity) of each metabolite detected across samples rather than just counts of unique structures (i.e., presence/absence data), and thus provides a direct readout of the surveyed environment, complementing a purely genomics-based approach. Although there is a clear need to use untargeted metabolomics to quantify the metabolic activities of microbiota, this methodology has been limited by the challenge of distinguishing the secondary metabolites produced exclusively by microbes from other compounds detected in the environment (e.g., those produced by multicellular hosts). To resolve this bottleneck, we devised a computational method for recognizing and annotating putative secondary metabolites of microbial origin from fragmentation spectra. The annotations were first obtained from spectral library matching and *in silico* annotation ^46^ using the GNPS web-platform ^47^. These annotations were then queried against microbial metabolite reference databases (i.e., Natural Products Atlas ^48^ and MIBiG ^49^), and molecular networking ^50^ was used to propagate the annotation to similar metabolites. Finally, a global chemical classification of these metabolites was achieved using a state-of-the-art annotation pipeline (i.e., SIRIUS) ^46^.

We used this methodology to quantify microbial secondary metabolites from diverse microbial communities from the Earth Microbiome Project (EMP, http://earthmicrobiome.org). The EMP was founded in 2010 to sample Earth’s microbial communities at unprecedented scale, in part to advance our understanding of biogeographic processes that shape community structure. To avoid confusion with terminology, we define ‘microbial community’ as consisting of members of the domains Bacteria and Archaea. To build on the first meta-analysis of the EMP archive focused on profiling bacterial and archaeal 16S rRNA ^1^, we crowdsourced a new set of roughly 900 samples from the scientific community specifically for multi-omics analysis. We expanded the scalable framework of the EMP to include standardized methods for shotgun metagenomic sequencing and untargeted metabolomics for cataloging microbiota globally. As a result, we provide a rich resource for addressing outstanding questions and to serve as a benchmark for acquiring new data. To provide an example for using this resource, we present a multi-omics, meta-analysis of this new sample set, tracking not just individual sequences but also genomes and metabolites. Our analysis includes diverse studies with sample types classified using an updated and standardized environmental ontology, describes large-scale ecological patterns, and explores important questions in microbial ecology.

Specifically, we explore the hypothesis that, “everything is everywhere, but the environment selects” ^51–55^. We predict that although most major classes of metabolites have cosmopolitan distributions ^14^, their relative abundances will vary strongly among different environments. Therefore, whereas the presence/absence of metabolites alone may show profiles that are relatively uniform across samples, their relative abundances will provide great power in distinguishing among habitats. We predict that similar to microbes ^1^, metabolites will exhibit both turnover and nestedness across habitats. Furthermore, we expect variation in metabolite profiles among environments to be in part driven by variation in microbial community composition. Therefore, we explore the hypothesis that metabolite alpha- and beta-diversity will be strongly correlated with microbial diversity. We anticipate strong, positive relationships between microbial diversity and metabolite diversity, but that environmental similarity based on microbial composition may be distinct from that based on metabolite composition. We suspect that this is in part due to deterministic processes unique to microbial community assembly, and similarity in metabolite profiles across the microbial phylogeny ^56–58^. Regardless, if profiles for metabolites and microbes are habitat-specific, we predict that certain features can be used to classify samples among environments. We also predict that metabolites will co-occur with specific microbial taxa such that metabolite–microbe co-occurrences can be described as features in the environment that define specific habitats.

## RESULTS

### A resource for a meta-analytical- and multi-omics approach to microbial ecological research

Here, we generated data for 880 environmental samples that span 19 major environments contributed by 34 principal investigators as part of the Earth Microbiome Project 500 (EMP500). The EMP500 is a novel sample set for multi-omics protocol development and data exploration (Fig. 1; Table S1). To normalize sample collection for this- and future studies, we updated and followed the existing Earth Microbiome Project (EMP) Sample Submission Guide (https://earthmicrobiome.org/protocols-and-standards/emp500-sample-submission-guide/) ^59^, which we highlight here to encourage its use. In parallel, we followed standardized protocols for sample collection, sample tracking, sample metadata curation, sample shipping, and data release. Importantly, we updated the previous EMP Metadata Guide to accommodate the EMP500 sampling design as well as updates to other standardized ontologies (see Online Methods), including our own application ontology, the Earth Microbiome Project Ontology (EMPO). In addition to the environments previously described by EMPO, EMPO (version 2) now recognizes an important split within host-associated samples representing saline and non-saline environments (Fig. 1a) not detected in the EMP’s previous analysis of 16S rRNA from a separate set of <23,000 samples ^1^.

**Figure 1.**
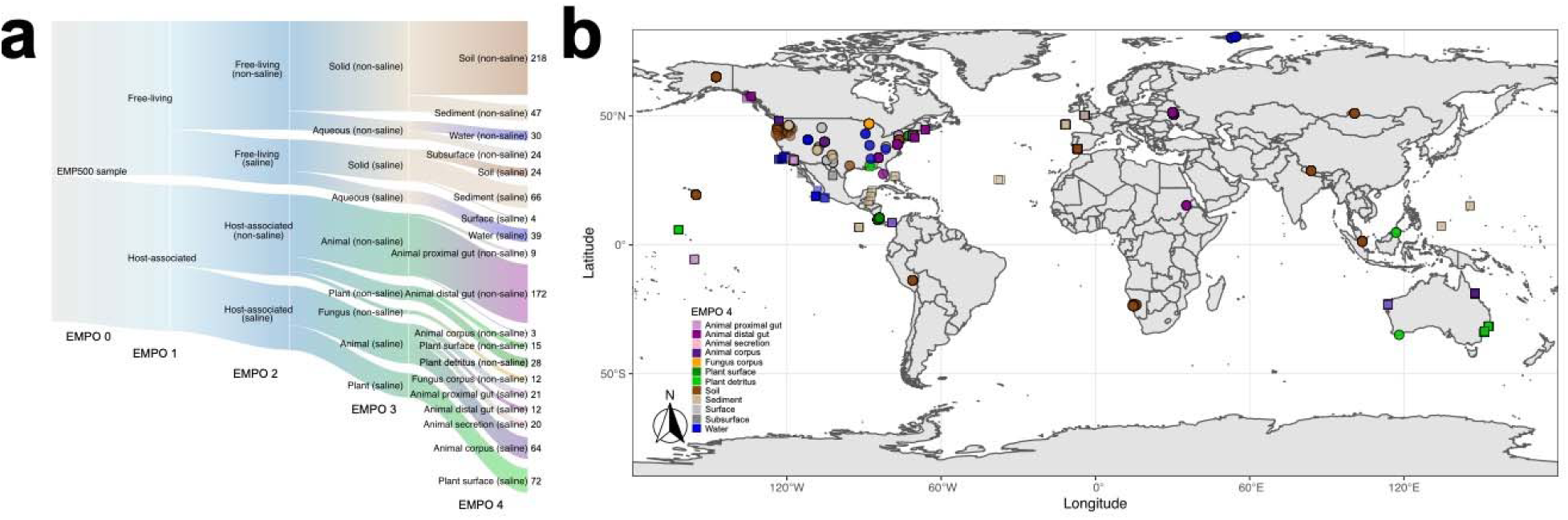
**a**, Distribution of samples (*n* = 880) among the Earth Microbiome Project Ontology (EMPO version 2) categories. This version of EMPO is updated to include the important distinction between saline- and non-saline samples from host-associated environments that was not detected in previous analyses of 16S rRNA alone. **b**, Geographic distribution of samples with points colored by EMPO 4. Points are transparent to highlight cases where multiple samples derive from a single location. We note here that our intent was to sample across environments rather than geography, in part as we previously showed that microbial community composition is more influenced by the former vs. the latter^33^, but also to motivate finer-grained geographic exploration as sample analyses decrease in cost. Extensive information about each sample set is described in Table S1.

For the majority of samples examined here, we successfully generated data for bacterial and archaeal 16S ribosomal RNA (rRNA), eukaryotic 18S rRNA, internal transcribed spacer (ITS) 1 of the fungal ITS region, bacterial full-length rRNA operon, shotgun metagenomics, and untargeted metabolomics (i.e., LC-MS/MS and gas chromatography coupled with mass spectrometry [GC-MS]). A summary of sample representation among each data layer, including read counts for sequence data, is presented in Table S2. To foster exploration of this novel dataset, we have made the raw sequence- and metabolomics data publicly available through Qiita (https://qiita.ucsd.edu; study ID: 13114) ^60^ and GNPS (https://gnps.ucsd.edu; MassIVE IDs: MSV000083475, MSV000083743) ^47^, respectively. We also provide complete protocols for laboratory- and computational workflows for both metagenomics and metabolomics data for use by the broader community, available on GitHub (https://github.com/biocore/emp/blob/master/methods/methods_release2.md). We hope that the dataset and workflows presented here serve as useful tools for others, in addition to providing a framework for launching additional future studies. To provide an example of the utility of the dataset for addressing important questions in microbial community ecology, we present an analysis of microbially-related metabolites and microbe–metabolite co-occurrences across the Earth’s environments (Fig. S1).

### Everything is everywhere, but the environment selects: metabolite intensities reveal habitat-specific distributions

In total, we generated untargeted metabolomics data (i.e., LC-MS/MS) for 618 of 880 samples (Table S2), resulting in 52,496 unique molecular structures, or metabolites, across all samples. We then refined that dataset to include only putative, microbially-related metabolites, resulting in 6,588 metabolites across all samples (12.55% of all metabolites). Focusing on this subset, we found that although the presence/absence of major classes of microbially-related metabolites is indeed relatively conserved across habitats, their relative intensities (i.e., analogous to abundances for microbes) reveal specific chemistry that is lacking or enriched in particular environments, especially when considering more resolved chemical class ontology levels (Fig. 2, Fig. S2).

**Figure 2.**
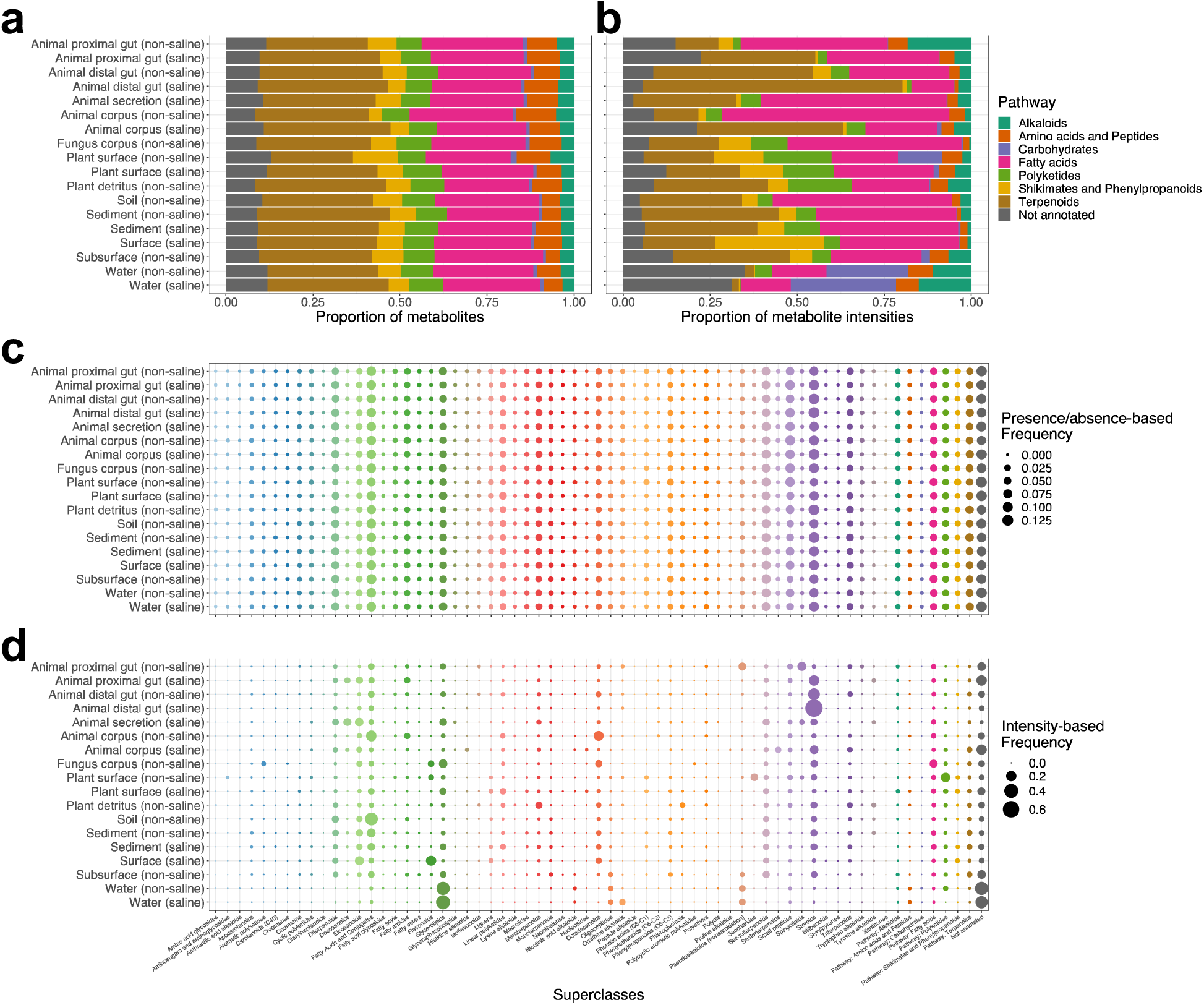
**Distribution of microbially-related secondary metabolite pathways and superclasses among environments** described using the Earth Microbiome Project Ontology (EMPO 4). Individual metabolites are represented by their higher level classifications. Both chemical pathway and chemical superclass annotations are shown based on presence/absence (**a, c**) and relative intensities (**b, d**) of molecular features, respectively. For superclass annotations in panels **c** and **d**, we included pathway annotations for metabolites where superclass annotations were not available, when possible.

Importantly, when considering differences in the relative intensities of all microbially-related metabolites, using methods robust to compositionality ^61^, profiles among environments were so distinct that we could identify particular metabolites whose abundances were significantly enriched in certain environments (Fig. 3a, Table S3). For example, microbially-related metabolites assigned as carbohydrates (i.e., excluding glycosides) were especially enriched in aquatic samples (log fold change [LFC]_Water (non-saline)_ = 0.31±1.22, LFC_Water (saline)_ = 0.54±1.45) (Fig. 3a). Similarly, sediment, marine plant surface, and fungal samples were enriched in polyketides (LFC_Sediment (non-saline)_ = 1.69±0.64, LFC_Sediment (saline)_ = 1.56±1.11, LFC_Plant surface (saline)_ = 1.22 ±0.35, LFC_Fungus corpus (non-saline)_ = 1.68±1.10) and soil, lake sediment, marine plant surface samples were enriched in shikimates and phenylpropanoids (LFC_Sediment (non-saline)_ = 1.90±0.69, LFC_Soil (non-saline)_ = 1.33 ±0.65, LFC_Plant surface (saline)_ = 1.09 ±0.43) (Fig. 3a).

**Figure 3.**
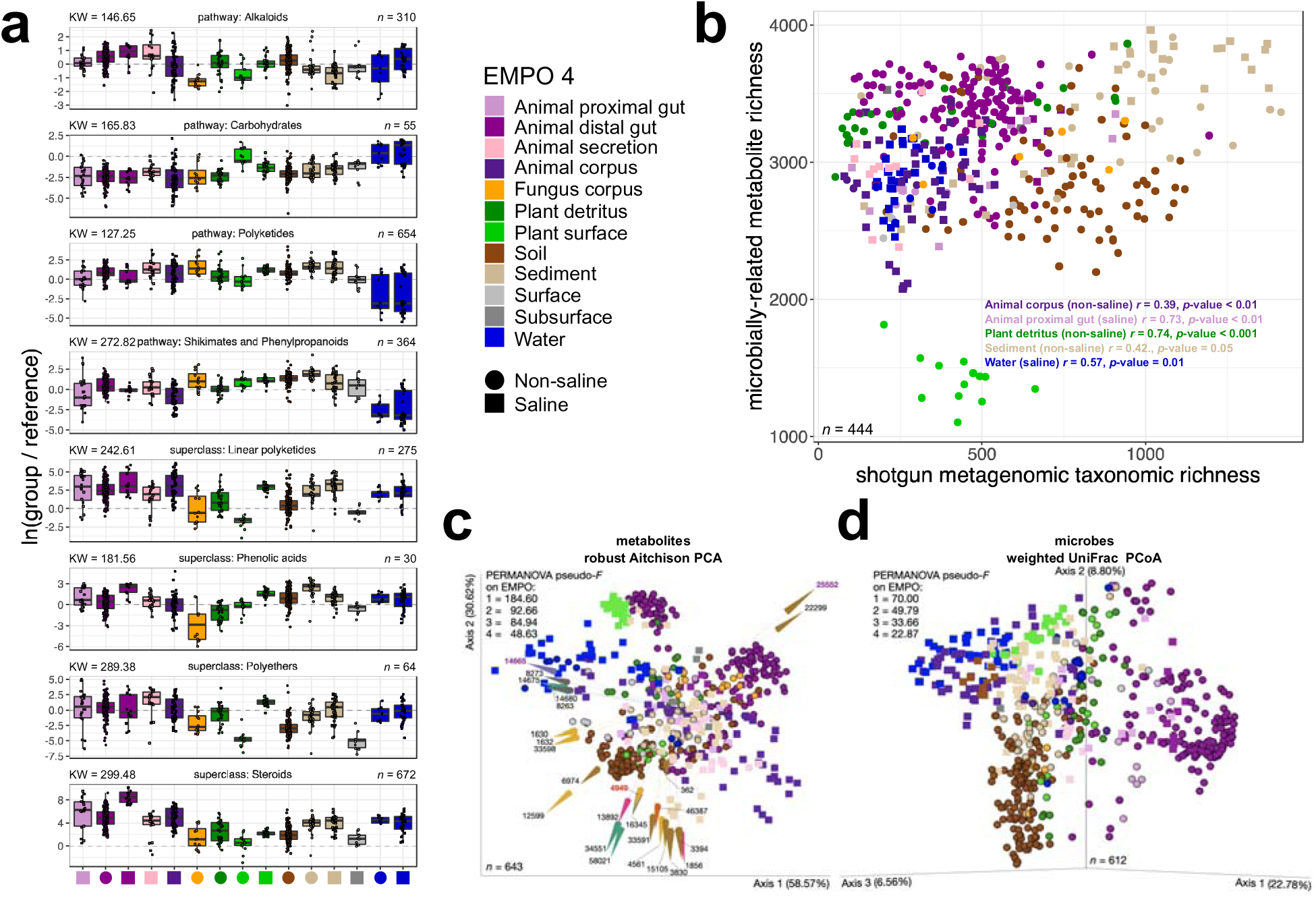
**Structural-level associations between microbially-related secondary metabolites and specific environments** described using the Earth Microbiome Project Ontology (EMPO 4). **a,** Differential abundance of metabolites across environments, highlighting four pathways and four superclasses in separate panels. For each panel, the *y*-axis represents the natural log-ratio of the intensities of metabolites annotated as the listed ingroup divided by the intensities of metabolites annotated as the reference group (i.e., pathway reference: *Amino Acids and Peptides*, *n* = 615; superclass reference: *Flavonoids*, *n* = 42). The number of metabolites in each ingroup is shown, as well as the chi-squared statistic from a Kruskal-Wallis rank sum test for differences in log-ratios across environments (i.e., each test had *p*-value < 2.2 x 10^-16^). Each test included 606 samples. Boxplots are in the style of Tukey, where the center line indicates the median, lower and upper hinges the first- and third quartiles, respectively, and each whisker 1.5 x the interquartile range (IQR) from its respective hinge. Outliers from boxplots are colored red to highlight that they are also represented in the overlaid, jittered points. **b,** Relationship between metabolite richness and microbial taxon richness across samples and environments, with significant correlations noted. **c,** Turnover in composition of metabolites across environments, visualized using Robust Aitchison Principal Components Analysis (RPCA), showing samples separated based on LC-MS/MS metabolite abundances. Shapes represent samples and are colored and shaped by EMPO. Arrows represent metabolites, and are colored by chemical pathway. The direction and magnitude of each arrow corresponds to the strength of the correlation between the relative abundance (i.e., intensity) of the metabolite it represents and the ordination axes. Samples close to arrow heads have strong, positive associations with respective metabolites, whereas samples at and beyond arrow origins have strong, negative associations. The 25 most important metabolites are shown and are described in Table S4. Features annotated in red/purple were among the top ranked metabolites based on log fold changes with respect each environment (Tables S3), and those in purple were additionally identified as important in co-occurrence analyses (Fig. 4). **d,** Turnover in composition of microbial taxa across environments, visualized using Principal Coordinates Analysis (PCoA) of weighted UniFrac distances. Results from PERMANOVA (999 permutations) for each level of EMPO are shown (all tests had *p*-value = 0.001; group sizes for LC-MS/MS data: *k*_EMPO1_ = 2, *k*_EMPO2_ = 4, *k*_EMPO3_ = 9, *k*_EMPO4_ = 18; group sizes for shotgun metagenomics data: *k*_EMPO1_ = 2, *k*_EMPO2_ = 4, *k*_EMPO3_ = 9, *k*_EMPO4_ = 19). Sample sizes in panel **a** refer to metabolites, but in all other panels refer to samples.

We also observed differences in the total number of distinct microbially-related metabolites (i.e., richness), which varied strongly across environments (Fig. 3b). We note that whereas sediment samples were most rich, with sediments from saline environments exhibiting relatively greater metabolite diversity than those from non-saline environments, the surfaces of terrestrial plants were especially lacking in metabolite diversity (Fig. 3b). This was in contrast to metabolite diversity in detritus of terrestrial plants, which was also high (Fig. 3b).

When considering the identity and relative intensity of each metabolite in analysis of beta-diversity, we observed a separation of samples based in part on host-association and salinity (PERMANOVA for EMPO 2: pseudo-*F* = 92.66, *p*-value = 0.001), with specific environments clustering in ordination space (PERMANOVA for EMPO 4: pseudo-*F* = 48.63, *p*-value = 0.001) and certain metabolite features identified as important in separating all samples (Fig. 3c, Table S4). For the latter, we identified three metabolites also identified among the top ten most differentially abundant metabolites for each environment (Table S3): one chalcone associated with the surfaces of terrestrial plants (C_13_H10O, ID: 4949), one glycerolipid associated with freshwater (C_28_H_58_O_15_ ID: 14665), and one cholane steroid associated with the distal guts of terrestrial animals (C_24_H_34_O_2_, ID: 25552) (Fig. 3c). Somewhat unexpectedly, we observed the majority of saline environments to cluster as a third group between two distinct groups of non-saline ones, separating the soil from the animal distal gut samples (PERMANOVA for salinity: pseudo-*F* = 8.25, *p*-value = 0.001) (Fig. 3c). This is unique from what has been reported previously for samples based on microbes, which largely form two distinct clusters representing saline and non-saline environments ^1, 5^. Similarly, whereas the split in salinity was strong for host-associated environments, it was less so for free-living ones, with both water and sediment samples separately clustered regardless of salinity (Fig. 3c). With these observations, we predicted that the differences in the relative intensities of particular metabolites among environments were due in part to underlying differences in microbial community composition and diversity. To explore these relationships, we additionally explored our shotgun metagenomics data.

### Metabolite and microbial alpha-diversity have strong positive and environment-specific relationships

We found significant, positive correlations between microbially-related metabolite richness and microbial taxon richness across all samples (*r* = 0.20, *p*-value < 0.001), within host-associated samples (*r* = 0.19, *p*-value < 0.01), within free-living samples (*r* = 0.18, *p*-value < 0.05), and for certain environments: Animal proximal gut (saline) (*r* = 0.73, *p*-value < 0.01), Plant detritus (non-saline) (*r* = 0.74, *p*-value < 0.001), Sediment (non-saline) (*r* = 0.42., *p*-value = 0.05), and Water (saline) (*r* = 0.57, *p*-value = 0.01) (Fig. 3b, Table S5). We observed non-significant trends in correlations for Plant surface (non-saline) (*r* = -0.36, *p*-value = 0.2) and Sediment (saline) (*r* = 0.27, *p*-value = 0.1) (Fig. 3b, Table S5). Relationships for other environments were weaker (Fig. 3b, Table S5). Sediment samples had the highest alpha-diversity of both microbial taxa and metabolites (Fig. 3b). Correlations with metabolite richness were weaker when using Faith’s Phylogenetic Diversity (PD) and weighted Faith’s PD for quantifying microbial alpha-diversity (Table S5).

### Turnover and nestedness of metabolite and microbial taxon profiles is related to the environment

When considering the identity and relative abundance of features (i.e., intensity for metabolites), we found similarity in the clustering of samples by environment between microbially-related metabolite and microbial taxon datasets (Fig. 3c,d). We also observed a strong correlation between sample–sample distances based on metabolites vs. microbial taxa (Table 1). With the exception of certain animal-associated samples (e.g., Animal corpus [non-saline], Animal proximal gut [non-saline]), the distribution of environments in ordination space was nearly identical between datasets (Fig. 3c,d). For example, the separation of free-living (e.g., Water) and host-associated (e.g., Animal distal gut) environments along Axis 1, and a gradient from living hosts (e.g., Plant surface), to dead organic material (e.g., Plant detritus), to soils and sediments along Axis 2, was clear in both datasets (Fig. 3c,d). When focusing on the separation of samples within a single environment such as soil, we observed much more variability between metabolite and microbial taxon datasets (Mantel *r* = 0.32 for soil vs. 0.43 for all environments, *p*-value = 0.001 for both tests). This highlights not only novelty among soil samples from distinct geographic locations (Fig. S3), but also the insight that can be gained from using a multi-habitat dataset. To assess whether metabolite profiles were more similar to those for microbial taxa vs. microbial functions, we annotated our metagenomic reads to profile enzymes. We found the separation of samples based on microbial functions to be unique and largely driven by animal gut samples as compared to when based on either metabolites or microbial taxa (Fig. S4). However, correlations in sample–sample distances between microbial functional data and other datasets were relatively strong (Table 1).

**Table 1.**
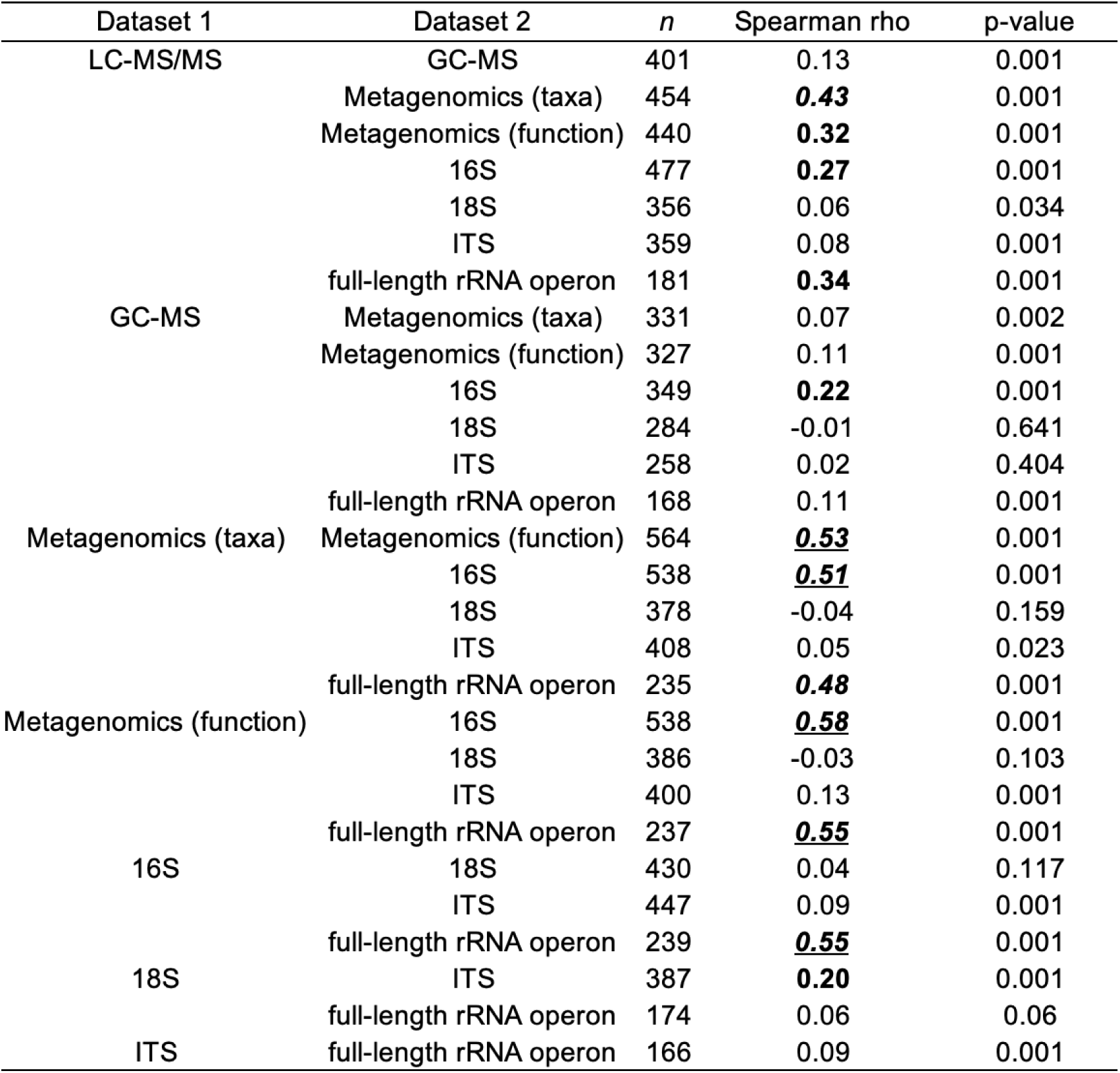
Mantel test results comparing data layers generated for the EMP500 samples. Note the strong relationships between the metabolomics data (i.e., LC-MS/MS and GC-MS) and the sequence data from Bacteria and Archaea (i.e., shotgun metagenomics, 16S, and full-length rRNA operon)as compared to between metabolomics data and sequence data from eukaryotes (i.e., 18S and ITS). There are also strong relationships between difference sequence data from Bacteria and Archaea (rho > 0.2 in bolded font; > 0.4 in bolded, italics; > 0.5 additionally underlined).

Interestingly, we observed a more consistent split between saline and non-saline samples when based on microbial taxa as compared to the metabolites (PERMANOVA on salinity: pseudo-*F* = 40.94 for microbes vs. 8.25 for metabolites, *p*-value = 0.001 for both tests) (Fig. 3c,d). Conversely, for each level of EMPO, the effect size for explaining variation in the separation of samples based on metabolites was roughly twice that for explaining it based on microbes (Fig. 3c,d). The effect sizes for EMPO when separating samples based on microbial functions (i.e., enzymes) were even stronger, however as noted above only animal gut samples clustered separately from all other environments along the major axis of variation when using dimensionality reduction (Fig. S4).

In the absence of complete turnover in microbially-related metabolites and microbial taxa across environments, apparent by the overlap of clusters of samples from different habitats in our ordinations (Fig. 3c,d), we quantified the extent of nestedness among all samples. Nestedness describes the degree to which features in one environment are nested subsets of another environment, and can provide insight into community assembly dynamics ^1, 62^. We found that samples were significantly nested based on both metabolites (Fig. S5) and microbial taxa (Fig. S6), and that certain environments were consistently nested within others, however this pattern varied between datasets. For example, based on microbial taxa we observed host-associated samples to be nested within free-living ones (Fig. S6a), however the opposite was true for metabolites, although the pattern was weaker (Fig. S5a). When considering host-association and salinity (i.e., EMPO 2) for metabolites, free-living samples were more nested than host-associated ones, and within each group non-saline samples were more nested than saline ones (Fig. S5d). This pattern remained consistent when describing metabolites at the superclass, class, and molecular formula levels (Fig. S5d). Patterns of nestedness were less consistent across taxonomic levels when based on microbial taxa, although non-saline, free-living samples were the most nested across the family, genus, and species levels (Fig. S6d). When considering all environments together (i.e., for EMPO 3 and 4), we observed stronger patterns of nestedness among environments for microbial taxa (Fig. S5b,c) vs. metabolites (Fig. S6b,c). Focusing on gradients of host-association and salinity, we observed patterns of nestedness were more similar between microbial taxa and metabolites for host-associated environments (Fig. S5e, Fig. S6e). However, there was strong disagreement in the nestedness ranks of plant surfaces among all non-saline, host-associated samples based on microbial taxa (Fig. S5e) vs. metabolites (Fig. S6e).

### Certain metabolites and microbes can be used to distinguish among habitats

Based on the strong relationships among metabolites, microbes, and the environment, we next tested the hypothesis that specific metabolites, microbial taxa, or microbial functional products (i.e., enzymes) could be used to classify samples among environments. Using a machine-learning classifier (see Online Methods), we were able to identify specific metabolites that could classify samples among environments with 88.0% overall accuracy (Fig. 4a, Fig. 5a, Fig. S7a, Fig. S8, Table S6). After ranking all microbially-related metabolites based on their impact in distinguishing environments, we found the top ranked metabolites to include a diterpenoid negatively associated with non-saline soils (C_20_H_32_, ID: 04492), an undescribed metabolite positively associated with marine sediments (ID: 42202), a lignan negatively associated with freshwater sediments (C_20_H_20_O_5_, ID: 07899), a diterpenoid negatively associated with the surfaces of terrestrial plants (C_20_H_28_O_3_, ID: 07719), and an undescribed metabolite positively associated with non-saline subsurfaces (ID: 14598) (Fig. 5a, Table S6). Among the top twenty ranked metabolites with annotations, the majority were alkaloids, fatty acids, or terpenoids, with terpenoids being most impactful among the top ten ranked metabolites, including the most highly ranked one (Fig. 5a, Table S6).

**Figure 4.**
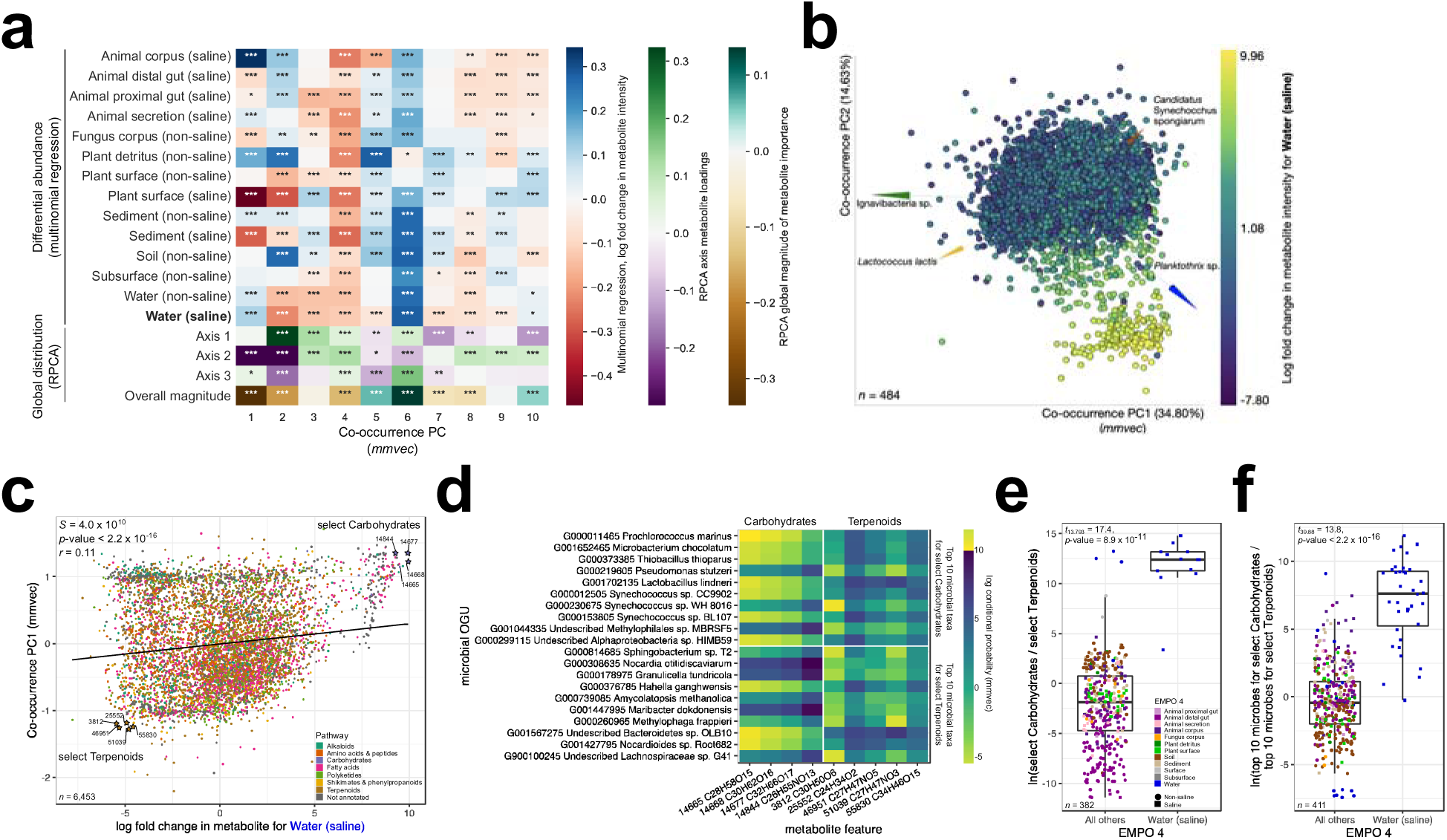
**Metabolite-microbe co-occurrences vary strongly across environments. a,** Co-occurrence analysis results showing correlation between the metabolite loadings from the co-occurrence ordination (i.e., co-occurrence principal coordinates [PCs]) and (i) log fold changes in metabolite abundances across environments, (ii) metabolite loadings from the ordination in Fig. 3d corresponding to clustering of samples by environment based on metabolite profiles (i.e., Global distribution, Axes 1-3), and (iii) a vector representing the overall magnitude of microbial taxon abundances from the same ordination (i.e., Global distribution, Overall magnitude). Values are Spearman correlation coefficients. Asterisks indicate significant correlations (\**p* < 0.05, \*\**p* < 0.01, \*\*\**p* < 0.001). **b,** The relationship between log fold changes in metabolite abundance with respect to ‘Water (non-saline)’ and the first three PCs of the co-occurrence ordination shown as a multi-omics biplot of metabolite–microbe co-occurrences. Points represent metabolites, and the distance between metabolites indicates similarity in their co-occurrences with microbial taxa (i.e., two metabolites that are close together have similar co-occurrence probabilities to the same microbes). Metabolites are colored based on their relative log fold changes with respect to ‘Water (non-saline)’. Vectors represent specific microbial taxa, where distances between arrow tips indicate similarity in their co-occurrence with specific metabolites (i.e., two microbes that are close together have similar co-occurrence probabilities to the same metabolites), and the direction of each arrow indicates which metabolites each microbe co-occurs most strongly with. **c,** The relationship between log fold changes in metabolite abundances with respect to ‘Water (non-saline)’ and loadings for metabolites on PC1 of the co-occurrence ordination. The correlation is one example of those summarized in panel **a**. Metabolites are colored by pathway. Select Carbohydrates (excluding glycosides) representing the focal group and select Terpenoids representing reference group are highlighted. **d,** The top 10 co-occurring microbial taxa for all select Carbohydrates and all select Terpenoids, with a heatmap showing the co-occurrence strength between each metabolite-microbe pair. **e,** Log-ratio of metabolite intensities for select Carbohydrates and select Terpenoids. **f,** Log-ratio of abundances of the top ten microbial taxa associated with select Carbohydrates and the top ten microbial taxa associated with Terpenoids. For panels **e** and **f** samples are colored by environment (based on EMPO 2), and results from a *t*-test comparing ‘Water (saline)’ vs. all other environments are shown. Boxplots are in the style of Tukey, where the center line indicates the median, lower and upper hinges the first- and third quartiles, respectively, and each whisker 1.5 x the interquartile range (IQR) from its respective hinge.

**Figure 5.**
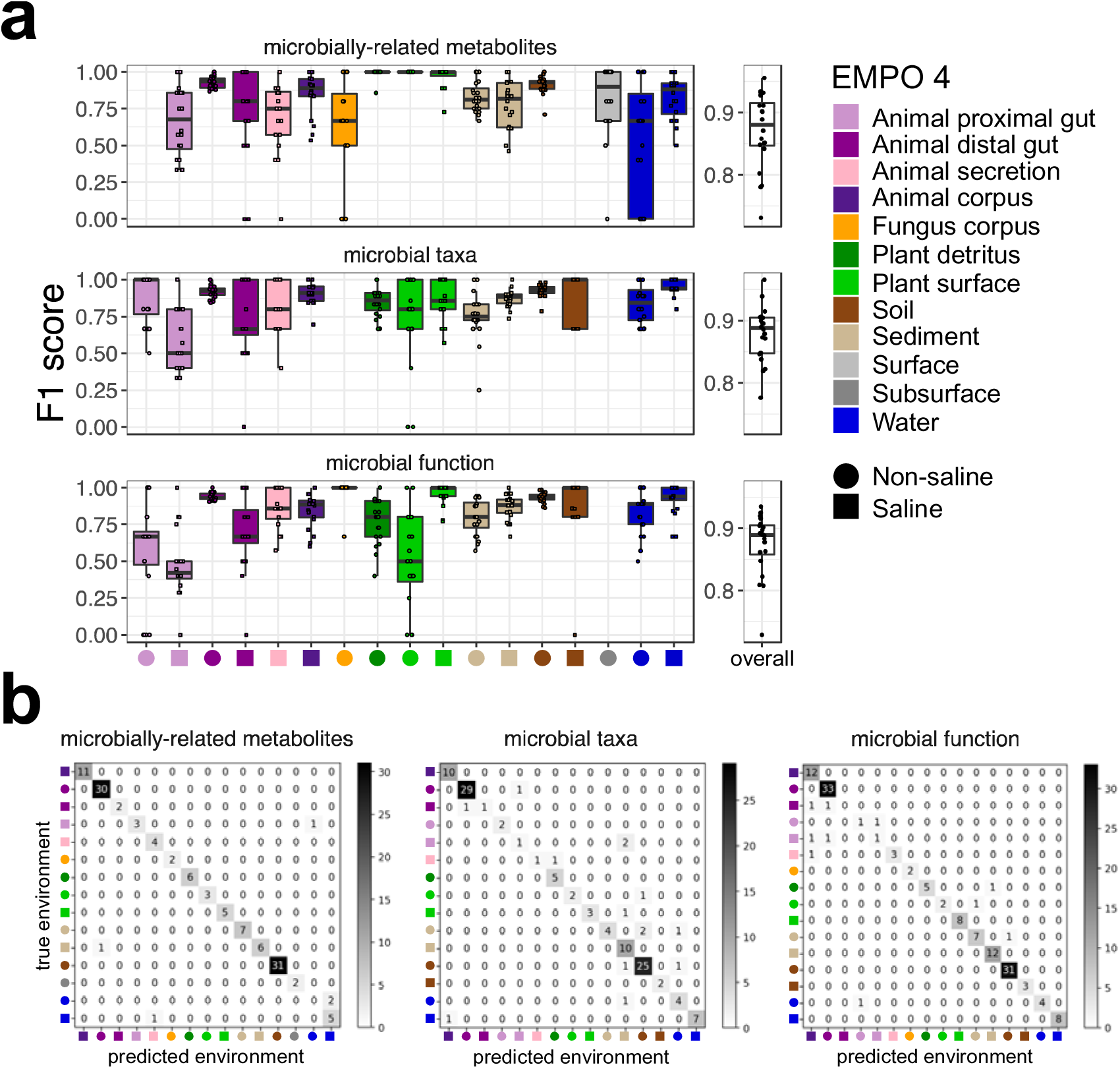
**Machine-learning analysis of microbially-related metabolites, microbial taxa, and microbial functions**, **highlighting per-environment classification performance. a,** The F1 score (i.e., which considers precision and recall) for each environment as well as overall across all environments. **b,** Confusion matrices for each data layer highlighting which pairs of environments are confused. Boxplots are in the style of Tukey, where the center line indicates the median, lower and upper hinges the first- and third quartiles, respectively, and each whisker 1.5 x the interquartile range (IQR) from its respective hinge. For all analyses, environments are described by the Earth Microbiome Project Ontology (EMPO 4).

We also found strong support among methods for the importance of particular metabolites in distinguishing environments. For example, the undescribed metabolite positively associated with marine sediments (i.e., ID: 42202) and one fatty acid – a monoacylglycerol (i.e., ID: 42202) – revealed as useful in classification in this analysis also stood out in our analysis of differential abundance (Fig. 5a, Table S3, Table S6). Similarly, distinct analytical approaches identified specific metabolites as particularly important for distinguishing aquatic samples (i.e., one glycerolipid, C_28_H_58_O_15_, ID: 14665 and one pseudoalkaloid, C_18_H_22_N_7_O_5_, ID: 14675), non-saline plant surface samples (i.e., one chalcone, C_13_H_10_O, ID: 4949), and non-saline animal distal gut samples (i.e., one cholane steroid, C_24_H_38_O_4_, ID: 2552 and one prenyl quinone monoterpenoid, C_29_H_46_O_2_, ID: 22299) (Fig. 3c, Table S3, Table S4). We explored these relationships further in our multi-omics analyses.

Using the same machine learning approach on our metagenomic sequence data, we identified specific microbial taxa and microbial functional products (i.e., enzymes) useful in classifying samples to environments, with 88.8% and 88.9% overall accuracy, respectively (Fig 4, Fig. 5, Fig. S7, Fig. S9, Fig. S10). Regarding microbial taxa, we observed the majority of the top twenty ranked taxa with respect to classification performance were Proteobacteria (Fig. 5b). The Cyanobacteria, Firmicutes, and Actinobacteria were represented by a few members each, and *Candidatus* Tectomicrobia and Euryarchaeota represented as singletons (Fig. 5b). The most highly ranked taxon, *Conexibacter woesei* (G000424625, Actinobacteria), was positively associated with non-saline soils, and is an early-diverging member of the class *Actinobacteria* first isolated from temperate forest soil in Italy ^63^ (Fig. 5b). Also among the top ranked taxa were *Haloquadratum walsbyi* (Euryarchaeota) positively associated with saline soils, *Pantoea dispersa* (*Gammaproteobacteria*) and an undescribed species of *Bacillus* (Firmicutes) positively associated with the detritus of terrestrial plants, *Serratia fonticola* (*Gammaproteobacteria*) positively associated with the surfaces of terrestrial plants, and *Oenococcus oeni* (Firmicutes) positively associated with marine animal secretions (Fig. 5b). *Roseovarius nubinhibens* (*Alphaproteobacteria*), *Ca.* Synecochoccus spongiarum (Cyanobacteria), and *Paraclostridium bifermentans* (Firmicutes) were also highly ranked (Fig. 5b). With respect to microbial functions, we note that the majority of the top twenty most highly ranked enzymes with respect to classification performance were oxidoreductases or transferases, followed by hydrolases, and then isomerases and lyases (Fig. 5c). The most highly ranked enzyme was positively associated with non-saline soils and was a trehalohydrase (EC: 3.2.1.141), an enzyme that binds trehalose, a carbon-source commonly produced by soil-inhabitants including plants, invertebrates, bacteria, and fungi, with potential roles in symbioses ^64^. Also among the most highly ranked enzymes were a glutamate carboxylase (EC: 4.1.1.90) positively associated with the surfaces of marine plants, a linoleate lipoxygenase (EC: 1.13.11.60) positively associated with lichen thalli, and a glyceraldehyde dehydrogenase (EC: 1.2.1.59) positively associated with saline soils (Fig. 5c). A glucosyltransferase (EC: 2.4.1.256), a uroporphyrinogen decarboxylase (EC: 4.1.1.37), and a hydroxylamine dehydrogenase (EC: 1.7.2.6) were also highly ranked (Fig. 5c).

### Multi-omics co-occurrence analysis reveals strong relationships between specific metabolites, microbes, and the environment

In addition to exploring relationships between metabolite and microbial diversity, we sought to explicitly quantify metabolite–microbe co-occurrence patterns. In particular, we examined associations between metabolites and the environment (e.g., Fig. 3a,c), while also considering each metabolite’s co-occurrence with all microbes in the dataset (Fig. S1). In that regard, we first generated metabolite–microbe co-occurrences learned from both LC-MS/MS metabolomics- and shotgun metagenomic profiles across all samples, for a cross-section of 6,501 microbially-related metabolites and 4,120 microbial taxa (Fig. S11, Fig. S12). Whereas most metabolites co-occurred with at least a few microbes, few were found to co-occur with many microbes (Fig. S11a). The distribution of co-occurrences was not heavily shifted towards any particular pathway (Fig. S11b), however certain superclasses exhibited co-occurrences with many microbes, including diarylheptanoids and phenylethanoids (C6-C3) (Fig. S11c). Similarly for microbes, co-occurrences with metabolites were not heavily skewed towards particular phyla, although specific clades were enriched, such as the most recently diverged members of the Bacteroidetes (Fig. S12). In contrast to their co-occurrences with metabolites, log fold changes in microbial abundances with respect to the environment appear to be phylogenetically conserved, and correlated with salinity and association with the animal gut environment (Fig. S12).

Next, using metabolite–metabolite distances based on co-occurrence profiles considering all microbes, we ordinated metabolites in microbe space. We then examined correlations between metabolite loadings from the first ten principal coordinates of that co-occurrence ordination and (1) log-fold changes of metabolites across environments (e.g., Fig. 3a), and (2) distributions of metabolites across all samples (i.e., loadings and overall magnitude from ordination of all samples) (Fig. 3c), and found strong relationships with each (Fig. 6a). In particular, the abundances of microbially-related metabolites in plant surface (saline), sediment (saline), and aquatic samples (i.e., those from water) had strong correlations with microbe– metabolite co-occurrences (Fig. 6a). Focusing on seawater (i.e., Water [saline]), we visualized the correlation between metabolite loadings on PC1 of the co-occurrence ordination, which represent differences based on co-occurrences with microbes (Fig. 6b), and log fold changes in metabolite abundances with respect to seawater (Fig. 6c). In this space, features with high values for both vectors should be associated with the same microbes and also highly abundant in the ocean, whereas features with low values for both vectors should be associated with the same microbes and have low-to-zero abundance in the ocean (Fig. 6c). Focusing on one group of carbohydrates (excluding glycosides) and one group of terpenoids (Fig. 6c,d), we found significant differences in their intensities in seawater vs. all other environments (Fig. 6e), as well in the abundances of their top co-occurring microbial taxa (Fig. 6f). Importantly, by relying on our metabolite intensity data, this result validates patterns identified in our analyses of differential abundance across environments and co-occurrence with microbial taxa. We used this same approach to explore metabolite–microbe co-occurrences specific to other environments (Fig. S13), further revealing strong turnover in metabolite–microbe co-occurrences across habitats. Visualizing the differential abundance of metabolites with respect to seawater and other environments in the broader context of metabolite–microbe co-occurrences highlights the especially unique community profiles across habitats (Fig. 6, Fig. S13). Importantly, these results demonstrate that metabolites and microbes can be used to classify- and co-occur among environments.

**Figure 6.**
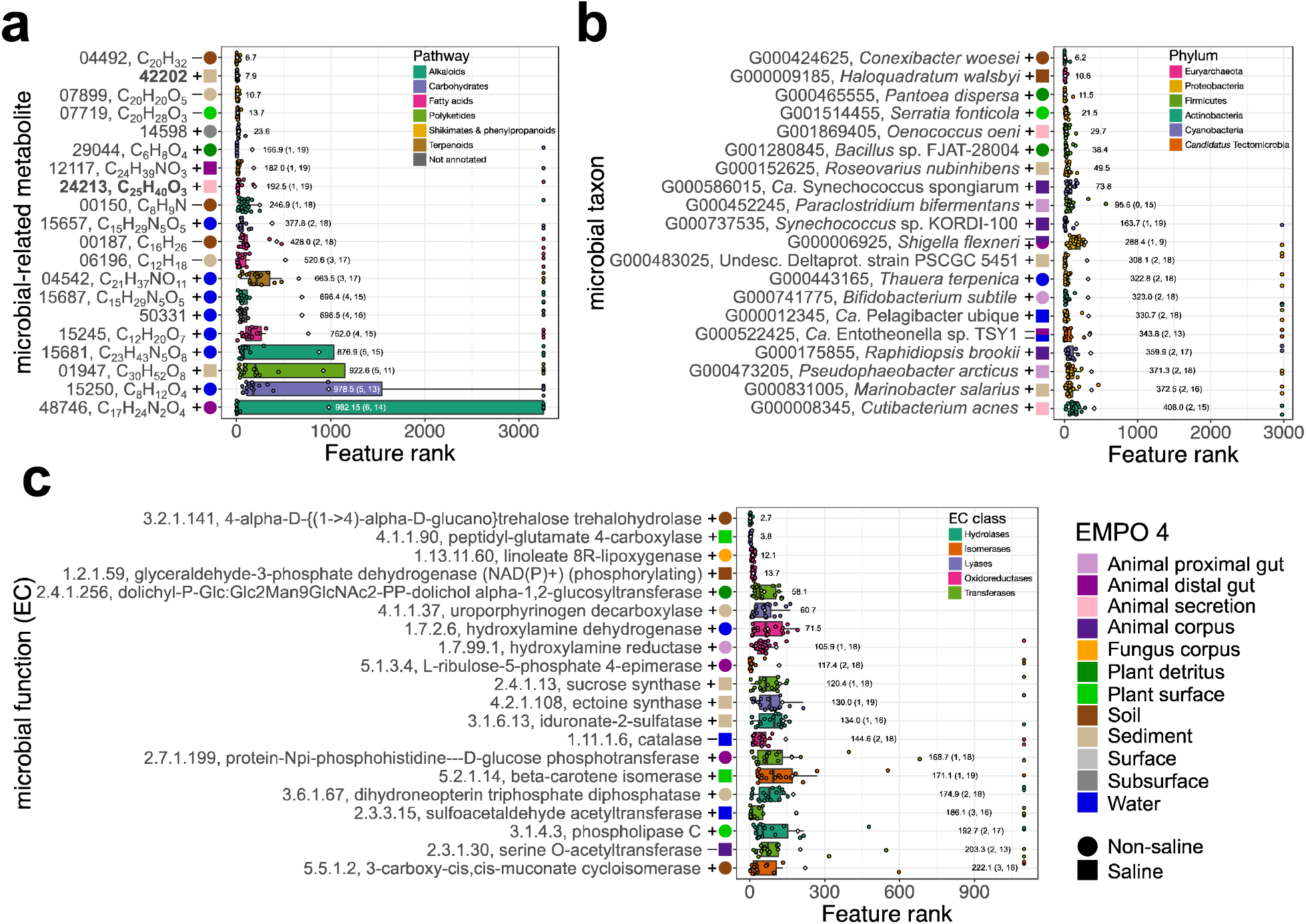
**Machine-learning analysis of microbially-related metabolites, microbial taxa, and microbial functions, highlighting the top twenty most impactful features for each dataset. a,** The top twenty most impactful microbially-related metabolites. Features are colored by metabolite pathway. Metabolites in bold font are those also identified as important in analysis of differential abundance analysis (Table S3). **b,** The top twenty most impactful microbial taxa (i.e., OGUs). Taxa are colored by phylum. **c,** The top twenty most impactful microbial functions (i.e., KEGG ECs). Boxplots are in the style of Tukey, where the center line indicates the median, lower and upper hinges the first- and third quartiles, respectively, and each whisker 1.5 x the interquartile range (IQR) from its respective hinge. Enzymes are colored by class. For all features, ranks are based on impacts derived from SHAP values. Associations with environments are indicated, where + indicates a positive association and – indicates a negative association based on feature abundances. Diamonds and values to the right of boxes indicate means. Values in parentheses indicate (1) the number of iterations (*n* = 20) in which a feature had no impact, and (2) the number of iterations in which the reported association was observed, for cases in which values were < 20. Environments are described by the Earth Microbiome Project Ontology (EMPO 4).

### Correlations with amplicon sequence data and GC-MS data

Harnessing additional data generated for EMP500 samples, including GC-MS and amplicon sequence data (i.e., bacterial and archaeal 16S and full-length rRNA operon, eukaryotic 18S, fungal ITS), we compared sample–sample distances (i.e., beta-diversity) between each pair of datasets. Importantly, we found further support for a strong relationship between microbially-related metabolites and microbial taxa (*r* = 0.43, *p*-value = 0.001) (Table 1). The relationships between the metabolomics data (i.e., LC-MS/MS or GC-MS) and sequence data from eukaryotes (i.e., 18S or ITS) were weaker (e.g., LC-MS/MS vs. ITS; *r* = 0.08, *p*-value = 0.001) (Table 1). The weakest relationships were between the GC-MS metabolomics profiles and those from sequence data from eukaryotic (i.e., 18S or ITS) (e.g., GC-MS vs. ITS; *r* = 0.02, *p*-value = 0.4) (Table 1). The strongest relationships were between different layers of sequence data from bacteria and archaea (Table 1). For example, correlations between 16S rRNA profiles and those from full-length rRNA operons had *r* = 0.55 (*p*-value = 0.001), and 16S vs. shotgun metagenomics had *r* = 0.51 (*p*-value = 0.001) (Table 1). These results highlight the strong relationship between metabolic profiles and microbial taxonomic composition across habitats spanning the globe.

## DISCUSSION

Here, we produced as a resource a novel, multi-omics dataset comprising 880 samples that span 19 major environments, contributed by 34 principal investigators for the EMP500 (Fig. 1, Table S1). Whereas we updated EMPO to include these environments, we recognize that certain environments are represented here by only a handful of samples (Fig. 1) and/or a single sample set (Table S1), and note that we had to exclude them from some of our analyses due to low representation (e.g., machine learning and co-occurrence analyses). In that regard, we recommend that future similar efforts focus on additional sampling of these environments in order to further generalize our findings to those habitats. Similarly, we hope to expand sampling geographically to broaden our scope of inference, as many important environments and locations could not be included here (or, indeed, in the EMP’s 27,000-sample dataset^1^), which makes exploring certain questions difficult. To foster this research activity, we expanded upon the widely-adopted set of the EMP’s standardized protocols for guiding microbiome research – from sample collection to data release ^59^ – with new protocols for performing untargeted metabolomics and shotgun metagenomics across a diversity of sample types (Fig. 1a; Online Methods).

Across all 880 samples, we generated eight layers of data, including untargeted metabolomics and shotgun metagenomics (Table S2), providing a valuable resource for both multi-omics and meta-analyses of microbiome data. As expected, sample dropout in any one data layer was non-negligible (Table S2), reducing the number of samples in any one layer to at most roughly 500 samples (hence the EMP500). For future similar studies, we note that considering multi-omics applications during the experimental design and/or sample collection phases of studies is crucial, in part because certain metabolomics approaches are not amenable to samples stored in particular storage solutions commonly used for metagenomics (e.g., RNAlater).

We also included an example of how to apply this dataset towards addressing important questions in microbial ecological research, by describing the Earth’s microbial metabolome using an integrated ‘omics approach (Fig. S1). We first explored whether every metabolite is everywhere, but the environment selects (i.e., the Baas Becking hypothesis^51, 52^, but for microbially-related metabolites). Our results confirm that all major groups (e.g., pathways) of metabolites are present in each environment ^14^, but additionally demonstrate that their relative abundances (i.e., intensity) can be limited- or enriched across environments (Fig. 2, Fig. 3a,c, Table S3). Considering the relative intensities of secondary metabolites vs. presence/absence alone drastically strengthens differences in metabolite profiles across environments for many metabolite groups including apocarotenoids, fatty acids and conjugates, glycerolipids, steroids, and polyketides (Fig. 2c,d, Fig. 3a). Interestingly, the groups of metabolites that exhibited the most obvious changes in representation were those that appeared in the fewest samples (i.e., those at low prevalence from a presence/absence perspective; e.g., carbohydrates [excluding glycosides], alkaloids) (Fig. 2a,b). Further, the environments with the most unannotated metabolites include terrestrial animal cadavers, bioreactors that mimic the rumen of cows, freshwater, and the ocean (Fig. 2b). Similarly, environments with the most unannotated shotgun metagenomic reads for microbial functions (i.e., enzymes) included animal cadavers, marine animal secretions, terrestrial plants, and fungi (Fig. S14f). These environments merit further attention with respect to feature description, and represent valuable opportunities for the discovery of novel metabolites and functional products. Whereas we interpret these results as strong evidence that every metabolite is everywhere, but the environment selects, as our study was not designed to address this hypothesis explicitly, further evidence is needed. For example, features at abundances below the detection limit of our approach could not be considered here, but may alter our view of these patterns.

Next, we explored whether the richness and composition of metabolites in any given sample reflect those of co-occurring microbial communities. We compared alpha-diversity between metabolites and microbes, and found strong positive relationships across all samples and for many environments (Fig. 3b, Table S5). As unannotated features are included in these metrics, reference database coverage does not influence the results. Similarly, whereas estimates may be influenced by our use of rarefaction to normalize sampling effort, estimating diversity in absence of such an approach has been shown to be problematic ^65^. Further, we avoided marrying estimates of alpha-diversity from our 16S vs. metagenomic data, as taxonomic profiling used a distinct reference database curated specifically for its respective data type. Future similar studies should consider the development of a reference combining both 16S and shotgun metagenomic data from bacteria and archaea. Similarly, the absence of a relationship between metabolite and microbial richness for other environments may be due to low sample representation as two lowly sampled environments, marine plant surfaces and sediments, both exhibited trends (Table S5) . In absence of technical variation, unique community carrying capacities for metabolite vs. microbes across environments may also skew trends, and we recognize that certain environments simply may not exhibit clear or underlying relationships. Still, when ranking environments based on alpha-diversity, certain patterns are clear. For example, in addition to confirming previous observations that marine sediments are one of the most microbially diverse environments on Earth ^1^, we showed that marine sediments are also the most metabolically diverse (Fig. 3b). To our knowledge, this is the first assessment of the metabolic alpha-diversity in marine sediments as it relates to microbial diversity in those samples in a context including other diverse environments. Although not the most lacking with respect to annotation rates for metabolites, the high diversity in marine sediments merits further exploration of those molecules.

We also found a strong correlation in sample–sample distances between metabolite and microbial datasets (Table 1), and significant turnover of features across environments (Fig. 3c,d). For both metabolites and microbial taxa, the effect of host-association was much stronger than salinity in explaining variation in community composition (Fig. 3c,d). We also observed a much weaker influence of salinity in separating samples based on metabolites vs. microbes (Fig. 3c,d). Together, these findings support recognizing host-association as EMPO 1, and confirm our prediction that environmental similarity between datasets may be distinct when based on one dataset vs. the other. This may indicate that although microbial cells respond strongly to salinity gradients, the taxa they represent can have similar metabolic profiles. Our hypothesis is supported by our observation that co-occurrences with metabolites appear to be more structured by environment than phylogeny for microbial taxa (Fig. S12). Additional support lies in our finding that the differential intensities of important pathways and superclasses are highly similar between freshwater and marine environments (Fig. 3a), and that the same groups of metabolites can occur in both habitats yet are associated with distinct microbes within each (Fig. 6e,f).

The lack of complete turnover in metabolites vs. microbes with respect to the environment generated unique patterns of nestedness between datasets (Fig. S5, Fig. S6). Whereas nestedness patterns among environments with respect to microbial taxon profiles matched our expectations based on assembly dynamics and dispersal patterns (e.g., host-associated communities are a subset of free-living ones), as well as previous observations based on 16S data ^1^, those based on metabolite profiles were more weakly correlated with our description of environments based on EMPO. This may be in part due to the weaker effect of salinity on sample beta-diversity for metabolites (Fig. S5a, Fig. S6a), and similarity in metabolite profiles among microbes from disparate environments (Fig. 6d-f). It may also indicate that microbially-related metabolites assembly and structure uniquely from the microbes that produce them in nature. Nevertheless, future efforts to expand database coverage for metabolites should consider this, as the expectation that diversity will continue to increase with sampling of these distinct environments may not be realized.

Given that profiles for metabolites and microbes were habitat-specific, we used machine-learning to identify several metabolites, microbial taxa, and microbial functions that could accurately classify samples among environments (Fig. 4, Fig. 5, Table S6). Overall accuracy for each dataset was ≥88% (Fig. 4, Fig. S7), confirming our prediction that certain features could distinguish among environments. Although infrequent among 20 iterations, certain environments were occasionally confused (Fig. 4b, Figs. S8-S10). For example, when based on metabolites, marine animal proximal gut was once misclassified as seawater, marine sediment was once misclassified as non-saline animal distal gut, freshwater was twice misclassified as seawater, and seawater was once misclassified as marine animal secretion (i.e., during a single iteration) (Fig. 4b). We note that the majority of misclassifications were between compositionally similar environments (Fig. 3b, Fig. 4b, Figs. S8-S10). Features identified here as important for classification should prove useful as indicators of particular environments (Fig. 5), which can be used for applications such as source tracking ^66^ and forensics ^67^. When considering the twenty most highly ranked metabolites regarding impacting classification performance, metabolites classified to the pathways amino acids and peptides were not present (Fig. 5a), although metabolites from this pathway were differentially abundant across samples (Fig. 3c, Table S3, Table S4). Rather, the most abundant and highly ranked pathway among those highly predictive metabolites was for terpenoids, highlighting the importance of this group of metabolites in distinguishing Earth’s environments (Fig. 5a, Table S6). Terpenoids are the largest class of natural products recognized to date, and are known to be the most prevalent secondary metabolites in nature ^68^, which we also showed with our presence/absence data (Fig. 2a). Although known most commonly from plants, recent work has described a diversity of terpenoids produced by microbes, which range in activity from stress responses to signaling and communication ^68^. Future work should aim to further characterize the terpenoids discovered in this dataset.

We also identified metabolite–microbe co-occurrences, and as a first step towards characterizing them as salient features of the environment, showed that these relationships can be specific to certain habitats (Fig. 6, Fig. S4). We view strongly co-occurring metabolite–microbe pairs as features of the environment that in part can be grappled for further exploration, for example as predictors in models of environmental change ^69^. We demonstrated that both distinct metabolite pathways (e.g., carbohydrates vs. terpenoids) and metabolites within the same pathway (e.g., two groups of fatty acids), can be used to distinguish environments based on their co-occurrences with microbes (Fig. 6c-f, Fig. S4f-j). Similarly, we showed that certain metabolites and microbes have an especially high number of strong co-occurrences with one another (Fig. S11, Fig. S12). We hypothesize that microbes co-occurring with relatively many metabolites represent ‘chemically-talented’ taxa that may be useful for discovery of novel compounds. Further culture-based studies should continue to explore and characterize the metabolic diversity among these microbes.

Here, we described patterns of turnover, nestedness, and co-occurrence of metabolites and microbes across a diverse set of environments while addressing ecological questions surrounding the distribution of metabolites and their relationships with microbial diversity. Our results highlight the advantages of using standardized methods and a multi-omics approach including metagenomics and metabolomics to interpret and predict the contributions of microbes and their environments to chemical profiles in nature. One outstanding question in microbial ecology asks how microbial taxon profiles can be married with functional ones ^70^. Here, in addition to describing microbial taxa, their functions, and their metabolites, we explicitly tested for metabolite–microbe co-occurrences and explored how they relate to the environment, for which we have outlined our approach (Fig. S1). We recognize that previous studies describing microbial taxa and function using globally distributed sample sets, such as for the human gut, soils, and the ocean, have shown that both can vary across locations ^71–74^. Similarly, studies examining metabolite profiles across changes in microbial community composition, or environmental stress such as from heat, have shown variation associated with either ^75, 76^ or both ^77^. Furthermore, among previous multi-omics studies combining metagenomics with metatranscriptomics, metaproteomics, and/or metabolomics, some of which have shown the correlation between data layers to vary across sites, the majority are focused on a single environment ^78–88^. To our knowledge this is its first application of multi-omics integration of a dataset encompassing a diversity of environmental sample types representing several habitats, generated using standardized methods allowing for robust meta-analysis. Standardization of methods is of utmost importance, as no single lab can sample everything, and because a multitude of methods for performing a microbiome study exist ^89–91^. Due to inherent biases among distinct methods such as towards describing particular taxa ^91–92^, such lack of standardization prevents robust meta-analysis ^1, 2, 4^. Issues surrounding such bias extend from sequencing to metabolomics, which may be subject to greater technical variation due in part to unavoidable batch effects and use of extraction methods unique to particular sample types ^93^. The EMP500 overcomes these challenges by using standardized approaches, allowing for robust tracking of microbes and metabolites that permits the description of features that distinguish one habitat from another. This insight fosters understanding of the processes that make each habitat unique, and that may be vital to the functional diversity in the environment. By using standardized methods for sample processing and data analysis, the EMP500 allows for additional contributions, further expanding our insight into these communities.

We argue that using only sequence-based approaches to interpret functional potential can be misleading, as the presence of genomic loci and/or transcripts does not equate to the presence of a functional product in the environment. Using our presence/absence data for metabolites, we observed a trend in the uniform distribution of metabolite pathways across environments (Fig. 2a,c). However, when taking into account the relative intensities of metabolites – to date only possible using metabolomics – we observed significant differences in the distribution of particular groups of metabolites across the Earth’s environments (Fig. 2b,d, Fig. 3a). This emphasizes the utility and importance of directly measuring functional products in the environment, rather than estimating their potential from underlying genomic elements. We note that the uniform distributions of metabolite pathways and superclasses across environments based on presence/absence data (Fig. 2a,c) are similar to previous observations based on BGC annotation of a global dataset of MAGs^20^. It could be that abundance/intensity data for the products of BGCs may provide a different view, as they have here. We also recognize that using only metabolomics-based approaches can make the detection of certain molecules difficult, as some metabolites have relatively short lifespans, are consumed rapidly, and/or are cycled between members of the community therefore escaping detection ^94, 95^.

Beyond the important ecological questions explored here, several others such as those surrounding host-microbe interactions, microbial ecology in a changing world, and environmental processes merit future exploration ^70^. In some cases, addressing these questions will only be feasible following the collection of additional samples that span additional environments and/or geographic locations. For example, although we explored turnover and nestedness, one major question is whether these communities conform to the same biogeographic and ecological principles as in other types of communities, such as those of animals or plants ^70, 96, 97^. For example, we were unable to explore whether our features follow the latitudinal diversity gradient. The increase in species richness towards lower latitudes is apparent in many populations including those of several animals and plant species, but also planktonic marine bacteria ^98^ and soilborne *Streptomyces* ^99^. This trend has been less-explored at the community level, outside of soils ^100, 101^. Although highly host-specific groups such as ectomycorrhizal fungi do not follow this gradient due to the distributions of their host populations^102^, it is unclear what pattern metabolites and microbes exhibit, and whether there is variation among all of the environments recognized here. As another example, we did not explicitly explore the importance of rare features with respect to differences among environments. In addition to rare features serving as potential indicators of particular interactions ^103^ or ecological trends ^104, 105^, little is known regarding the relationships between rare features from distinct data layers (e.g., metabolites and microbial taxa). Although we might expect metabolites produced by rare microbes to also be rare in the environment, the suite of community interactions acting on those metabolites may alter distributions in context-dependent ways.

Our approach illustrates that recent advances in computational annotation tools offer a powerful toolbox to interpret untargeted metabolomics data ^45^. We anticipate that advances in metagenomic sequencing, genome assembly, and genome mining will improve the discovery and classification of functional products from among microbes and provide additional insight into these findings. By following standardized methods available on GitHub and making this dataset publicly available in Qiita and GNPS, this study will serve as an important resource for continued collaborative investigations. In the same manner, the development of novel instrumentation and computational methods for metabolomics will expand the depth of metabolites surveyed in microbiome studies.

## Acknowledgements

We thank Gennadi Milivenvsky, Anders Møller, Igor Chizhevsky, Serhii Kirieiev, Anatoly Nosovsky, and Maksym Ivanenko for logistic support with fieldwork in Ukraine; Lindsay Goldasich and Julia Toronczak for assistance with sample processing for sequencing; Joshua Ladau for assistance running nestedness scripts; Marcus Fedarko, Rachel Diner, Joshua Ladau, Elisha Wood-Charlson, Stephen Nayfach, Daniel Udwary, and Emiley Eloe-Fadrosh for reviewing the manuscript. This work was supported in part by the Samuel Freeman Charitable Trust, United States (US) National Institute of Health (NIH) (awards 1RF1-AG058942-01, 1DP1AT010885, R01HL140976, R01DK102932, R01HL134887, U19AG063744, U01AI124316), US Department of Agriculture – National Institute of Food and Agriculture (USDA-NIFA) (award 2019-67013-29137), the US National Science Foundation (NSF) - Center for Aerosol Impacts on Chemistry of the Environment, Crohn’s & Colitis Foundation Award (CCFA) (award 675191), US Department of Energy - Office of Science - Office of Biological and Environmental Research - Environmental System Science Program, Semiconductor Research Corporation and Defense Advanced Research Projects Agency (SRC/DARPA) (award GI18518), Department of Defense (award W81XWH-17-1-0589), the Office of Naval Research (ONR) (award N00014-15-1-2809), the Emerald Foundation (award 3022), IBM Research AI through the AI Horizons Network, and the Center for Microbiome Innovation. J.P.S. was supported by NIH/NIGMS IRACDA K12 GM068524. L.F.N. was supported by the NIH (award R01-GM107550). A.D.B. was supported by the Danish Council for Independent Research (DFF) (award 9058-00025B). W.B. was supported by the Research Foundation – Flanders (12W0418N). K.D. and S.B. were supported by Deutsche Forschungsgemeinschaft (BO 1910/20 and 1910/23). P.C.D. was supported by the Gordon and Betty Moore Foundation (award GBMF7622) and the NIH (award R01-GM107550). Metabolomics analyses at Pacific Northwest National Laboratory (PNNL) were supported by the Laboratory Directed Research and Development program via the Microbiomes in Transition Initiative and performed in the Environmental Molecular Sciences Laboratory, a national scientific user facility sponsored by the US Office of Biological and Environmental Research and located at PNNL. This contribution originates in part from the River Corridor Scientific Focus Area project at PNNL. PNNL is a multiprogram national laboratory operated by Battelle for the Department of Energy (DOE) under contract DE-AC05-76RLO 1830. We thank Eppendorf, Illumina, and Integrated DNA Technologies for in-kind support at various phases of the project.

## Author contributions

The EMP500 Consortium collected and provided samples. J.A.G., J.K.J., and R.K. conceived the idea for the project. P.C.D., and R.K. designed the multi-omics component of the project and provided project oversight. J.P.S. managed the project, performed preliminary data exploration, coordinated data analysis, analyzed data, and provided data interpretation. L.F.N. coordinated and performed LC-MS/MS analysis, and the processing, annotation, and interpretation of LC-MS/MS data. M.E.-N. performed sample preparation and extraction prior to LC-MS/MS analysis. L.R.T. designed the multi-omics component of the project, solicited sample collection, curated sample metadata, processed samples, performed preliminary data exploration, and provided project oversight. J.G.S. designed the multi-omics component, managed the project, developed protocols and tools, coordinated and performed sequencing, and performed preliminary exploration of sequence data. R.A.S. developed protocols and coordinated and performed sequencing. S.P.C. and T.O.M. coordinated and performed GC-MS sample processing and provided interpretation of GC-MS data. A.D.B. conceived the idea for the paper, performed preliminary data exploration, analyzed data, and provided data interpretation. S.H. performed machine-learning analyses. F.L. performed co-occurrence analysis, multinomial regression analyses, and correlations with co-occurrence data. H.L.L. performed multinomial regression analyses. Q.Z. developed tools and provided interpretation of shotgun metagenomics data. C.Mart. and J.T.M. provided oversight and interpretation of RPCA, multinomial regression, and co-occurrence analyses. S.K. performed preliminary exploration of shotgun metagenomics data. K.D., S.B, and H.W.K. contributed to the annotation of LC-MS/MS data. A.A.A. processed GC-MS data. W.B. provided oversight for machine-learning analyses. C.Maro. processed samples for sequencing. Y.V.B. performed preliminary data exploration and provided oversight for machine-learning analysis. A.T. and D.P. performed preliminary data exploration. L.P., A.P.C., N.H., and K.L.B. performed preliminary exploration of shotgun metagenomic data and performed machine learning analyses. P.D. performed preliminary exploration of shotgun metagenomics data. A.G. developed tools, provided interpretation of shotgun metagenomics data, and analyzed shotgun metagenomics data. G.H. coordinated short-read amplicon and shotgun metagenomics sequencing. M.M.B. and K.S. performed short-read amplicon and shotgun metagenomics sequencing. T.S. assisted with DNA extraction. D.M. coordinated long-read amplicon sequencing, analyzed shotgun metagenomics data, and provided interpretation of the data. S.M.K. and M.A. coordinated and performed long-read amplicon sequencing and long-read sequence data analysis. J.J.M. collected samples, coordinated field logistics, developed protocols, and performed short-read amplicon and shotgun metagenomics sequencing. S.S. collected samples, coordinated field logistics, and provided interpretation of the data. G.L.A. curated sample metadata and organized sequence data. J.D. processed sequence data. A.D.S. provided project oversight and data interpretation. T.T., A.S., and J.S. collected samples, coordinated field logistics, and provided interpretation of the data. J.P.S. wrote the manuscript, with contributions from all authors.

## Earth Microbiome Project 500 (EMP500) Consortium

Lars T. Angenant^1^, Alison M. Berry^2^, Leonora S. Bittleston^3^, Jennifer L. Bowen^4^, Max Chavarría^5,6^, Don A. Cowan^7^, Dan Distel^4^, Peter R. Girguis^8^, Jaime Huerta-Cepas^9^, Paul R. Jensen^10^, Lingjing Jiang^11^, Gary M. King^12^, Anton Lavrinienko^13^, Aurora MacRae-Crerar^14^, Thulani P. Makhalanyane^7^, Tapio Mappes^13^, Ezequiel M. Marzinelli^15^, Gregory Mayer^16^, Katherine D. McMahon^17^, Jessica L. Metcalf^18^, Sou Miyake^19^, Timothy A. Mousseau^13^, Catalina Murillo-Cruz^5^, David Myrold^20^, Brian Palenik^10^, Adrián A. Pinto-Tomás^5^, Dorota L. Porazinska^21^, Jean-Baptiste Ramond^7,22^, Forest Rowher^23^, Taniya RoyChowdhury^24,25^, Stuart A. Sandin^10^, Steven K. Schmidt^26^, Henning Seedorf^19,27^, J. Reuben Shipway^28,29^, Jennifer E. Smith^10^, Frank J. Stewart^30^, Karen Tait^31^, Yael Tucker^32^, Jana M. U’Ren^33^, Phillip C. Watts^13^, Nicole S. Webster^34,35^, Jesse R. Zaneveld^36^, Shan Zhang^37^

^1^University of Tuebingen, Tuebingen, Germany. ^2^University of California, Davis, Davis, California, USA. ^3^Boise State University, Boise, Idaho, USA. ^4^Northeastern University, Boston, Massachusetts, USA. ^5^University of Costa Rica, San José, Costa Rica. ^6^CENIBiot, San José, Costa Rica. ^7^University of Pretoria, Pretoria, South Africa. ^8^Harvard University, Cambridge, Massachusetts, USA. ^9^Universidad Politécnica de Madrid, Instituto Nacional de Investigación y Tecnología Agraria y Alimentaria, Madrid, Spain. ^10^University of California San Diego, La Jolla, California, USA. ^11^Janssen Research & Development, San Diego, California, USA. ^12^Louisiana State University, Baton Rouge, Louisiana, USA. ^13^University of Jyväskylä, Jyväskylä, Finland. ^14^University of Pennsylvania, Philadelphia, Pennsylvania, USA. ^15^The University of Sydney, Sydney, Australia. ^16^Texas Technology University, Lubbock, Texas, USA. ^17^University of Wisconsin, Madison, Wisconsin, USA. ^18^Colorado State University, Fort Collins, Colorado, USA. ^19^Temasek Life Sciences Laboratory, Singapore, Singapore. ^20^Oregon State University, Corvallis, Oregon, USA. ^21^University of Florida, Gainesville, Florida, USA. ^22^Pontificia Universidad Católica de Chile, Santiago, Chile. ^23^San Diego State University, San Diego, California, USA. ^24^Pacific Northwest National Laboratory, Richland, Washington, USA. ^25^University of Maryland, College Park, Maryland, USA. ^27^National University of Singapore, Singapore, Singapore. ^28^University of Plymouth, Drake Circus, Plymouth, United Kingdom. ^29^University of Massachusetts Amherst, Amherst, Massachusetts, USA. ^30^Montana State University, Bozeman, Montana, USA. ^31^Plymouth Marine Laboratory, Plymouth, United Kingdom. ^32^National Energy Technology Laboratory, USA. ^33^University of Arizona, Tucson, Arizona, USA. ^34^Australian Institute of Marine Science, Townsville, Qld, Australia. ^35^University of Queensland, Brisbane, Australia. ^36^University of Washington Bothell, Bothell, Washington, USA. ^37^University of New South Wales, Sydney, Australia.

## ONLINE METHODS

### DATASET DESCRIPTION

#### Sample collection

Samples were contributed by 34 principal investigators (PIs) of the Earth Microbiome Project 500 (EMP500) Consortium. Samples were contributed as distinct sets referred to here as studies, where each study represented a single environment (e.g., terrestrial plant detritus). To achieve more even coverage across microbial environments, we devised an ontology of sample types (microbial environments), the EMP Ontology (EMPO) (http://earthmicrobiome.org/protocols-and-standards/empo/)^1^ and selected samples to fill out EMPO categories as broadly as possible. As we anticipated previously^1^, we have updated the number of levels as well as states therein for EMPO (Fig. 1b), based on an important, additional salinity gradient observed among host-associated samples when considering the novel shotgun metagenomic and metabolomic data generated here (Fig. 3c,d). We note that although we were able to acquire samples for all EMPO categories, some categories are represented by a single study.

Samples were collected following the Earth Microbiome Project sample submission guide^2^. Briefly, samples were collected fresh, split into 10 aliquots, and then frozen, or alternatively collected and frozen, and subsequently split into 10 aliquots with minimal perturbation. Aliquot size was sufficient to yield 10–100 ng genomic DNA (approximately 10^7^– 10^8^ cells). To leave samples amenable to chemical characterization (metabolomics), buffers or solutions for sample preservation (e.g., RNAlater) were avoided. Ethanol (50–95%) was allowed as it is compatible with LC-MS/MS though should also be avoided if possible.

Sampling guidance was tailored for four general sample types: bulk unaltered (e.g., soil, sediment, feces), bulk fractionated (e.g., sponges, corals, turbid water), swabs (e.g., biofilms), and filters. Bulk unaltered samples were split fresh (or frozen) sampled into 10 pre-labeled 2-mL screw-cap bead beater tubes (Sarstedt cat. no. 72.694.005 or similar), ideally with at least 200 mg biomass, and flash frozen in liquid nitrogen (if possible). Bulk fractionated samples were fractionated as appropriate for the sample type, split into 10 pre-labeled 2-mL screw-cap bead beater tubes, ideally with at least 200 mg biomass, and flash frozen in liquid nitrogen (if possible). Swabs were collected as 10 replicate swabs using 5 BD SWUBE dual cotton swabs with wooden stick and screw cap (cat. no. 281130). Filters were collected as 10 replicate filters (47 mm diameter, 0.2 um pore size, polyethersulfone (preferred) or hydrophilic PTFE filters), placed in a pre-labeled 2-mL screw-cap bead beater tubes, and flash frozen in liquid nitrogen (if possible). All sample types were stored at –80 °C if possible, otherwise –20 °C.

To track the provenance of sample aliquots, we employed a QR coding scheme. Labels were affixed to aliquot tubes before shipping, when possible. QR codes had the format “name.99.s003.a05”, where “name” is the PI name, “99” is the study ID, “s003” is the sample number, and “a05” is the aliquot number. QR codes (version 2, 25 pixels x 25 pixels) were printed on 1.125” x 0.75” rectangular and 0.437” circular cap Cryogenic Direct Thermal Labels (GA International, part no. DFP-70) using a Zebra model GK420d printer and ZebraDesigner Pro software for Windows. After receipt but before aliquots were stored in freezers, QR codes were scanned into a sample inventory spreadsheet using a QR scanner.

#### Sample metadata

Environmental metadata was collected for all samples based on the EMP Metadata Guide2, which combines guidance from the Genomics Standards Consortium MIxS (Minimum Information about any Sequence) standard3 and the Qiita Database (https://qiita.ucsd.edu)4. The metadata guide provides templates and instructions for each MIxS environmental package (i.e., sample type). Relevant information describing each PI submission, or study, was organized into a separate study metadata file (Table S1).

### METABOLOMICS

#### LC-MS/MS sample extraction and preparation

To profile metabolites among all samples, we used liquid chromatography with untargeted tandem mass spectrometry (LC-MS/MS), a versatile method that detects tens of thousands of metabolites in biological samples^12^. All solvents and reactants used were LC-MS grade. To maximize the biomass extracted from each sample, the samples were prepared depending on their sampling method (e.g., bulk, swabs, filter, and controls). The bulk samples were transferred into a microcentrifuge tube (polypropylene, PP) and dissolved in 7:3 MeOH:H_2_O using a volume varying from 600 µL to 1.5 mL, depending on the amounts of sample available, and homogenized in a tissue-lyser (QIAGEN) at 25 Hz for 5 min. Then, the tubes were centrifuged at 15,000 rpm for 15 min, and the supernatant was collected in a 96-well plate (PP). For swabs, the swabs were transferred into a 96-well plate (PP) and dissolved in 1.0 mL of 9:1 EtOH:H_2_O. The prepared plates were sonicated for 30 min, and after 12 hours at 4°C, the swabs were removed from the wells. The filter samples were dissolved in 1.5 mL of 7:3 MeOH:H_2_O in microcentrifuge tubes (PP) and were sonicated for 30 min. After 12 hours at 4°C, the filters were removed from the tubes. The tubes were centrifuged at 15,000 rpm for 15 min, and the supernatants were transferred to 96-well plates (PP). The process control samples (bags, filters, and tubes) were prepared by adding 3.0 mL of 2:8 MeOH:H_2_O and by recovering 1.5 mL after 2 min. After the extraction process, all sample plates were dried with a vacuum concentrator and subjected to solid phase extraction (SPE). The SPE is used to remove salts that could reduce ionization efficiency during mass spectrometry analysis, as well as the most polar and non-polar compounds (e.g., waxes) that cannot be analyzed efficiently by reversed-phase chromatography. The protocol was as follows: The samples (in plates) were dissolved in 300 µL of 7:3 MeOH:H_2_O and put in an ultrasound bath for 20 min. SPE was performed with SPE plates (Oasis HLB, Hydrophilic-Lipophilic-Balance, 30 mg with particle sizes of 30-µm). The SPE beds were activated by priming them with 100% MeOH, and equilibrated with 100% H_2_O. The samples were loaded on the SPE beds, and 100% H_2_O was used as wash solvent (600 µL). The eluted washing solution was discarded, as it contains salts and very polar metabolites that subsequent metabolomics analysis is not designed for. The sample elution was carried out sequentially with 7:3 MeOH:H_2_O (600 µL) and with 100% MeOH (600 µL). The obtained plates were dried with a vacuum concentrator. For mass spectrometry analysis, the samples were resuspended in 130 µL of 7:3 MeOH:H_2_O containing 0.2 µM of amitriptyline as an internal standard. The plates were centrifuged at 2,000 rpm for 15 min at 4°C. 100 µL of samples were transferred into a new 96-well plate (PP) for mass spectrometry analysis.

#### LC-MS/MS sample analysis

The extracted samples were analyzed by ultra-high performance liquid chromatography (UHPLC, Vanquish, Thermo Scientific, Waltham, Massachusetts, USA) coupled to a quadrupole-Orbitrap mass spectrometer (Q Exactive, Thermo Scientific, Waltham, Massachusetts, USA) operated in data-dependent acquisition mode (LC-MS/MS in DDA mode). Chromatographic separation was performed using a Kinetex C_18_ 1.7-µm (Phenomenex, Torrance, California, USA), 100-Å pore size, 2.1-mm (internal diameter) x 50-mm (length) column with a C_18_ guard cartridge (Phenomenex). The column was maintained at 40°C. The mobile phase was composed of a mixture of (A) water with 0.1% formic acid (v/v) and (B) acetonitrile with 0.1% formic acid. Chromatographic elution method was set as follows: 0.00–1.00 min, isocratic 5% B; 1.00–9.00 min, gradient from 5% to 100% B; 9.00–11.00 min, isocratic 100% B; and followed by equilibration 11.00–11.50 min, gradient from 100% to 5% B; 11.50–12.50 min, isocratic 5% B. The flow rate was set to 0.5 mL/min.

The UHPLC was interfaced to the orbitrap using a heated electrospray ionization (HESI) source with the following parameters: ionization mode: positive, spray voltage, +3496.2 V; heater temperature, 363.90 °C; capillary temperature, 377.50 °C; S-lens RF, 60 (arb. units); sheath gas flow rate, 60.19 (arb. units); and auxiliary gas flow rate, 20.00 (arb. units). The MS^1^ scans were acquired at a resolution (at *m/z* 200) of 35,000 in the *m/z* 100–1500 range, and the fragmentation spectra (MS^2^) scans at a resolution of 17,500 from 0 to 12.5 min. The automatic gain control (AGC) target and maximum injection time were set at 1.0 x 10^6^ and 160 ms for MS^1^ scans, and set at 5.0 x 10^5^ and 220 ms for MS^2^ scans, respectively. Up to three MS^2^ scans in data-dependent mode (Top 3) were acquired for the most abundant ions per MS^1^ scans using the apex trigger mode (4 to 15 s), dynamic exclusion (11 s), and automatic isotope exclusion. The starting value for MS^2^ was *m/z* 50. Higher-energy collision induced dissociation (HCD) was performed with a normalized collision energy of 20, 30, 40 eV in stepped mode. The major background ions originating from the SPE were excluded manually from the MS^2^ acquisition. Analyses were randomized within plate and blank samples analyzed every 20 injections. A QC mix sample assembled from 20 random samples across the sample-types was injected at the beginning, the middle, and the end of each plate sequence. The chromatographic shift observed throughout the batch is estimated as less than 2 s, and the relative standard deviation of ion intensity was 15% per replicates.

#### LC-MS/MS data processing

The mass spectrometry data were centroided and converted from the proprietary format (.raw) to the *m/z* extensible markup language format (.mzML) using ProteoWizard (ver. 3.0.19, MSConvert tool)^5^. The mzML files were then processed with MZmine toolbox^6^ using the ion-identity networking modules^7^ that allows advanced detection for adduct/isotopologue annotations. The MZmine processing was performed on Ubuntu 18.04 LTS 64-bits workstation (Intel Xeon E5-2637, 3.5 GHz, 8 cores, 64 Gb of RAM) and took ∼3 d. The MZmine project, the MZmine batch file (.XML format), and results files (.MGF and .CSV) are available in the MassIVE dataset <u>MSV000083475</u>. The MZmine batch file contains all the parameters used during the processing. In brief, feature detection and deconvolution was performed with the ADAP chromatogram builder^8^, and local minimum search algorithm. The isotopologues were regrouped, and the features (peaks) were aligned across samples. The aligned peak list was gap filled and only peaks with an associated fragmentation spectrum and occurring in a minimum of three files were conserved. Peak shape correlation analysis grouped peaks originating from the same molecule, and to annotate adduct/isotopologue with ion-identity networking^7^. Finally the feature quantification table results (.CSV) and spectral information (.MGF) were exported with the GNPS module for feature-based molecular networking analysis on GNPS^9^ and with SIRIUS export modules.

#### LC-MS/MS data annotation

The results files of MZmine (.MGF and .CSV files) were uploaded to GNPS (http://gnps.ucsd.edu)^10^ and analyzed with the feature-based molecular networking workflow^9^. Spectral library matching was performed against public fragmentation spectra (MS^2^) spectral libraries on GNPS and the NIST17 library.

For the additional annotation of small peptides, we used the DEREPLICATOR tools available on GNPS^11, 12^. We then used SIRIUS^13^ (v. 4.4.25, headless, Linux) to systematically annotate the MS^2^ spectra. Molecular formulas were computed with the SIRIUS module by matching the experimental and predicted isotopic patterns^14^, and from fragmentation trees analysis^15^ of MS^2^. Molecular formula prediction was refined with the ZODIAC module using Gibbs sampling^16^ on the fragmentation spectra (chimeric spectra or had a poor fragmentation were excluded). *In silico* structure annotation using structures from biodatabase was done with CSI:FingerID^17^. Systematic class annotations were obtained with CANOPUS^18^ and used the NPClassifier ontology^19^.

The parameters for SIRIUS tools were set as follows, for SIRIUS: molecular formula candidates retained (80), molecular formula database (ALL), maximum precursor ion *m/z* computed (750), profile (orbitrap), *m/z* maximum deviation (10 ppm), ions annotated with MZmine were prioritized and other ions were considered (i.e., [M+H3N+H]+, [M+H]+, [M+K]+,[M+Na]+, [M+H-H2O]+, [M+H-H4O2]+, [M+NH4]+); for ZODIAC: the features were split into 10 random subsets for lower computational burden and computed separately with the following parameters: threshold filter (0.9), minimum local connections (0); for CSI:FingerID: *m/z* maximum deviation (10 ppm) and biological database (BIO).

To establish putative microbially-related secondary metabolites, we collected annotations from spectral library matching and the DEREPLICATOR tools and queried them against the largest microbial metabolite reference databases (Natural Products Atlas^20^ and MIBiG^21^). Molecular networking^9^ was then used to propagate the annotation of microbially-related secondary metabolites throughout all molecular families (i.e., the network component).

#### LC-MS/MS data analysis

We combined the annotation results from the different tools described above to create a comprehensive metadata file describing each metabolite feature observed. Using that information, we generated a feature-table including only secondary metabolite features determined to be microbially-related. We then excluded very low-intensity features introduced to certain samples during the gap-filing step described above. These features were identified based on presence in negative controls that were universal to all sample types (i.e., bulk, filter, and swab), and by their relatively low per-sample intensity values. Finally, we excluded features present in positive controls for sampling devices specific to each sample type (i.e., bulk, filter, or swab). The final feature-table included 618 samples and 6,588 putative microbially-related secondary metabolite features that were used for subsequent analysis.

We used QIIME 2’s^22^ *diversity* plugin to quantify alpha-diversity (i.e., feature richness) for each sample, and *deicode*^23^ to quantify beta-diversity (i.e., robust Aitchison distances, which are robust to both sparsity and compositionality in the data) between each pair of samples. We parameterized our robust Aitchison Principal Components Analysis (RPCA)^23^ to exclude samples with fewer than 500 features, and features present in fewer than 10% of samples. We used the *taxa* plugin to quantify the relative abundance of microbially-related secondary metabolite pathways and superclasses (i.e., based on NPClassifier) within each environment (i.e., for each level of EMPO 4), and *songbird*^24^ to identify sets of microbially-related secondary metabolites whose abundances were associated with certain environments. We parameterized our *songbird* model as follows: epochs = 1,000,000, differential prior = 0.5, learning rate = 1.0 x 10^-5^, summary interval = 2, batch size = 400, minimum sample count = 0, and training on 80% of samples at each level of EMPO 4, using ‘Animal distal gut (non-saline)’ as the reference environment. Environments with fewer than 10 samples were excluded to optimize model training (i.e., ‘Animal corpus [non-saline]’, ‘Animal proximal gut [non-saline]’, ‘Surface [saline]’). The output from *songbird* includes a rank value for each metabolite in every environment, which represents the log fold change for a given metabolite in a given environment^24^. We compared log fold changes for each metabolite from this run to those from (1) a replicate run using the same reference environment and (2) a run using a distinct reference environment: ‘Water (saline)’. We found strong Spearman correlations in both cases (Table S7), and therefore focused on results from the original run using ‘Animal distal gut (non-saline)’ as the reference environment, as it has previously been shown to be relatively unique among other habitats^1^. In addition to summarizing the top 10 metabolites for each environment (Table S3), we used the log fold change values in our multi-omics analyses described below.

We used the RPCA biplot and EMPeror^25^ to visualize differences in composition among samples, as well as the association with samples of the 25 most influential microbially-related secondary metabolite features (i.e., those with the largest magnitude across the first three principal component loadings). We tested for significant differences in metabolite composition across all levels of EMPO using permutational multivariate analysis of variance (PERMANOVA), implemented with QIIME 2’s *diversity* plugin^22^ and using our robust Aitchison distance matrix as input. In parallel, we used the differential abundance results from *songbird* described above to identify specific microbially-related secondary metabolite pathways and superclasses that varied strongly across environments. We then went back to our metabolite feature-table to visualized differences in the relative abundances of those pathways and superclasses within each environment by first selecting features and calculating log-ratios using *qurro*^26^, and then plotting using the *ggplot2* package^27^ in R^28^. We tested for significant differences in relative abundances across environments using Kruskal–Wallis tests, implemented using the base *stats* package in R^28^.

#### GC-MS sample extraction and preparation

To profile volatile small molecules among all samples in addition to what was captured with LC-MS/MS, we used gas chromatography coupled with mass spectrometry (GC-MS). All solvents and reactants were GC-MS grade. Two protocols were used for sample extraction, one for the 105 soil samples and second for the 356 fecal and sediment samples that were treated as biosafety level 2. The 105 soil samples were received at the Pacific Northwest National Laboratory and processed as follows. Each soil sample (1 g) was weighed into microcentrifuge tubes (Biopur Safe-Lock, 2.0 mL, Eppendorf, Hamburg, Germany). 1 mL of H_2_O and one scoop (∼0.5 g) of a 1:1 (v/v) mixture of garnet (0.15-mm, Omni International, Kennesaw, Georgia, USA) and stainless steel (SS) (0.9 – 2.0-mm blend, Next Advance, Troy, New York) beads and one 3-mm SS bead (Qiagen, Hilden, Germany) was added to each tube. Samples were homogenized in a tissue lyser (Qiagen, Hilden, Germany) for 3 min at 30 Hz and transferred into 15-mL polypropylene tubes (Olympus, Genesee Scientific, San Diego, California, USA). Ice-cold water (1 mL) was used to rinse the smaller tube and combined into the 15-mL tube. 10 mL of 2:1 (v/v) chloroform:methanol added and samples were rotated at 4°C for 10 min followed by cooling at -70°C for 10 min and centrifuged at 4,000 rpm for 10 min to separate phases. The top and bottom layers were combined into 40 mL glass vials and dried using a vacuum concentrator. 1 mL of 2:1 chloroform:methanol was added to each large glass vial and the sample was transferred into 1.5-mL tubes and centrifuged at 12,000 g. The supernatant was transferred into glass vials and dried for derivatization.

The remaining 356 samples that were received from UCSD that included fecal and sediment samples were processed as follows: 100 µL of each sample was transferred to a 2 mL microcentrifuge tube using a scoop (MSP01, Next Advance, Tustin, California, USA). The final volume of sample was brought to 1.5 mL ensuring the solvent ratio is 3:8:4 H_2_O:CHCl_3_:MeOH by adding the appropriate volumes of H_2_O, MeOH, and CHCl_3_. After transfer, one 3-mm SS bead (QIAGEN), 400 µL of methanol, and 300 µL of H2O were added to each tube and the samples were vortexed for 30 s. Then, 800 µL of chloroform was added and samples were vortexed for 30 s. After centrifuging at 4,000 rpm for 10 min to separate phases, the top and bottom layers were combined in a vial and dried for derivatization.

The samples were derivatized for GC-MS analysis as follows: 20 µL of a methoxyamine solution in pyridine (30 mg/mL) was added to the sample vial and vortexed for 30 s. A bath sonicator was used to ensure the sample was completely dissolved. Samples were incubated at 37°C for 1.5 h while shaking at 1,000 rpm. 80 µL of N-methyl-N-trimethylsilyltrifluoroacetamide and 1% trimethylchlorosilane (MSTFA) solution was added and samples were vortexed for 10 s, followed by incubation at 37°C for 30 min with 1000 rpm shaking. The samples were then transferred into a vial with an insert.

An Agilent 7890A gas chromatograph coupled with a single quadrupole 5975C mass spectrometer (Agilent Technologies, Santa Clara, California, USA) and a HP-5MS column (30- m × 0.25-mm × 0.25-μm; Agilent Technologies, Santa Clara, California, USA) was used for untargeted analysis. Samples (1 μL) were injected in splitless mode, and the helium gas flow rate was determined by the Agilent Retention Time Locking function based on analysis of deuterated myristic acid (Agilent Technologies, Santa Clara, California, USA). The injection port temperature was held at 250°C throughout the analysis. The GC oven was held at 60°C for 1 min after injection, and the temperature was then increased to 325°C by 10°C/min, followed by a 10 min hold at 325°C. Data were collected over the mass range of *m/z* 50–600. A mixture of FAMEs (C8–C28) was analyzed each day with the samples for retention index alignment purposes during subsequent data analysis.

#### GC-MS data processing and annotation

The data were converted from vendor’s format to the .mzML format and processed using GNPS GC-MS data analysis workflow (https://gnps.ucsd.edu)^29^. The compounds were identified by matching experimental spectra to the public libraries available at GNPS, as well as NIST 17 and Wiley libraries. The data are publicly available at the MassIVE depository (https://massive.ucsd.edu); dataset ID: MSV000083743. The GNPS deconvolution is available on GNPS (https://gnps.ucsd.edu/ProteoSAFe/status.jsp?task=d5c5135a59eb48779216615e8d5cb3ac), as is the library search (https://gnps.ucsd.edu/ProteoSAFe/status.jsp?task=59b20fc8381f4ee6b79d35034de81d86).

#### GC-MS data analysis

For multi-omics analyses including GC-MS data, we first removed noisy (i.e., suspected background contaminant- and artifactual) features by excluding those with balance scores <50%. Balance scores describe compositional consistency of deconvoluted spectra across the dataset, where high values indicate reproducible spectral patterns and thus high-quality spectra. We then used QIIME 2’s *deicode*^23^ plugin to estimate beta-diversity for each dataset using robust Aitchison distances. The final feature table for GC-MS beta-diversity analysis included 460 samples and 216 features.

### METAGENOMICS

#### DNA extraction

For each round of DNA extractions described below for both amplicon and shotgun metagenomic sequencing, a single aliquot of each sample was processed for DNA extraction. DNA was extracted following the EMP 96-sample, magnetic bead-based DNA extraction protocol^30^, following Marotz et al. (2017)^31^, Minich et al. (2018)^32^, and Minich et al. (2019)^33^, and using the QIAGEN® MagAttract® PowerSoil® DNA KF Kit (384-sample) (i.e., optimized for KingFisher). Importantly, material from each sample was added to a unique bead tube (containing garnet beads) for single-tube lysis, which has been shown to reduce sample-to-sample contamination common in plate-based extractions^33^. For bulk samples, 0.1 to 0.25 g of material was added to each well; for filtered samples, one entire filter was added to each well; for swabbed samples, one swab head was added to each well. The lysis solution was dissolved at 60°C before addition to each tube, then capped tubes were incubated at 65°C for 10 min prior to mechanical lysis at 6000 rpm for 20 min using a MagNA Lyser (Roche Diagnostics, California, USA). Lysate from each bead tube was then randomly assigned and added to wells of a 96-well plate, and then cleaned-up using the KingFisher Flex system (Thermo Scientific, Waltham, Massachusetts, USA). Resulting DNA was stored at –20°C for sequencing. We note that whereas QIAGEN does not offer a ‘hybrid’ extraction kit allowing for single-tube lysis and plate-based clean-up, the Thermo MagMAX Microbiome Ultra kit does, and was recently shown to be comparable to the EMP protocol used here^34^.

#### Amplicon sequencing

We generated amplicon sequence data for variable region four (V4) of the bacterial and archaeal 16S ribosomal RNA (rRNA) gene, variable region nine (V9) of the eukaryotic 18S rRNA gene, and the fungal internal transcribed spacer one (ITS1). For amplifying and sequencing all targets, we used a low-cost, miniaturized (i.e., 5-µL volume), high-throughput (384-sample) amplicon library preparation method implementing the Echo 550 acoustic liquid handler (Beckman Coulter, Brea, California, USA)^35^. The same protocol was modified with different primer sets and PCR cycling parameters depending on the target. Two rounds of DNA extraction and sequencing were performed for each target to obtain greater coverage per sample. For a subset of 500 samples, we also generated high-quality sequence data for full-length bacterial rRNA operons following the protocol described by Karst et al. (2021)^36^, which is outlined briefly below.

The protocol for 16S is outlined fully in Caporaso et al. (2018)^37^. To target the V4 region, we used the primers 515F (Parada) (5’-GTGYCAGCMGCCGCGGTAA-3’) and 806R (Apprill) (5’-GGACTACNVGGGTWTCTAAT-3’). These primers are updated from the original EMP 16S-V4 primer sequences^38, 39^ in order to (1) remove bias against Crenarchaeota/Thaumarchaeota^40^ and the marine freshwater clade SAR11 (Alphaproteobacteria)^41^, and (2) enable the use of various reverse primer constructs (e.g., the V4-V5 region using the reverse primer 926R^42^) by moving the barcode/index to the forward primer^40^. We note that whereas we previously named these updated primers “515FB” and “806RB” to distinguish them from the original primers, the “B” may be misinterpreted to indicate “Barcode”. To avoid ambiguity, we now use the original names suffixed with the lead author name (i.e., “515F (Parada)”, “806R (Apprill)”, and “926R (Quince)”. We highly recommend to always check the primer sequence in addition to the primer name. For Qiita users, studies with “library_construction_protocol” as “515f/806rbc” used the original primers, whereas “515fbc/806r” indicates use of updated primers, where “bc” refers to the location of barcode.

To facilitate sequencing on Illumina platforms, the following primer constructs were used to integrate adapter sequences during amplification^38, 39, 43^. For the barcoded forward primer, constructs included (5’ to 3’): the 5’ Illumina adapter (AATGATACGGCGACCACCGAGATCTACACGCT), a Golay barcode (12-bp variable sequence), a forward primer pad (TATGGTAATT), a forward primer linker (GT), and the forward primer (515F [Parada]) (GTGYCAGCMGCCGCGGTAA). For the reverse primer, constructs included (5’ to 3’): The reverse complement of 3’ Illumina adapter (CAAGCAGAAGACGGCATACGAGAT), a reverse primer pad (AGTCAGCCAG), a reverse primer linker (CC), and the reverse primer (806R [Apprill]) (GGACTACNVGGGTWTCTAAT).

For each 25-µL reaction, we combined 13 µL PCR-grade water (Sigma St. Louis, MO, USA, cat. no. W3500 or QIAGEN, Hilden, Germany, cat. no. 17000-10), 10 µL Platinum Hot Start PCR Master Mix (2X) (Thermo Scientific, Waltham, Massachusetts, USA, cat. no. 13000014), 0.5 µL of each primer (10 µM), and 1 µL of template DNA. The final concentration of the master mix in each 1X reaction was 0.8X and that of each primer was 0.2 µM. Cycling parameters for a 384-well thermal cycler were as follows: 94°C for 3 min; 35 cycles of 94°C for 1 min, 50°C for 1 min, and 72°C for 105 s; and 72°C for 10 min. For a 96-well thermal cycler, we recommend the following: 94°C for 3 min; 35 cycles of 94°C for 45 s, 50°C for 1 min, and 72°C for 90 s; and 72°C for 10 min.

We amplified each sample in triplicate (i.e., each sample was amplified in three replicate 25-µL reactions), and pooled products from replicate reactions for each sample into a single volume (75 µL). We visualized expected products between 300-350 bp on agarose gels, and note that whereas low-biomass samples may yield no visible bands, instruments such as a Bioanalyzer or TapeStation (Agilent, Santa Clara, California, USA) can be used to confirm amplification. We quantified amplicons using the Quant-iT PicoGreen dsDNA Assay Kit (Thermo Scientific, Waltham, MA, USA, cat. no. P11496), following the manufacturer’s instructions. To pool samples, we combined an equal amount of product from each sample (240 ng) into a single tube, and cleaned the pool using the UltraClean PCR Clean-Up Kit (QIAGEN, Hilden, Germany, cat. no. 12596-4), following the manufacturer’s instructions. We checked DNA quality using a Nanodrop (Thermo Scientific, Waltham, Massachusetts, USA), confirming A260/A280 ratios were between 1.8-2.0.

For sequencing, the following primer constructs were used. Read 1 constructs included (5’ to 3’): a forward primer pad (TATGGTAATT), a forward primer linker (GT), and the forward primer (515F [Parada]) (GTGYCAGCMGCCGCGGTAA). Read 2 constructs included (5’ to 3’): a reverse primer pad (AGTCAGCCAG), a reverse primer linker (CC), and the reverse primer (806R [Apprill]) (GGACTACNVGGGTWTCTAAT). The index primer sequence was AATGATACGGCGACCACCGAGATCTACACGCT, which we highlight as having an extra GCT at the 3’ end compared to Illumina’s index primer sequence, in order to increase the T_m_ for read 1 during sequencing.

The protocol for 18S is outlined fully in Amaral-Zettler et al. (2018)^44^. To target variable region nine (V9), we used the primers 1391f (5’-GTACACACCGCCCGTC-3’) and EukBr (5’-TGATCCTTCTGCAGGTTCACCTAC-3’). These primers are based on those of Amaral-Zettler et al. (2009)^45^ and Stoek et al. (2010)^46^, and are designed for use with Illumina platforms. The forward primer is a universal small-subunit primer, whereas the reverse primer favors eukaryotes but with mismatches can bind and amplify Bacteria and Archaea. In addition to deviations from the 16S protocol above with respect to primer construct sequences and PCR cycling parameters, we included a blocking primer that reduces amplification of vertebrate host DNA for host-associated samples, based on the strategy outlined by Vestheim et al. (2008)^47^. We note that the blocking primer is particularly useful for host-associated samples with a low biomass of non-host eukaryotic DNA.

The following primer constructs were used to integrate adapter sequences during amplification. For the barcoded forward primer, constructs included (5’ to 3’): the 5’ Illumina adapter (AATGATACGGCGACCACCGAGATCTACAC), a forward primer pad (TATCGCCGTT), a forward primer linker (CG), and the forward primer (Illumina_Euk_1391f) (GTACACACCGCCCGTC). For the reverse primer, constructs included (5’ to 3’): The reverse complement of 3’ Illumina adapter (CAAGCAGAAGACGGCATACGAGAT), a Golay barcode (12-bp variable sequence), a reverse primer pad (AGTCAGTCAG), a reverse primer linker (CA), and the reverse primer (806R [Apprill]) (TGATCCTTCTGCAGGTTCACCTAC). The construct for the blocking primer is as such and is formatted for ordering from IDT (Coralville, Iowa, USA): “GCCCGTCGCTACTACCGATTGG/ideoxyI//ideoxyI//ideoxyI//ideoxyI//ideoxyI/TTAGTGAG GCCCT/3SpC3/”.

Reaction mixtures without the blocking primer (i.e., those for non-vertebrate hosts or free-living sample types as defined by EMPO) were prepared as described for 16S. For reactions including the blocking primer, we combined 9 µL PCR-grade water, 10 µL master mix, 0.5 µL of each primer (10 µM), 4 µL of blocking primer (10 µM), and 1 µL of template DNA. The final concentration of the master mix in each 1X reaction was 0.8X, that of each primer was 0.2 µM, and that of the blocking primer was 1.6 µM. Without blocking primers, cycling parameters for a 384-well thermal cycler were as follows: 94°C for 3 min; 35 cycles of 94°C for 45 s, 57°C for 1 min, and 72°C for 90 s; and 72°C for 10 min. With blocking primers, cycling parameters for a 384-well thermal cycler were as follows: 94°C for 3 min; 35 cycles of 94°C for 45 s, 65°C for 15 s, 57°C for 30 s, and 72°C for 90 s; and 72°C for 10 min. Expected bands ranged between 210-310 bp.

For sequencing, the following primer constructs were used. Read 1 constructs (Euk_illumina_read1_seq_primer) included (5’ to 3’): a forward primer pad (TATCGCCGTT), a forward primer linker (CG), and the forward primer (1391f) (GTACACACCGCCCGTC). Read 2 constructs (Euk_illumina_read2_seq_primer) included (5’ to 3’): a reverse primer pad (AGTCAGTCAG), a reverse primer linker (CA), and the reverse primer (EukBr) (TGATCCTTCTGCAGGTTCACCTAC). The index primer construct (Euk_illumina_index_seq_primer) included (5’ to 3’): the reverse complement of the reverse primer (EukBr) (GTAGGTGAACCTGCAGAAGGATCA), the reverse complement of the reverse primer linker (TG), and the reverse complement of the reverse primer pad (CTGACTGACT).

The protocol for ITS is outlined fully in Smith et al. (2018)^48^. To target the fungal internal transcribed spacer (ITS1), we used the primers ITS1f (5’-CTTGGTCATTTAGAGGAAGTAA-3’) and ITS2 (5’-GCTGCGTTCTTCATCGATGC-3’). These primers are based on those of White et al. (1990) ^49^, and we note that primer ITS1f used here binds 38 bp upstream of ITS1 reported in that study.

The following primer constructs were used to integrate adapter sequences during amplification. For the barcoded forward primer, constructs included (5’ to 3’): the 5’ Illumina adapter (AATGATACGGCGACCACCGAGATCTACAC), a forward primer linker (GG), and the forward primer (ITS1f) (CTTGGTCATTTAGAGGAAGTAA). For the reverse primer, constructs included (5’ to 3’): The reverse complement of 3’ Illumina adapter (CAAGCAGAAGACGGCATACGAGAT), a Golay barcode (12-bp variable sequence), a reverse primer linker (CG), and the reverse primer (ITS2) (GCTGCGTTCTTCATCGATGC).

Reaction mixtures without the blocking primer were prepared as described for 16S. Cycling parameters for a 384-well thermal cycler were as follows: 94°C for 1 min; 35 cycles of 94°C for 30 s, 52°C for 30 s, and 68°C for 30 s; and 68°C for 10 min. Expected bands ranged between 250-600 bp^50, 51^.

For sequencing, the following primer constructs were used. Read 1 sequencing primer constructs included (5’ to 3’): a forward primer segment (TTGGTCATTTAGAGGAAGTAA), and a region extending into the amplicon (AAGTCGTAACAAGGTTTCC). Read 2 sequencing primer constructs included (5’ to 3’): a reverse primer segment (CGTTCTTCATCGATGC), and a region extending into the amplicon (VAGARCCAAGAGATC). The index sequencing primer construct included (5’ to 3’): the reverse complement of the region extending into the amplicon (TCTC), the reverse complement of the reverse primer (GCATCGATGAAGAACGCAGC), and the reverse complement of the linker (CG).

The protocol for generating bacterial full-length rRNA operon data is described in Karst et al. (2021)^36^. The method uses a unique molecular identifier (UMI) strategy to remove PCR errors and chimeras, resulting in a mean error rate of 0.0007% and a chimera rate of 0.02% of the final amplicon data. Briefly, the bacterial rRNA operons were targeted with an initial PCR using tailed versions of 27f (AGRGTTYGATYMTGGCTCAG)^52^ and 2490r (GACGGGCGGTGWGTRCA)^53^. The primer tails contained synthetic priming sites and 18-bp long patterned UMIs (NNNYRNNNYRNNNYRNNN). The PCR reaction (50-µL) contained 1-2 ng DNA template, 1U Platinum SuperFi DNA Polymerase High Fidelity (Thermo Fisher Scientific, Waltham, Massachusetts, USA) and a final concentration of 1× SuperFi buffer, 0.2mM of each dNTP, 500nM of each tailed 27f and tailed 2490r. The PCR program consisted of initial denaturation (3 min at 95°C) and two cycles of denaturation (30 s at 95°C), annealing (30 s at 55°C) and extension (6 min at 72°C). The PCR product was purified using a custom bead purification protocol ‘SPRI size selection protocol for >1.5–2 kb DNA fragments’ (Oxford Nanopore Technologies)’. The resulting product consists of uniquely tagged rRNA operon amplicons. The uniquely tagged rRNA operons were amplified in a second PCR, where the reaction (100-µL) contained 2U Platinum SuperFi DNA Polymerase High Fidelity (Thermo Fisher Scientific, Waltham, Massachusetts, USA) and a final concentration of 1X SuperFi buffer, 0.2 mM of each dNTP, 500 nM of each forward and reverse synthetic primer targeting the tailed primers from above. The PCR program consisted of initial denaturation (3 min at 95°C) and then 25-35 cycles of denaturation (15 s at 95°C), annealing (30 s at 60°C) and extension (6 min at 72°C) followed by final extension (5 min at 72°C). The PCR product was purified using the custom bead purification protocol above. Batches of 25 amplicon libraries were barcoded and sent for PacBio Sequel II library preparation and sequencing (Sequel II SMRT Cell 8M and 30 h collection time) at the DNA Sequencing Center at Brigham Young University. Circular consensus sequencing (CCS) reads were generated using CCS v.3.4.1 (https://github.com/PacificBiosciences/ccs) using default settings. UMI consensus sequences were generated using the longread_umi pipeline (https://github.com/SorenKarst/longread_umi) using the following command: longread_umi pacbio_pipeline -d ccs_reads.fq -o out_dir -m 3500 -M 6000 -s 60 -e 60 -f CAAGCAGAAGACGGCATACGAGAT -F AGRGTTYGATYMTGGCTCAG -r AATGATACGGCGACCACCGAGATC -R CGACATCGAGGTGCCAAAC -U ’0.75;1.5;2;0’ -c 2.

#### Amplicon data analysis

For multi-omics analyses including amplicon sequence data, we processed each dataset for comparison of beta-diversity. For all amplicon data except that for bacterial full-length rRNA amplicons, raw sequence data were converted from bcl to fastq, and then multiplexed files for each sequencing run uploaded as separate preparations to Qiita (study: 13114). For each sequencing run, data were then demultiplexed, trimmed to 150-bp, and denoised using Deblur^54^ to generate a feature-table of sub-operational taxonomic units (sOTUs) per sample.

For each 16S sequencing run, we placed denoised reads into the GreenGenes 13_8 phylogeny^55^ via fragment insertion using QIIME 2’s^22^ SATé-Enabled Phylogenetic Placement (SEPP)^56^ plugin, to produce a phylogeny for diversity analyses. To allow for phylogenetically-informed diversity analyses, reads not placed during SEPP (i.e., 513 sOTUs, 0.1% of all sOTUs) were removed from each feature-table. We then used QIIME 2’s *feature-table* plugin to merge feature-tables across sequencing runs, exclude singleton sOTUs, and rarefy the data to 5,000 reads per sample. Rarefaction depths for all amplicon analyses were chosen to best normalize sampling effort per sample while maintaining ≥75% of samples representative of the Earth’s environments, and also to maintain consistency with the analyses from EMP release 1^1^. We then used QIIME 2’s^22^ *diversity* plugin to estimate alpha-diversity (i.e., sOTU richness) and beta-diversity (i.e., unweighted UniFrac distances). The final feature-table for 16S beta-diversity analysis included 681 samples and 93,260 features. We performed a comparative analysis of the data including and excluding the reads not placed during SEPP, and note that that both alpha-diversity (i.e., sOTU richness) and beta-diversity (i.e., sample-sample RPCA distances) were highly correlated between datasets (Spearman *r* = 1.0) (Fig. S15). We thus proceeded with the SEPP-filtered dataset, and used phylogenetically-informed diversity metrics where applicable.

For 18S data, we used QIIME 2^22^ to first merge feature-tables across sequencing runs, and then classify taxonomy for each sOTU via pre-fitted machine-learning classifiers^57^ and the SILVA 138 reference database^58^. We then used QIIME 2’s^22^ *feature-table* plugin to exclude singleton sOTUs and samples with a total frequency <3,000 reads, and the *deicode*^23^ plugin to estimate beta-diversity for each dataset using robust Aitchison distances^23^. The final feature table for 18S beta-diversity analysis included 496 samples and 40,587 features.

For fungal ITS data, we used QIIME 2^22^ to first merge feature-tables across sequencing runs, and then classify taxonomy for each sOTU as above but using the UNITE 8 reference database^59^. We then used QIIME 2’s *feature-table* plugin to exclude singleton sOTUs and samples with a total frequency <500 reads, and the *deicode*^23^ plugin to estimate beta-diversity for each dataset using robust Aitchison distances^23^. The final feature table for fungal ITS beta-diversity analysis included 488 samples and 10,821 features.

For full-length rRNA operon data, per-sample fasta files were re-formatted for importing to QIIME 2 as *SampleData[Sequences]* (i.e., with each header as ‘>{sample_identifier}_{sequence_identifier}), concatenated into a single fasta file, and imported. We then used QIIME 2’s *vsearch* plugin^60^ to dereplicate sequences and then cluster them at 65% similarity (i.e., due to rapid evolution at bacterial ITS regions). The 65% OTU feature-table had 365 samples and 285 features. The concatenated fasta file and 65% OTU feature-table were uploaded to Qiita as a distinct preparations (study: 13114). We then used QIIME 2’s^22^ *feature-table* plugin to exclude singleton OTUs and samples with a total frequency <500 reads, and the *deicode*^23^ plugin to estimate beta-diversity for each dataset using robust Aitchison distances^23^. The final feature table for full-length rRNA operon beta-diversity analysis included 242 samples and 196 features.

#### Shotgun metagenomic sequencing

One round of DNA extraction was performed as above for shotgun metagenomic sequencing. Sequencing libraries were prepared using a high-throughput version of the HyperPlus library chemistry (Kapa Biosystems) miniaturized to approximately 1:10 reagent volume and optimized for nanoliter-scale liquid-handling robotics^61^. An exhaustive, step-by-step protocol and accompanying software can be found in Sanders et al. (2019)^61^. Briefly, DNA from each sample was transferred to a 384-well plate and quantified using the Quant-iT PicoGreen dsDNA Assay Kit, and then normalized to 5 ng in 3.5 µL of molecular-grade water using an Echo 550 acoustic liquid-handling robot (Labcyte, San Jose, CA, USA). For library preparation, reagents for each step (i.e., fragmentation, end repair and A-tailing, ligation, and PCR) were added in 1:10 the recommended volumes using a Mosquito HTS micropipetting robot (SPT Labtech, Tokyo, Japan). Fragmentation was performed at 37°C for 20 min and A-tailing at 65°C for 30 min.

Sequencing adapters and barcode indices were added in two steps^62^. First, the Mosquito HTS robot was used to add universal adapter “stub” adapters and ligase mix to the end-repaired DNA, and the ligation reaction performed for 20°C for 1 h. Adapter-ligated DNA was then cleaned-up using AMPure XP magnetic beads and a BlueCat purification robot (BlueCat Bio, Concord, Massachusetts, USA) by adding 7.5 µL magnetic bead solution to the total sample volume, washing twice with 70% EtOH, and resuspending in 7 µL molecular-grade water. Then, the Echo 550 robot was used to add individual i7 and i5 indices to adapter-ligated samples without repeating any barcodes, and by iterating the assignment of i7 to i5 indices such to minimize repeating unique i7:i5 pairs. Cleaned, adapter-ligated DNA was then amplified by adding 4.5 µL of each sample to 5.5 µL PCR master mix and running for 15 cycles, and then purified again using magnetic beads and the BlueCat robot. Each sample was eluted into 10 µL water, and then transferred to a 384-well plate using the Mosquito HTS robot. Each library was quantified using qPCR and then pooled to equal molar fractions using the Echo 550 robot. The final pool was sequenced at Illumina on a NovaSeq6000 using S2 flow cells and 2x150-bp chemistry (Illumina, San Diego, California, USA). To increase sequence coverage for certain samples, libraries were re-pooled and a second sequencing run performed as above.

#### Shotgun data analysis

Raw sequence data were converted from bcl to fastq and demultiplexed to produce per-sample fastq files. The mean sequencing depth was 7,580,347 ± 7.82 x 10^13^ reads per sample. We processed raw reads with Atropos (v1.1.24)^63^ to trim universal adapter sequences, poly-G tails introduced by the NovaSeq instrument (i.e., from use of two-color chemistry), and low-quality bases from reads. Atropos parameters included poly-G trimming (nextseq-trim=30), inclusion of ambiguous bases (match-read-wildcards), a maximum error rate for adapter matching (error-rate=0.1, default), removal of low-quality bases at 3’ and 5’ ends prior to adapter removal (quality-cutoff=15), a maximum error rate for adapter matching (insert-match-error-rate=0.2, default), discarding of short, trimmed reads (minimum-length=100), and discarding of paired reads if even one fails filtering (pair-filter=any). Trimmed reads were then mapped to the Web of Life database of microbial genomes^64^, using bowtie2^65^ in very-sensitive mode, to produce alignments that were used for taxonomic and exploratory functional analysis of microbial communities. Bowtie2 settings included maximum and minimum mismatch penalties (mp=[1,1]), a penalty for ambiguities (np=1; default), read and reference gap open- and extend penalties (rdg=[0,1], rfg=[0,1]), a minimum alignment score for an alignment to be considered valid (score-min=[L,0,-0.05]), a defined number of distinct, valid alignments (k=16), and the suppression of SAM records for unaligned reads, as well as SAM headers (no-unal, no-hd). The Web of Life database is particularly attractive as it includes a phylogeny that can be used for diversity analyses, and was curated to represent phylogenetic breadth of Bacteria and Archaea^64^, ideal for analyses across diverse environments. We compared mapping to the Web of Life to Rep200, a curated database of NCBI representative and reference microbial genomes (i.e., corresponding to RefSeq release 200, released May, 14, 2020), and found little difference across environments (Fig. S16). We therefore chose the Web of Life as it allows for phylogenetically-informed analyses.

For taxonomic analysis, we generated a feature-table of counts of operational genomic units (OGUs) for each sample using a reference-based approach. We chose this method vs. the *de novo* or reference-free approach, as the latter uses assembly/clustering to deconvolute short reads into larger sequence units; it allows for the direct observation of the actual organisms in the community, but alone does not allow for meaningful characterization of them^66^. Reference-based approaches use reference sequences from described organisms, which allow us to find the closest matches, using them to describe the taxa in a community^66^. This strategy is advantageous as results are not dependent on the samples included and it is less difficult because sequences can more easily be aligned to a reference vs. assembled into MAGS^67, 68^. Most importantly, it allows for comparisons of results across samples and studies, therefore representing a standardized method. Specifically, we used Woltka’s^69^ *classify* function, with per-genome alignments and default parameters. Woltka’s default normal mode is such that for one query sequence mapped to *k* genomes, each genome receives a count of 1/*k*. To permit examination of rare taxa across environments, no genomes were excluded. For diversity analyses, to best normalize sampling effort per sample while maintaining ≥75% of samples representative of the Earth’s environments, we rarefied the OGU feature-table to 6,550 reads per sample. The final feature-table for analyses of shotgun metagenomic taxonomic diversity included 612 samples and 8,692 OGUs.

For alpha-diversity, we quantified three metrics, in part to see which had the strongest correlations with microbially-related metabolite richness. We used the R package *geiger*^70^ to quantify weighted Faith’s PD for each sample following the method of Swensen^71^. We used QIIME 2’s *diversity* plugin^22^ to quantify richness and Faith’s PD (i.e., unweighted), as well as beta-diversity (i.e., using weighted UniFrac distance) between each pair of samples. We performed PERMANOVA on that distance matrix to test for significant differences in microbial community composition across the various levels of EMPO. We then used Principal Coordinates Analysis (PCoA) and EMPeror^25^ to visualize differences in microbial community composition among samples. We used *songbird*^24^ to identify sets of microbial taxa whose abundances were associated with certain environments, and parameterized our *songbird* model as above for our LC-MS/MS data. We then mapped the differential abundance results from *songbird* onto a phylogeny representing all microbial taxa using *empress*^72^ to visualize phylogenetic relationships related to log fold changes in abundance relative to specific environments.

For the functional analysis, we initially generated two sets of annotations for comparison of read mapping across environments. First, we generated a feature-table of counts of Gene Ontology (GO) Terms (i.e., for biological process, molecular function, and cellular compartment) for each sample using Woltka’s *collapse* function, inputting per-gene alignments and with default parameters for mapping to GO Terms through MetaCyc. For subsequent analysis, we used QIIME 2’s^22^ *feature-table* plugin to exclude singleton features and rarefy the data to 5,000 sequences per sample. The final feature-table included 517 samples and 3,776 features (i.e., GO terms). We also generated a feature-table of counts of KEGG^73–75^ Enzyme Code (EC) features (i.e., enzymes) for each sample using PRROMenade^76^. Trimmed, quality-controlled reads were mapped to the PRROMenade index of bacterial and viral protein domains via the IBM Functional Genomics Platform^77^ following Haiminen *et al.* (2021)^78^, searching for maximal exact matches with a length ≥11 amino acids, and retaining samples with ≥10,000 annotated reads (i.e., summed across R1 and R2 read files). Annotated read counts were pushed to leaf level nodes in the four-level EC hierarchy (e.g., EC 1.2.3.4). For diversity analysis, we used QIIME 2’s^22^ *feature-table* plugin to exclude singleton features and samples with fewer than 150,000 reads. The final feature-table included 616 samples (representing 18 environments) and 1,250 enzymes (i.e., KEGG ECs). We performed a comparative analysis comparing the Woltka GO-term analysis and the PRROMenade KEGG EC analysis, and found PRROMenade to more efficiently map reads across the majority of environments (Fig. S14). We therefore proceeded with our analysis of microbial functions using PRROMenade. With that table, we used QIIME 2’s *deicode*^23^ plugin to estimate beta-diversity for each dataset using robust Aitchison distances^23^ and EMPeror^25^ to visualize differences in microbial community composition among samples. We then performed PERMANOVA as above to test for significant differences in microbial functional composition across the various levels of EMPO.

#### Nestedness analysis of metabolomics data and shotgun metagenomic data for microbial taxa

As our analysis of turnover (replacement) of microbial taxa suggested a degree of nestedness (gain or loss of taxa promoting differences in richness) among environments in line with previous observations based on EMP 16S release 1^1^, we tested for nestedness in our shotgun metagenomics data for microbial taxa. We used the NODF statistic^79^ to quantify nestedness based on the degree to which less diverse communities are subsets of more diverse communities, which we quantified at each major taxonomic level from phylum to species^1^. We used the rarefied feature-table described above, and a null model (i.e., equiprobable rows, fixed columns) for assessing observed values of NODF, which we considered at each taxonomic level, and for all of the samples and each subset of the samples at EMPO 2^1^. To compute standardized effect sizes (SES) and *p*-values for significance, we used simulated results (*n* = 10,000 iterations) to find the expectation and variance of the NODF statistic under the null model. SES values were large (>90).

### MULTI-OMICS

#### Alpha-diversity correlations

Using the alpha-diversity metrics for LC-MS/MS (i.e., richness) and shotgun metagenomic taxonomic data (i.e., richness, unweighted Faith’s PD, and weighted Faith’s PD), we performed correlation analysis to better understand relationships therein. We used the function *multilevel* available in the R package *correlation*^80^ to perform Spearman correlations for each environment (i.e., based on EMPO 4), treating study (i.e., the variable representing distinct PI submissions of samples), and adjusting for multiple comparisons using the Benjamini-Hochberg correction.

#### Machine-learning analyses

To better understand community composition of microbes and metabolites across environments and specifically which features are predictive of certain habitats, we performed machine-learning. For analyses of LC-MS/MS and shotgun metagenomic taxonomic- and functional data, additional samples were filtered from the feature-tables noted previously such to exclude environments with relatively low sample representation (i.e., <9 samples). For the LC-MS/MS feature-table, we excluded samples in the four EMPO environments (i.e., “Animal corpus (non-saline)”, “Animal proximal gut (non-saline)”, “Soil (saline)”, and “Surface (saline)”). The final feature-table included 605 samples (representing 15 environments), and 6,588 microbially-related metabolites. For the shotgun metagenomic feature-table for taxonomic analysis, we excluded samples in four EMPO environments (i.e., “Animal corpus (non-saline)”, “Fungus corpus (non-saline)”, “Surface (saline)”, and “Subsurface (non-saline)”). The final feature-table included 598 samples (representing 15 environments), and 8,587 microbial taxa (i.e., Woltka OGUs). For the shotgun metagenomic feature-table for functional analysis, we used QIIME2’s^22^ *feature-table* plugin to excluded samples in three EMPO environments (i.e., “Animal corpus (non-saline)”, “Surface (saline)”, and “Subsurface (non-saline)”), exclude singleton features, and normalize the total count per sample to 10,000 sequences. The final feature-table included 706 samples (representing 16 environments) and 1,133 enzymes (i.e., KEGG ECs).

For each feature-table, we trained an auto-AI classifier^81^ with SHAP explanations^82^ and the hyper-tuned XGBoost method^83^ for predicting environments (based on EMPO 4). Each dataset was split into a training set (80%) and a testing set (20%), with similar environmental distributions in each iteration for the classification of samples. We evaluated the predictive performance of each classifier by quantifying accuracy statistics across 20 randomized iterations, and specifically by using resulting confusion matrices to quantify the overall and per-environment precision, recall, and F1 score. To identify the most important features contributing to the classification, we examined SHAP explanations, which we used to describe the impact of each feature for prediction. For features with an impact in at least one of 20 iterations examined, we assigned absolute ranks for each feature per-iteration, and then assigned final ranks based on the mean of absolute ranks across iterations. For the top twenty ranked features per feature-table, we visualized the environment for which each feature was impactful, as well as the direction of impact. Direction was determined by assessing differences in the mean relative abundances of the focal environment vs. all other environments combined. Positive impact indicates a feature was predictive of the focal environment when it was more abundant there vs. the other environments.

#### Metabolite–microbe co-occurrence analysis

To begin to explore co-occurrences between microbes and metabolites across environments, we implemented an approach that generates co-occurrence probabilities between all metabolite- and microbial features, clusters metabolites based on their co-occurrence with the microbial community, and highlights individual microbial features driving global patterns in metabolite distribution in this space. For co-occurrence analyses of LC-MS/MS metabolites and genomes profiled from shotgun metagenomic data, feature-tables were further filtered to retain only the 434 samples found in both datasets. For the LC-MS/MS feature-table of microbially-related secondary metabolites, we excluded 172 samples lacking shotgun metagenomics data, resulting in a final set of 6,501 microbially-related metabolites. For the shotgun metagenomics feature-table for taxonomy, we excluded 150 samples lacking LC-MS/MS data, resulting in a final set of 4,120 OGUs.

Specifically, we obtained co-occurrence probabilities and ordinated metabolites in microbial taxon space using *mmvec*, which uses the probabilities (i.e., log conditional probabilities, or co-occurrence strength) to predict metabolites based on microbial taxa from neural-network, compositionally-robust modeling^84^. The model was trained on 80% of the 434 samples, which were selected to balance environments (i.e., EMPO 4), and used the following parameters: epochs = 200, batch-size = 165, learning-rate = 1.0 x 10^-5^, summary-interval = 1, and with ‘equalize-biplot’. For training and testing, we filtered to retain only those features present in at least 10 samples (i.e., min-feature-count = 10), and restricted decomposition of the co-occurrence matrix to 10 principal components (PCs) (i.e., latent-dim = 10). The model predicting metabolite–microbe co-occurrences was more accurate than one representing a random baseline, with a pseudo-*Q*^2^ value of 0.18, indicating much reduced error during cross-validation.

To relate these metabolite-microbe co-occurrences to the distribution of metabolites across environments, we calculated the Spearman correlation between the loadings of metabolites on each co-occurrence PC vs. (i) log fold changes in metabolite abundances for each environment (i.e., from *songbird*), (ii) loadings for metabolites on the first three axes from the ordination corresponding to clustering of samples by environment (i.e., from RPCA), and (iii) a vector representing the global magnitude of metabolite importance across all three axes from that same ordination. To explicitly highlight metabolite-microbe co-occurrences specific to particular environments, we visualized the relationships between metabolite–microbe co-occurrences and (i) by considering the first three PCs of the co-occurrence ordination (i.e., from *mmvec*) and coloring metabolites by their log fold change values for a focal environment (e.g., Fig. 4b, Fig. S13). Then, focusing on the co-occurrence PC exhibiting the strongest correlation with log fold changes in metabolite abundances with respect to the focal environment, we manually selected one subset of metabolites highly abundant with respect to the focal environment but similar with respect to co-occurrences with microbes (i.e., high values on both axes, the focal group of metabolites), and one subset of metabolites lowly abundant with respect to the focal environment but similar with respect to co-occurrences with microbes (i.e., low values on both axes, the reference group of metabolites)^85^. Each select group of metabolites was chosen to represent a single pathway. Then, depending on the focal environment, we chose either the top 10 or top 10% of co-occurring microbes (i.e., based on co-occurrence strength) for each of the focal and reference groups of metabolites^83^. Finally, we visualized differences in the log-ratio of the focal group to the reference group between the focal environment and all other environments, separately for metabolites and microbes^83^.

#### Mantel correlations between datasets

To explore the relationships between sample–sample distances for any two datasets (e.g., LC-MS/MS vs. shotgun metagenomic for taxonomy), we used QIIME 2’s *diversity* plugin^22^ to perform Mantel tests on all pairings of the datasets using Spearman correlations. Input distance matrices are those described above for each dataset.

## Data availability

The mass spectrometry method and data (.RAW and .mzML) were deposited on the MassIVE public repository and are available under the dataset accession number <u>MSV000083475</u>. The processing files were also added to the deposition (updates/2019-08-21_lfnothias_7cc0af40/other/1908_EMPv2_INN/). GNPS molecular networking job is available at https://gnps.ucsd.edu/ProteoSAFe/status.jsp?task=929ce9411f684cf8abd009670b293a33 and was also performed in analogue mode https://gnps.ucsd.edu/ProteoSAFe/status.jsp?task=fafdbfc058184c2b8c87968a7c56d7aa. The *DEREPLICATOR* jobs can be accessed here: https://gnps.ucsd.edu/ProteoSAFe/status.jsp?task=ee40831bcc314bda928886964d853a52 and https://gnps.ucsd.edu/ProteoSAFe/status.jsp?task=1fafd4d4fe7e47dd9dd0b3d8bb0e6606. The SIRIUS results are available on the GitHub repository [‘emp/data/metabolomics/FBMN/SIRIUS’]. The notebooks for metabolomics data preparation and microbially-related molecules establishment are available on this repository (https://github.com/lfnothias/emp_metabolomic). Amplicon and shotgun metagenomic sequence data are submitted to the European Nucleotide Archive under Project: PRJEB42019 (https://www.ebi.ac.uk/ena/browser/view/PRJEB42019). Raw and demultiplexed amplicon and shotgun sequence data, the feature-table for full-length rRNA operon analysis, feature-tables for LC-MS/MS classical molecular networking and feature-based molecular networking, and the feature-table for GC-MS molecular networking data are available for download and analysis through Qiita at https://www.qiita.ucsd.edu (study: 13114).

## Code availability

We provide complete protocols for laboratory- and computational workflows for both metagenomics and metabolomics data for use by the broader community, available on GitHub (https://github.com/biocore/emp/blob/master/methods/methods_release2.md).

