## Supplementary material for "Multi-omics profiling of Earth’s biomes reveals patterns of diversity and co-occurrence in microbial and metabolite composition across environments"

[15]Bioinformatics and Systems Biology Program, Jacobs School of Engineering, University of California San Diego, La Jolla, California, USA
[16]Center for Computational Biology, Flatiron Institute, Simons Foundation, USA
[17]Chair for Bioinformatics, Friedrich Schiller University, Jena, Germany
[18]College of Pharmacy and Integrated Research Institute for Drug Development, Dongguk University, Gyeonggi-do, Korea
[19]Department of Computer Science, University of Antwerp, Antwerp, Belgium
[20]IBM Research Europe - Daresbury, UK
[21]IBM Research, Almaden Research Center, San Jose, California, USA
[22]Department of Virus and Microbiological Special Diagnostics, Statens Serum Institute, Copenhagen, Denmark
[23]Department of Chemistry and Bioscience, Aalborg University, Aalborg, Denmark
[24]Centre for Marine Science and Innovation, School of Biological, Earth and Environmental Science, The University of New South Wales, Sydney, 2052, Australia
[25]Interfaculty Institute of Microbiology and Infection Medicine, University of Tübingen, Tübingen, Baden-Württemberg 72076, Germany
[26]Department of Microbiology and Molecular Genetics, Michigan State University, East Lansing, Michigan, USA
[27]Department of Computer Science and Engineering, Jacobs School of Engineering, University of California San Diego, La Jolla, California, USA
[#]Co-first author
*Corresponding author
[δ]A list of authors and their affiliations appears at the end of the main text

**SUPPLEMENTARY TABLES**

**Table S1 |** Study metadata, including study identifiers, principal investigator, title, sampling method, number of samples, environmental package, EMPO level 3 (version 1), and EMPO level 4 (version 2) for each study.

| emp500_study_id | emp500_principal_investigator | emp500_title | sampling_method | number_of_samples | environmental_package | empo_3 | empo_v2_4 |
|---|---|---|---|---|---|---|---|
| 1 | Kshtrika | Biota oil | bulk | 14 | misc environment | Subsurface (non-saline) | Subsurface (non-saline) |
| 2 | Berry | Postglacial pond sediment profiles | bulk | 20 | sediment | Sediment (non-saline) | Sediment (non-saline) |
| 3 | MacRae-Crerar | Mongolia soils (NSF PIRE) | bulk | 12 | soil | Soil (non-saline) | Soil (non-saline) |
| 5 | Myrold | Tree-associated soils | bulk | 12 | soil | Soil (non-saline) | Soil (non-saline) |
| 9 | Zaneveld | Scleractinian corals | fractionated | 22 | host-associated, water | Animal secretion, Water (saline) | Animal secretion, Water (saline) |
| 18 | Thomas | Ocean sponges | fractionated | 57 | host-associated | Animal corpus | Animal corpus (saline) |
| 19 | Thomas | Australian algae | swab | 64 | host-associated | Plant surface | Plant surface (saline) |
| 21 | McMahon | Bioreactors for wastewater and anammox | bulk | 17 | wastewater/sludge | Water (non-saline) | Water (non-saline) |
| 23 | Shade | Centralia soil | bulk | 10 | soil | Soil (non-saline) | Soil (non-saline) |
| 26 | Stewart | Marine oxygen minimum zone water | filter | 8 | water | Water (saline) | Water (saline) |
| 27 | King | Puhimau Hawai'i volcanic soil | bulk | 21 | soil | Soil (non-saline) | Soil (non-saline) |
| 33 | Mayer | West Texas playa | bulk | 7 | sediment | Sediment (non-saline) | Sediment (non-saline) |
| 34 | Mayer | Animas watershed | bulk | 10 | sediment | Sediment (non-saline) | Sediment (non-saline) |
| 36 | Stegen | Permafrost soil | bulk | 15 | soil | Soil (non-saline) | Soil (non-saline) |
| 37 | Stegen | Hyporheic sediment | bulk | 10 | sediment | Sediment (non-saline) | Sediment (non-saline) |
| 38 | Stegen | Active layer soil | bulk | 18 | soil | Soil (non-saline) | Soil (non-saline) |
| 39 | Smith | Palmyra algae | bulk or swab | 8 | host-associated | Plant surface | Plant surface (saline) |
| 40 | Metcalf | Mouse decomposition soil | bulk | 14 | soil | Soil (non-saline) | Soil (non-saline) |
| 42 | Palenik | San Diego seawater | filter | 20 | water | Water (saline) | Water (saline) |
| 43 | Jensen | Ocean sediments | bulk | 19 | sediment | Sediment (saline) | Sediment (saline) |
| 45 | Roy-Chowdhury | Spanish hypersaline sandy soils | bulk | 24 | soil | Soil (non-saline) | Soil (non-saline) |
| 46 | Makhalanyane | Namibia gravel plain gradient soils | bulk | 10 | soil | Soil (non-saline) | Soil (non-saline) |
| 47 | Makhalanyane | Namibia dune aridity gradient soils | bulk | 9 | soil | Soil (non-saline) | Soil (non-saline) |
| 50 | Girguis | Whale feces | bulk | 12 | host-associated | Animal distal gut | Animal distal gut (saline) |
| 51 | Song | Captive terrestrial mammal feces | bulk | 20 | host-associated | Animal distal gut | Animal distal gut (non-saline) |
| 52 | Song | Captive herp feces | bulk | 15 | host-associated | Animal distal gut | Animal distal gut (non-saline) |
| 53 | Song | Captive bird feces | bulk | 15 | host-associated | Animal distal gut | Animal distal gut (non-saline) |
| 54 | Sandin | Fish gut contents | bulk | 15 | host-associated | Animal proximal gut | Animal proximal gut (saline) |
| 56 | Schmidt | Peru and Nepal alpine soils | bulk | 26 | soil | Soil (non-saline) | Soil (non-saline) |
| 58 | Tucker | Coal and coal-associated water | filter | 16 | misc environment | Subsurface (non-saline) | Subsurface (non-saline) |
| 59 | Myrold | Oregon soils | bulk | 22 | soil | Soil (non-saline) | Soil (non-saline) |
| 62 | Pinto | Costa Rican leaf-cutter ant soils | bulk | 24 | soil | Soil (non-saline) | Soil (non-saline) |
| 63 | Pinto | Costa Rican leaf-cutter ant and bee material | bulk | 28 | host-associated | Plant detritus | Plant detritus (non-saline) |
| 65 | Angenent | Rumen animals and bioreactor | bulk | 9 | host-associated | Animal proximal gut | Animal proximal gut (non-saline) |
| 68 | Bittleston | Pitcher plant fluids | bulk | 15 | plant-associated | Plant surface | Plant surface (non-saline) |
| 71 | Seedorf | Singapore plant-associated soils | bulk | 14 | soil | Soil (non-saline) | Soil (non-saline) |
| 72 | Distel | Shipworm tissue and wood | fractionated | 15 | host-associated | Animal corpus, Animal proximal gut, Surface (non-saline) | Animal corpus, Animal proximal gut, Surface (non-saline) |
| 74 | Bowen | Massachusetts salt marsh sediments | bulk | 16 | sediment | Sediment (saline) | Sediment (saline) |
| 75 | Song | Captive aquatic mammal feces | bulk | 5 | host-associated | Animal distal gut | Animal distal gut (non-saline) |
| 76 | Minich | Livestock feces | bulk | 6 | host-associated | Animal distal gut | Animal distal gut (non-saline) |
| 77 | Tait | English Channel water | filter | 9 | water | Water (saline) | Water (saline) |
| 78 | Tait | English Channel sediment | bulk | 3 | sediment | Sediment (saline) | Sediment (saline) |
| 81 | Metcalf | Mouse decomposition skin | bulk | 3 | host-associated | Animal corpus | Animal corpus (non-saline) |
| 82 | Uren | Michigan forest lichen | bulk | 12 | plant-associated | Fungus corpus | Fungus corpus (non-saline) |
| 84 | Rohwer | Franz Josef Land glacier soils and water | bulk | 15 | soil, water | Soil (non-saline), Water (non-saline) | Soil (non-saline), Water (non-saline) |
| 85 | Rohwer | Iberia and Guatemala deep-sea sediments | bulk | 27 | sediment | Sediment (saline) | Sediment (saline) |
| 86 | Rohwer | Mexico microbial mats | bulk | 3 | microbial mat/biofilm, sediment | Surface (saline), Sediment (saline) | Surface (saline), Sediment (saline) |
| 88 | Mousseau | Chernobyl vole feces | bulk | 112 | host-associated | Animal distal gut | Animal distal gut (non-saline) |

Note: Scientific justification and metadata columns for each study are included in the version of the table here and will be included with the final Supplementary Material, but were excluded here due to space limitations.

**Table S2 |** Summary of data generated across eight layers for each sample. Read counts are shown for sequence data.

TABLE TOO LARGE TO DISPLAY – WILL BE INCLUDED AS A SEPARATE FILE

**Table S3 |** The top 10 ranked microbially-related metabolites for each environment, described using the Earth Microbiome Project Ontology (EMPO, version 2, level 4; Fig. 1). Data are from Songbird multinomial regression (model: *composition* = EMPO level 4; pseudo-$Q^2$ = 0.21). Metabolites in bold font highlighted gray are those also identified to be strongly associated with RPCA axes (Fig. 3c, Table S4), and those in bold font highlighted black also strongly separated environments in machine-learning analysis (Table S6).

| Environment | Feature rank | Feature ID | Molecular formula | Pathway | Superclass | Class |
|---|---|---|---|---|---|---|
| Animal corpus (saline) | 1 | 23235 | C27H40O2 | Terpenoids | Sesterterpenoids | Not annotated |
| Animal corpus (saline) | 2 | 41006 | Not annotated | Not annotated | Not annotated | Not annotated |
| Animal corpus (saline) | 3 | 28910 | C27H38 | Terpenoids | Steroids | Cholestane steroids |
| Animal corpus (saline) | 4 | 23136 | C25H32O3 | Terpenoids | Cyclic polyketides | Not annotated |
| Animal corpus (saline) | 5 | 21140 | C29H46O4 | Terpenoids | Triterpenoids | Not annotated |
| Animal corpus (saline) | 6 | 3559 | Not annotated | Not annotated | Not annotated | Not annotated |
| Animal corpus (saline) | 7 | 23517 | Not annotated | Not annotated | Not annotated | Not annotated |
| Animal corpus (saline) | 8 | 41001 | Not annotated | Not annotated | Not annotated | Not annotated |
| Animal corpus (saline) | 9 | 3955 | Not annotated | Not annotated | Not annotated | Not annotated |
| Animal corpus (saline) | 10 | 39515 | Not annotated | Not annotated | Not annotated | Not annotated |
| **Animal distal gut (non-saline)** | **1** | **25552** | **C24H38O4** | **Terpenoids** | **Steroids** | **Cholane steroids** |
| Animal distal gut (non-saline) | 2 | 7792 | C24H38O4 | Terpenoids | Steroids | Cholane steroids |
| Animal distal gut (non-saline) | 3 | 25862 | C27H42O7 | Terpenoids | Steroids | Ecdysteroids |
| Animal distal gut (non-saline) | 4 | 51039 | C27H44O3 | Terpenoids | Steroids | Cholestane steroids |
| Animal distal gut (non-saline) | 5 | 2081 | C15H10O4 | Shikimates and Phenylpropanoids | Isoflavonoids | Isoflavones |
| Animal distal gut (non-saline) | 6 | 1236 | C18H34O5 | Fatty acids | Octadecanoids | Other Octadecanoids |
| Animal distal gut (non-saline) | 7 | 50823 | C26H45NO6S | Terpenoids | Steroids | Cholane steroids |
| **Animal distal gut (non-saline)** | **8** | **22299** | **C29H46O2** | **Terpenoids** | **Meroterpenoids** | **Prenyl quinone meroterpenoids** |
| Animal distal gut (non-saline) | 9 | 2826 | C19H34O4 | Fatty acids | Octadecanoids | Other Octadecanoids |
| Animal distal gut (non-saline) | 10 | 12107 | C24H41NO3 | Terpenoids | Steroids | Cholane steroids |
| Animal distal gut (saline) | 1 | 29157 | Not annotated | Not annotated | Not annotated | Not annotated |
| Animal distal gut (saline) | 2 | 38117 | C40H52O4 | Terpenoids | Carotenoids (C40) | Carotenoids (C40, β-β) |
| Animal distal gut (saline) | 3 | 7942 | C17H24N2O4 | Alkaloids | Ornithine alkaloids | Not annotated |
| Animal distal gut (saline) | 4 | 53059 | C15H18BrO | Terpenoids | Sesquiterpenoids | Not annotated |
| Animal distal gut (saline) | 5 | 30947 | C7H9NO2 | Shikimates and Phenylpropanoids | Not annotated | Shikimic acids and derivatives |
| Animal distal gut (saline) | 6 | 20600 | Not annotated | Not annotated | Not annotated | Not annotated |
| Animal distal gut (saline) | 7 | 50368 | C27H31N3O3 | Alkaloids | Tryptophan alkaloids | Not annotated |
| Animal distal gut (saline) | 8 | 52381 | Not annotated | Not annotated | Not annotated | Not annotated |
| Animal distal gut (saline) | 9 | 50979 | C24H36O2S | Terpenoids | Steroids | Cholane steroids |
| Animal distal gut (saline) | 10 | 12107 | C24H41NO3 | Terpenoids | Steroids | Cholane steroids |
| Animal proximal gut (saline) | 1 | 46142 | C19H39NO2S | Fatty acids | Fatty Acids and Conjugates | Not annotated |
| Animal proximal gut (saline) | 2 | 50500 | Not annotated | Not annotated | Not annotated | Not annotated |
| Animal proximal gut (saline) | 3 | 46161 | C19H39NO2S | Fatty acids | Fatty Acids and Conjugates | Not annotated |
| Animal proximal gut (saline) | 4 | 50865 | Not annotated | Not annotated | Not annotated | Not annotated |
| Animal proximal gut (saline) | 5 | 8409 | C22H35NO2 | Fatty acids | Fatty amides | N-acyl ethanolamines (endocannabinoids) |
| Animal proximal gut (saline) | 6 | 15710 | Not annotated | Not annotated | Not annotated | Not annotated |
| Animal proximal gut (saline) | 7 | 19189 | Not annotated | Not annotated | Not annotated | Not annotated |
| Animal proximal gut (saline) | 8 | 44429 | Not annotated | Not annotated | Not annotated | Not annotated |
| Animal proximal gut (saline) | 9 | 47998 | Not annotated | Not annotated | Not annotated | Not annotated |
| Animal proximal gut (saline) | 10 | 49293 | C18H36N4O4 | Amino acids and Peptides | Small peptides | Not annotated |
| Animal secretion (saline) | 1 | 24213 | C25H40O3 | Fatty acids | Glycerolipids | Monoacylglycerols |
| Animal secretion (saline) | 2 | 46948 | C20H35N5O13 | Carbohydrates | Polyols | Amino cyclitols |
| Animal secretion (saline) | 3 | 24185 | C20H33ClO4 | Polyketides | Not annotated | Not annotated |
| Animal secretion (saline) | 4 | 10248 | C24H41NO5 | Not annotated | Steroids | Not annotated |
| Animal secretion (saline) | 5 | 16936 | C33H52O13 | Fatty acids | Glycerolipids | Glycosyldiacylglycerols |
| Animal secretion (saline) | 6 | 24147 | C25H40O3 | Terpenoids | Steroids | Cholane steroids |
| Animal secretion (saline) | 7 | 40413 | C22H46NO8P | Fatty acids | Glycerophospholipids | Glycerophosphates |
| Animal secretion (saline) | 8 | 44580 | C11H21N3O | Alkaloids | Small peptides | Dipeptides |
| Animal secretion (saline) | 9 | 22091 | C28H50O9 | Polyketides | Polyethers | Not annotated |
| Animal secretion (saline) | 10 | 46073 | C10H17N | Terpenoids | Monoterpenoids | Not annotated |
| Fungus corpus (non-saline) | 1 | 20438 | C18H10O7 | Shikimates and Phenylpropanoids | Flavonoids | Flavonols |
| Fungus corpus (non-saline) | 2 | 16650 | C18H12O8 | Shikimates and Phenylpropanoids | Flavonoids | Flavonols |
| Fungus corpus (non-saline) | 3 | 16117 | C25H43NO9 | Fatty acids | Glycerolipids | Glycosyldiacylglycerols |
| Fungus corpus (non-saline) | 4 | 16952 | Not annotated | Not annotated | Not annotated | Not annotated |
| Fungus corpus (non-saline) | 5 | 16752 | C18H30Cl2O2 | Fatty acids | Fatty Acids and Conjugates | Not annotated |
| Fungus corpus (non-saline) | 6 | 16681 | C18H12O6 | Shikimates and Phenylpropanoids | Flavonoids | Not annotated |
| Fungus corpus (non-saline) | 7 | 16632 | C17H10O6 | Polyketides | Xanthones | Methyl xanthones |
| Fungus corpus (non-saline) | 8 | 16615 | C18H12O7 | Shikimates and Phenylpropanoids | Not annotated | Not annotated |
| Fungus corpus (non-saline) | 9 | 16743 | C18H12O8 | Shikimates and Phenylpropanoids | Flavonoids | Flavonols |
| Fungus corpus (non-saline) | 10 | 15767 | C20H34O2 | Fatty acids | Fatty Acids and Conjugates | Not annotated |
| Plant detritus (non-saline) | 1 | 24670 | C33H40O4 | Polyketides | Meroterpenoids | Not annotated |
| Plant detritus (non-saline) | 2 | 24678 | C13H16BNO7 | Amino acids and Peptides | Not annotated | Not annotated |
| Plant detritus (non-saline) | 3 | 24692 | C33H48O4 | Polyketides | Meroterpenoids | Not annotated |
| Plant detritus (non-saline) | 4 | 7395 | C22H30O3 | Terpenoids | Meroterpenoids | Cannabinoids |
| Plant detritus (non-saline) | 5 | 24684 | Not annotated | Not annotated | Not annotated | Not annotated |
| Plant detritus (non-saline) | 6 | 24677 | C24H26O4 | Polyketides | Phloroglucinols | Acyl phloroglucinols |
| Plant detritus (non-saline) | 7 | 30111 | C27H42O6 | Terpenoids | Steroids | Ecdysteroids |
| Plant detritus (non-saline) | 8 | 24866 | C19H18O5 | Polyketides | Phloroglucinols | Not annotated |
| Plant detritus (non-saline) | 9 | 26285 | C28H42 | Terpenoids | Steroids | Vitamin D2 and derivatives |
| Plant detritus (non-saline) | 10 | 31735 | C24H39N5O8 | Amino acids and Peptides | Small peptides | Tripeptides |
| Plant surface (non-saline) | 1 | 11326 | C24H41NO4 | Terpenoids | Steroids | Cholane steroids |
| Plant surface (non-saline) | 2 | 36400 | C32H54O4 | Polyketides | Meroterpenoids | Not annotated |
| Plant surface (non-saline) | 3 | 20775 | C20H41NO4 | Fatty acids | Not annotated | Other Octadecanoids |
| **Plant surface (non-saline)** | **4** | **4949** | **C13H10O** | **Shikimates and Phenylpropanoids** | **Flavonoids** | **Chalcones** |
| **Plant surface (non-saline)** | **5** | **30751** | **C9H15NO4** | **Polyketides** | **Cyclic polyketides** | **2-pyrone derivatives** |
| Plant surface (non-saline) | 6 | 5118 | C21H24O3 | Terpenoids | Diterpenoids | Tetracyclic diterpenoids |
| Plant surface (non-saline) | 7 | 20098 | C7H8NO4 | Amino acids and Peptides | Small peptides | Aminoacids |
| Plant surface (non-saline) | 8 | 45169 | C26H41NO8 | Alkaloids | Not annotated | Not annotated |
| Plant surface (non-saline) | 9 | 38866 | Not annotated | Not annotated | Not annotated | Not annotated |
| Plant surface (non-saline) | 10 | 21815 | C30H50N6O4 | Amino acids and Peptides | Oligopeptides | Lipopeptides |

**Table S3 |** (continued)

| Environment | Feature rank | Feature ID | Molecular formula | Pathway | Superclass | Class |
|---|---|---|---|---|---|---|
| Plant surface (saline) | 1 | 55614 | C8H13N3O | Alkaloids | Nicotinic acid alkaloids | Pyridine alkaloids |
| Plant surface (saline) | 2 | 55975 | C22H41N3O8 | Polyketides | Not annotated | Not annotated |
| Plant surface (saline) | 3 | 56485 | Not annotated | Not annotated | Not annotated | Not annotated |
| Plant surface (saline) | 4 | 56324 | C21H37N3O7 | Polyketides | Not annotated | Not annotated |
| Plant surface (saline) | 5 | 34407 | C10H19N3O2 | Alkaloids | Not annotated | Primary amides |
| Plant surface (saline) | 6 | 56318 | C10H17N3O | Alkaloids | Monoterpenoids | Pyridine alkaloids |
| Plant surface (saline) | 7 | 34652 | C10H19N3O2 | Alkaloids | Not annotated | Primary amides |
| Plant surface (saline) | 8 | 56548 | C23H40O8 | Fatty acids | Linear polyketides | Open-chain polyketides |
| Plant surface (saline) | 9 | 56014 | C22H39N3O7 | Amino acids and Peptides | Not annotated | Not annotated |
| Plant surface (saline) | 10 | 56496 | C21H41N3O7 | Polyketides | Linear polyketides | Open-chain polyketides |
| Sediment (non-saline) | 1 | 15024 | C13H24N6O7 | Carbohydrates | Not annotated | Not annotated |
| Sediment (non-saline) | 2 | 4702 | C9H16O5 | Fatty acids | Glycerolipids | Not annotated |
| Sediment (non-saline) | 3 | 28221 | C36H54O8 | Terpenoids | Diterpenoids | Tigliane diterpenoids |
| Sediment (non-saline) | 4 | 8030 | C20H30 | Terpenoids | Steroids | Not annotated |
| Sediment (non-saline) | 5 | 21379 | C25H26O5 | Shikimates and Phenylpropanoids | Coumarins | Not annotated |
| Sediment (non-saline) | 6 | 5882 | C9H6O2 | Shikimates and Phenylpropanoids | Phenylpropanoids (C6-C3) | Cinnamic acids and derivatives |
| Sediment (non-saline) | 7 | 1721 | C27H38O5 | Terpenoids | Not annotated | Not annotated |
| Sediment (non-saline) | 8 | 11099 | C19H34O5 | Fatty acids | Linear polyketides | Open-chain polyketides |
| Sediment (non-saline) | 9 | 25351 | Not annotated | Not annotated | Not annotated | Not annotated |
| Sediment (non-saline) | 10 | 15086 | C12H18O6 | Polyketides | Not annotated | Not annotated |
| Sediment (saline) | 1 | 12617 | C27H47NO13 | Fatty acids | Not annotated | Not annotated |
| **Sediment (saline)** | **2** | **42202** | **Not annotated** | **Not annotated** | **Not annotated** | **Not annotated** |
| Sediment (saline) | 3 | 10361 | C42H56O5 | Terpenoids | Carotenoids (C40) | Carotenoids (C40, β-β) |
| Sediment (saline) | 4 | 15118 | C19H35NO9 | Fatty acids | Not annotated | Not annotated |
| Sediment (saline) | 5 | 46984 | C21H34O3 | Terpenoids | Steroids | Cholane steroids |
| Sediment (saline) | 6 | 33113 | C32H57NO9 | Polyketides | Linear polyketides | Open-chain polyketides |
| Sediment (saline) | 7 | 12537 | C40H58O5 | Terpenoids | Carotenoids (C40) | Carotenoids (C40, β-β) |
| Sediment (saline) | 8 | 12862 | C21H34O3 | Fatty acids | Eicosanoids | Not annotated |
| Sediment (saline) | 9 | 12623 | C28H49NO13 | Fatty acids | Not annotated | Not annotated |
| Sediment (saline) | 10 | 12750 | Not annotated | Not annotated | Not annotated | Not annotated |
| Soil (non-saline) | 1 | 2102 | C18H37NO2 | Fatty acids | Fatty Acids and Conjugates | Unsaturated fatty acids |
| Soil (non-saline) | 2 | 10177 | C32H52N8O5 | Amino acids and Peptides | Oligopeptides | Linear peptides |
| Soil (non-saline) | 3 | 2073 | C17H19NO2 | Alkaloids | Not annotated | Not annotated |
| Soil (non-saline) | 4 | 2217 | C13H21ClN2O3 | Fatty acids | Not annotated | Not annotated |
| Soil (non-saline) | 5 | 2161 | C32H51N5O5 | Amino acids and Peptides | Oligopeptides | Cyclic peptides |
| Soil (non-saline) | 6 | 7661 | C20H39NO | Fatty acids | Fatty amides | Primary amides |
| Soil (non-saline) | 7 | 2135 | C18H18N2O | Alkaloids | Tryptophan alkaloids | Simple indole alkaloids |
| Soil (non-saline) | 8 | 2288 | C11H12N2O2 | Alkaloids | Tryptophan alkaloids | Simple indole alkaloids |
| Soil (non-saline) | 9 | 7420 | C13H19FOS2 | Terpenoids | Monoterpenoids | Not annotated |
| Soil (non-saline) | 10 | 5153 | C20H22O4 | Shikimates and Phenylpropanoids | Lignans | Dibenzylbutane lignans |
| Subsurface (non-saline) | 1 | 14373 | C14H19NO | Terpenoids | Sesquiterpenoids | Not annotated |
| Subsurface (non-saline) | 2 | 14208 | C8H12O4 | Fatty acids | Fatty Acids and Conjugates | Dicarboxylic acids |
| Subsurface (non-saline) | 3 | 42569 | Not annotated | Not annotated | Not annotated | Not annotated |
| Subsurface (non-saline) | 4 | 17757 | C20H20O | Terpenoids | Steroids | Not annotated |
| Subsurface (non-saline) | 5 | 6222 | C18H22 | Terpenoids | Steroids | Estrane steroids |
| Subsurface (non-saline) | 6 | 27023 | C17H32O5 | Fatty acids | Glycerolipids | Monoacylglycerols |
| Subsurface (non-saline) | 7 | 14337 | Not annotated | Not annotated | Not annotated | Not annotated |
| Subsurface (non-saline) | 8 | 15209 | C16H20O2S | Terpenoids | Steroids | Not annotated |
| Subsurface (non-saline) | 9 | 2225 | C14H21NO | Fatty acids | Fatty amides | Phenylalanine-derived alkaloids |
| Subsurface (non-saline) | 10 | 9506 | C17H18O | Terpenoids | Not annotated | Not annotated |
| Water (non-saline) | 1 | 14668 | C30H62O16 | Carbohydrates | Glycerolipids | Not annotated |
| Water (non-saline) | 2 | 14675 | C18H33N7O5 | Alkaloids | Pseudoalkaloids (transamidation) | Not annotated |
| Water (non-saline) | 3 | 14677 | C32H66O17 | Carbohydrates | Glycerolipids | Not annotated |
| Water (non-saline) | 4 | 14832 | C19H31BN6O4 | Alkaloids | Not annotated | Not annotated |
| Water (non-saline) | 5 | 14725 | C18H38O10 | Fatty acids | Glycerolipids | Diacylglycerols |
| Water (non-saline) | 6 | 14693 | C22H46O12 | Fatty acids | Glycerolipids | Diacylglycerols |
| Water (non-saline) | 7 | 14821 | C14H28O7 | Fatty acids | Not annotated | Not annotated |
| **Water (non-saline)** | **8** | **14665** | **C28H58O15** | **Carbohydrates** | **Glycerolipids** | **Not annotated** |
| Water (non-saline) | 9 | 15877 | Not annotated | Not annotated | Not annotated | Not annotated |
| Water (non-saline) | 10 | 14797 | Not annotated | Not annotated | Not annotated | Not annotated |
| Water (saline) | 1 | 14668 | C30H62O16 | Carbohydrates | Glycerolipids | Not annotated |
| Water (saline) | 2 | 14677 | C32H66O17 | Carbohydrates | Glycerolipids | Not annotated |
| Water (saline) | 3 | 14832 | C19H31BN6O4 | Alkaloids | Not annotated | Not annotated |
| Water (saline) | 4 | 14665 | C28H58O15 | Carbohydrates | Glycerolipids | Not annotated |
| Water (saline) | 5 | 14693 | C22H46O12 | Fatty acids | Glycerolipids | Diacylglycerols |
| **Water (saline)** | **6** | **14675** | **C18H33N7O5** | **Alkaloids** | **Pseudoalkaloids (transamidation)** | **Not annotated** |
| Water (saline) | 7 | 15877 | Not annotated | Not annotated | Not annotated | Not annotated |
| Water (saline) | 8 | 14785 | Not annotated | Not annotated | Not annotated | Not annotated |
| Water (saline) | 9 | 14844 | C26H55NO13 | Carbohydrates | Aminosugars and aminoglycosides | Not annotated |
| Water (saline) | 10 | 14724 | C24H50O13 | Fatty acids | Glycerolipids | Diacylglycerols |

**Table S4 |** The top 25 microbially-related metabolites across all environments, ranked based on association with RPCA ordination axes (i.e., global magnitude across all axes). Metabolites in bold font highlighted gray are those also identified to be strongly associated with particular environments from analysis of differential abundance (Table S2).

| Feature ID | Molecular formula | Pathway | Superclass | Class |
|---|---|---|---|---|
| 362 | C17H24O3 | Terpenoids | Sesquiterpenoids | Not annotated |
| 1630 | C9H8O2 | Shikimates and Phenylpropanoids | Phenylpropanoids (C6-C3) | Cinnamic acids and derivatives |
| 1632 | C9H8O2 | Shikimates and Phenylpropanoids | Phenolic acids (C6-C1) | Simple phenolic acids |
| 1856 | C22H41NO | Fatty acids | Fatty amides | Primary amides |
| 3394 | C28H38 | Terpenoids | Steroids | Vitamin D2 and derivatives |
| 3830 | C28H38 | Terpenoids | Steroids | Vitamin D2 and derivatives |
| 4561 | C9H6O2 | Shikimates and Phenylpropanoids | Phenylpropanoids (C6-C3) | Cinnamic acids and derivatives |
| **4949** | **C13H10O** | **Shikimates and Phenylpropanoids** | **Flavonoids** | **Chalcones** |
| 6974 | C14H24O5 | Terpenoids | Not annotated | Not annotated |
| 8263 | Not annotated | Not annotated | Not annotated | Not annotated |
| 8273 | Not annotated | Not annotated | Not annotated | Not annotated |
| 12599 | C10H10O2 | Shikimates and Phenylpropanoids | Phenylpropanoids (C6-C3) | Cinnamic acids and derivatives |
| 13892 | C8H18O5 | Fatty acids | Glycerolipids | Diacylglycerols |
| **14665** | **C28H58O15** | **Carbohydrates** | **Glycerolipids** | **Not annotated** |
| 14675 | C18H33N7O5 | Alkaloids | Pseudoalkaloids (transamidation) | Not annotated |
| 14680 | C22H35N5O5 | Alkaloids | Pseudoalkaloids (transamidation) | Not annotated |
| 15105 | C26H37NO6 | Terpenoids | Diterpenoids | Tigliane diterpenoids |
| 16345 | Not annotated | Not annotated | Not annotated | Not annotated |
| 22299 | C29H46O2 | Terpenoids | Meroterpenoids | Prenyl quinone meroterpenoids |
| **25552** | **C24H34O2** | **Terpenoids** | **Steroids** | **Cholane steroids** |
| 33591 | C19H30O2 | Terpenoids | Diterpenoids | Androstane steroids |
| 33598 | C9H8O2 | Shikimates and Phenylpropanoids | Phenolic acids (C6-C1) | Simple phenolic acids |
| 34551 | C10H17N3O2 | Alkaloids | Histidine alkaloids | Imidazole alkaloids |
| 46387 | C19H35N7O3S | Amino acids and Peptides | Not annotated | Not annotated |
| 58021 | C8H10N4O2 | Alkaloids | Pseudoalkaloids (transamidation) | Purine alkaloids |

**Table S5 |** Alpha-diversity correlations comparing microbially-related metabolite richness vs. either shotgun metagenomic microbial taxonomic richness, Faith's PD, or weighted Faith's PD. Values are Spearman correlation coefficients, and *p*-values were adjusted using the Benjamini-Hochberg correction.

| EMPO | Environment | *n* | shotgun_metric | *r* | lower 95% CI | upper 95% CI | *t* | *S* | *p*-value |
|---|---|---|---|---|---|---|---|---|---|
| EMPO 0 | All | 454 | richness | **0.2** | 0.11 | 0.28 | 4.23 | N/A | **0.001** |
| EMPO 1 | Host-associated | 271 | richness | **0.19** | 0.08 | 0.31 | 3.23 | N/A | **0.01** |
| EMPO 1 | Free-living | 183 | richness | **0.18** | 0.03 | 0.31 | 2.4 | N/A | **0.05** |
| EMPO 4 | Animal corpus (non-saline) | 3 | richness | N/A | N/A | N/A | N/A | N/A | N/A |
| EMPO 4 | Animal corpus (saline) | 44 | richness | **0.39** | 0.11 | 0.62 | 2.74 | N/A | **0.01** |
| EMPO 4 | Animal distal gut (non-saline) | 139 | richness | 3.43E-04 | -0.17 | 0.17 | 4.01E-03 | N/A | 1 |
| EMPO 4 | Animal distal gut (saline) | 11 | richness | -0.12 | -0.68 | 0.53 | N/A | 274.06 | 0.7 |
| EMPO 4 | Animal proximal gut (non-saline) | 5 | richness | -0.21 | -0.93 | 0.84 | N/A | 24.1 | 0.7 |
| EMPO 4 | Animal proximal gut (saline) | 12 | richness | **0.73** | 0.26 | 0.92 | N/A | 76 | **0.01** |
| EMPO 4 | Animal secretion (saline) | 12 | richness | 0.05 | -0.55 | 0.62 | N/A | 272 | 0.9 |
| EMPO 4 | Fungus corpus (non-saline) | 6 | richness | 0.43 | -0.61 | 0.93 | N/A | 20 | 0.4 |
| EMPO 4 | Plant detritus (non-saline) | 26 | richness | **0.74** | 0.49 | 0.88 | N/A | 750.63 | **0.001** |
| EMPO 4 | Plant surface (non-saline) | 13 | richness | -0.36 | -0.77 | 0.26 | N/A | 494 | 0.2 |
| EMPO 4 | Plant surface (saline) | 0 | richness | N/A | N/A | N/A | N/A | N/A | N/A |
| EMPO 4 | Sediment (non-saline) | 31 | richness | **0.42** | 0.07 | 0.67 | 2.47 | N/A | **0.05** |
| EMPO 4 | Sediment (saline) | 31 | richness | 0.27 | -0.09 | 0.57 | 1.52 | N/A | 0.1 |
| EMPO 4 | Soil (non-saline) | 78 | richness | 0.06 | -0.16 | 0.28 | 0.54 | N/A | 0.6 |
| EMPO 4 | Soil (saline) | 0 | richness | N/A | N/A | N/A | N/A | N/A | N/A |
| EMPO 4 | Subsurface (non-saline) | 3 | richness | N/A | N/A | N/A | N/A | N/A | N/A |
| EMPO 4 | Surface (saline) | 2 | richness | N/A | N/A | N/A | N/A | N/A | N/A |
| EMPO 4 | Water (non-saline) | 8 | richness | 0.24 | -0.56 | 0.81 | 0.6 | N/A | 0.6 |
| EMPO 4 | Water (saline) | 30 | richness | **0.57** | 0.26 | 0.77 | 3.65 | N/A | **0.01** |
| EMPO 0 | All | 454 | Faith's PD | **0.16** | 0.07 | 0.25 | 3.52 | N/A | **0.001** |
| EMPO 1 | Host-associated | 271 | Faith's PD | **0.17** | 0.05 | 0.28 | 2.84 | N/A | **0.01** |
| EMPO 1 | Free-living | 183 | Faith's PD | 0.14 | 0 | 0.28 | 1.93 | N/A | 0.06 |
| EMPO 4 | Animal corpus (non-saline) | 3 | Faith's PD | N/A | N/A | N/A | N/A | N/A | N/A |
| EMPO 4 | Animal corpus (saline) | 44 | Faith's PD | **0.29** | -0.01 | 0.54 | 1.98 | N/A | **0.05** |
| EMPO 4 | Animal distal gut (non-saline) | 139 | Faith's PD | 0.02 | -0.15 | 0.18 | 0.19 | N/A | 0.8 |
| EMPO 4 | Animal distal gut (saline) | 11 | Faith's PD | -0.05 | -0.64 | 0.58 | N/A | 230 | 0.9 |
| EMPO 4 | Animal proximal gut (non-saline) | 5 | Faith's PD | 0.1 | -0.87 | 0.91 | N/A | 18 | 0.9 |
| EMPO 4 | Animal proximal gut (saline) | 12 | Faith's PD | **0.72** | 0.23 | 0.92 | N/A | 80 | **0.01** |
| EMPO 4 | Animal secretion (saline) | 12 | Faith's PD | 0.17 | -0.46 | 0.69 | N/A | 238 | 0.6 |
| EMPO 4 | Fungus corpus (non-saline) | 6 | Faith's PD | 0.43 | -0.61 | 0.93 | N/A | 20 | 0.4 |
| EMPO 4 | Plant detritus (non-saline) | 26 | Faith's PD | **0.74** | 0.49 | 0.88 | N/A | 760 | **0.001** |
| EMPO 4 | Plant surface (non-saline) | 13 | Faith's PD | -0.41 | -0.79 | 0.2 | N/A | 512 | 0.2 |
| EMPO 4 | Plant surface (saline) | 0 | Faith's PD | N/A | N/A | N/A | N/A | N/A | N/A |
| EMPO 4 | Sediment (non-saline) | 31 | Faith's PD | **0.44** | 0.1 | 0.69 | 2.65 | N/A | **0.05** |
| EMPO 4 | Sediment (saline) | 31 | Faith's PD | 0.09 | -0.28 | 0.43 | 0.47 | N/A | 0.6 |
| EMPO 4 | Soil (non-saline) | 78 | Faith's PD | 8.73E-04 | -0.22 | 0.22 | 7.61E-03 | N/A | 1 |
| EMPO 4 | Soil (saline) | 0 | Faith's PD | N/A | N/A | N/A | N/A | N/A | N/A |
| EMPO 4 | Subsurface (non-saline) | 3 | Faith's PD | N/A | N/A | N/A | N/A | N/A | N/A |
| EMPO 4 | Surface (saline) | 2 | Faith's PD | N/A | N/A | N/A | N/A | N/A | N/A |
| EMPO 4 | Water (non-saline) | 8 | Faith's PD | -0.24 | -0.81 | 0.56 | -0.6 | N/A | 0.6 |
| EMPO 4 | Water (saline) | 30 | Faith's PD | **0.6** | 0.31 | 0.79 | 4.01 | N/A | **0.001** |
| EMPO 0 | All | 454 | weighted Faith's PD | 0.08 | -0.01 | 0.17 | 1.77 | N/A | 0.08 |
| EMPO 1 | Host-associated | 271 | weighted Faith's PD | 0.08 | -0.04 | 0.19 | 1.2 | N/A | 0.2 |
| EMPO 1 | Free-living | 183 | weighted Faith's PD | 0.09 | -0.05 | 0.23 | 1.2 | N/A | 0.2 |
| EMPO 4 | Animal corpus (non-saline) | 3 | weighted Faith's PD | N/A | N/A | N/A | N/A | N/A | N/A |
| EMPO 4 | Animal corpus (saline) | 44 | weighted Faith's PD | -0.13 | -0.41 | 0.17 | -0.87 | N/A | 0.4 |
| EMPO 4 | Animal distal gut (non-saline) | 139 | weighted Faith's PD | 0.05 | -0.11 | 0.22 | 0.64 | N/A | 0.5 |
| EMPO 4 | Animal distal gut (saline) | 11 | weighted Faith's PD | -0.39 | -0.81 | 0.29 | N/A | 306 | 0.2 |
| EMPO 4 | Animal proximal gut (non-saline) | 5 | weighted Faith's PD | 0.1 | -0.87 | 0.91 | N/A | 18 | 0.9 |
| EMPO 4 | Animal proximal gut (saline) | 12 | weighted Faith's PD | 0.53 | -0.08 | 0.85 | N/A | 134 | 0.08 |
| EMPO 4 | Animal secretion (saline) | 12 | weighted Faith's PD | 0.31 | -0.34 | 0.76 | N/A | 198 | 0.3 |
| EMPO 4 | Fungus corpus (non-saline) | 6 | weighted Faith's PD | 0.6 | -0.44 | 0.95 | N/A | 14 | 0.2 |
| EMPO 4 | Plant detritus (non-saline) | 26 | weighted Faith's PD | **0.46** | 0.07 | 0.72 | N/A | 1585 | **0.05** |
| EMPO 4 | Plant surface (non-saline) | 13 | weighted Faith's PD | -0.45 | -0.81 | 0.15 | N/A | 528 | 0.12 |
| EMPO 4 | Plant surface (saline) | 0 | weighted Faith's PD | N/A | N/A | N/A | N/A | N/A | N/A |
| EMPO 4 | Sediment (non-saline) | 31 | weighted Faith's PD | **0.35** | 0 | 0.63 | 2.02 | N/A | **0.05** |
| EMPO 4 | Sediment (saline) | 31 | weighted Faith's PD | -0.19 | -0.51 | 0.17 | -0.105 | N/A | 0.3 |
| EMPO 4 | Soil (non-saline) | 78 | weighted Faith's PD | -6.16E+03 | -0.23 | 0.22 | -0.05 | N/A | 1 |
| EMPO 4 | Soil (saline) | 0 | weighted Faith's PD | N/A | N/A | N/A | N/A | N/A | N/A |
| EMPO 4 | Subsurface (non-saline) | 3 | weighted Faith's PD | N/A | N/A | N/A | N/A | N/A | N/A |
| EMPO 4 | Surface (saline) | 2 | weighted Faith's PD | N/A | N/A | N/A | N/A | N/A | N/A |
| EMPO 4 | Water (non-saline) | 8 | weighted Faith's PD | 0.12 | -0.64 | 0.76 | 0.29 | N/A | 0.8 |
| EMPO 4 | Water (saline) | 30 | weighted Faith's PD | **0.42** | 0.08 | 0.68 | 2.48 | N/A | **0.05** |

**Table S6 |** The twenty most highly ranked molecules among all microbially-related metabolites (*n* = 6,588) contributing to separation of environments, identified by machine-learning analysis. Mean ranks are based on impacts derived from SHAP values. Associations with environments are indicated, where + indicates a positive association and – indicates a negative association based on feature abundances. The number of iterations (*n* = 20) for which the association was observed is shown. Metabolites in bold font are those also identified to be strongly associated with particular environments in analyses of differential abundance (Table S3).

| Feature ID | Mean rank | Direction | Associated environment | Iterations | Molecular formula | Pathway | Superclass | Class |
|---|---|---|---|---|---|---|---|---|
| 4492 | 6.7 | – | Soil (non-saline) | 20 | C20H32 | Terpenoids | Diterpenoids | Not annotated |
| **42202** | **7.85** | **+** | **Sediment (saline)** | **20** | **Not annotated** | **Not annotated** | **Not annotated** | **Not annotated** |
| 7899 | 10.65 | – | Sediment (non-saline) | 20 | C20H20O5 | Shikimates and Phenylpropanoids | Lignans | Dibenzylbutyrolactone lignans |
| 7719 | 13.7 | – | Plant surface (non-saline) | 20 | C20H28O3 | Terpenoids | Diterpenoids | Pimarane and Isopimarane diterpenoids |
| 14598 | 23.55 | + | Subsurface (non-saline) | 20 | Not annotated | Not annotated | Not annotated | Not annotated |
| 29044 | 166.85 | + | Plant detritus (non-saline) | 19 | C6H8O4 | Carbohydrates | Saccharides | Monosaccharides |
| 12117 | 181.95 | + | Animal distal gut (saline) | 19 | C24H39NO3 | Terpenoids | Steroids | Cholane steroids |
| **24213** | **192.5** | **+** | **Animal secretion (saline)** | **19** | **C25H40O3** | **Fatty acids** | **Glycerolipids** | **Monoacylglycerols** |
| 150 | 246.85 | – | Soil (non-saline) | 18 | C8H9N | Alkaloids | Tyrosine alkaloids | Phenylethylamines |
| 15657 | 377.75 | + | Water (non-saline) | 18 | C15H29N5O5 | Carbohydrates | Not annotated | Not annotated |
| 187 | 428 | – | Soil (non-saline) | 18 | C16H26 | Fatty acids | Fatty Acids and Conjugates | Not annotated |
| 6196 | 520.6 | – | Sediment (non-saline) | 17 | C12H18 | Fatty acids | Fatty esters | Not annotated |
| 4542 | 663.5 | + | Water (non-saline) | 17 | C21H37NO11 | Terpenoids | Not annotated | Not annotated |
| 15687 | 696.4 | + | Water (non-saline) | 15 | C15H29N5O5 | Alkaloids | Not annotated | Not annotated |
| 50331 | 696.5 | + | Water (non-saline) | 16 | Not annotated | Not annotated | Not annotated | Not annotated |
| 15245 | 761.95 | + | Water (non-saline) | 15 | C12H20O7 | Fatty acids | Not annotated | Not annotated |
| 15681 | 876.9 | + | Water (non-saline) | 15 | C23H43N5O8 | Alkaloids | Not annotated | Not annotated |
| 1947 | 922.55 | + | Sediment (saline) | 11 | C30H52O8 | Polyketides | Linear polyketides | Open-chain polyketides |
| 15250 | 978.5 | + | Water (non-saline) | 13 | C8H12O4 | Carbohydrates | Saccharides | Not annotated |
| 48746 | 982.15 | + | Animal distal gut (non-saline) | 14 | C17H24N2O4 | Alkaloids | Ornithine alkaloids | Not annotated |

**Table S7 | Comparison of differentials from *songbird* runs using different references.** For each environment, we ran Spearman correlations between our original run using Animal distal gut (non-saline) as the reference, with one run using the same reference, and with one run using a different reference (Water [saline]).

| environment | *r* (same reference) | *r* (different reference) |
|---|---|---|
| Animal corpus (saline) | 1.00 | 0.89 |
| Animal distal gut (non-saline) | 1.00 | NA |
| Animal distal gut (saline) | 0.99 | 0.90 |
| Animal proximal gut (saline) | 1.00 | 0.91 |
| Animal secretion (saline) | 0.99 | 0.92 |
| Fungus corpus (non-saline) | 0.98 | 0.92 |
| Plant corpus (non-saline) | 1.00 | 0.88 |
| Plant surface (non-saline) | 0.87 | 0.83 |
| Plant surface (saline) | 0.99 | 0.91 |
| Sediment (non-saline) | 1.00 | 0.87 |
| Sediment (saline) | 1.00 | 0.87 |
| Soil (non-saline) | 1.00 | 0.86 |
| Water (non-saline) | 0.91 | 0.87 |
| Water (saline) | 0.99 | NA |

### SUPPLEMENTAL FIGURES

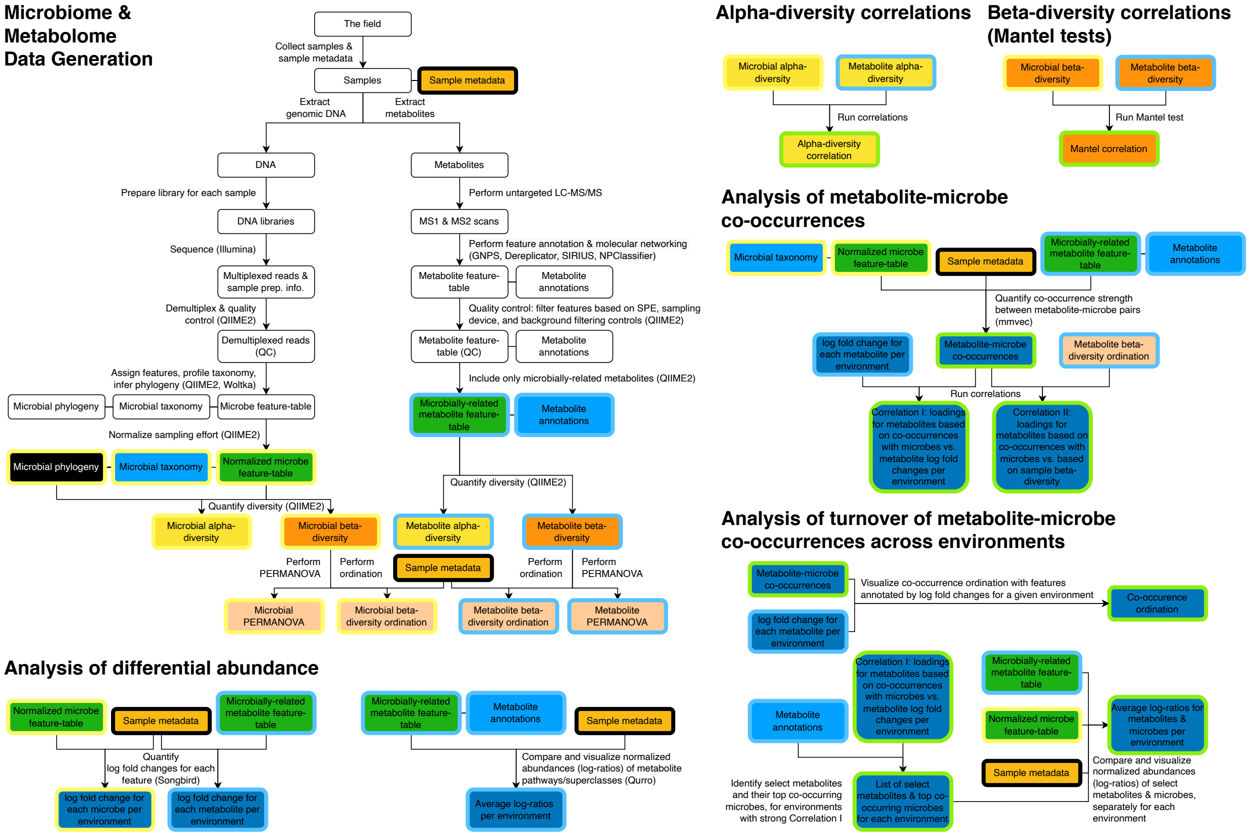


**Figure S1 | Diagrammatic overview of multi-omics analyses performed using the EMP500 dataset.** The process begins with data generation for both the microbiome and metabolome, which is then followed by analysis of differential abundance of both microbial taxa and microbially-related metabolites across environments. To begin multi-omics integration, correlations between alpha- and beta-diversity are explored, followed by explicit co-occurrence analysis of metabolite-microbe pairs. The results from analysis of co-occurrence are then combined with those from analysis of differential abundance, to reveal strong patterns of metabolite-microbe turnover across environments. Throughout the diagram, artifacts derived from microbial data are outlined in yellow, those derived from metabolite data are outlined in blue, and those derived from co-occurrence analysis are outlined in green.

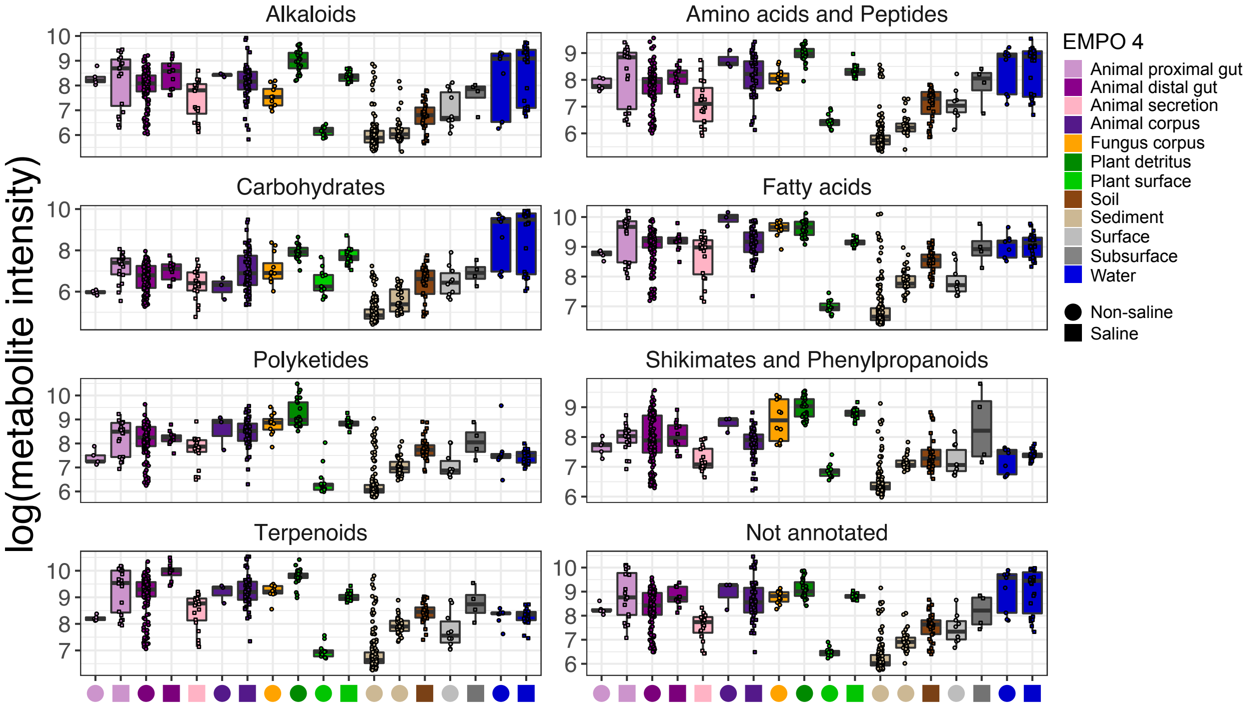


**Figure S2 | Relative abundance of microbially-related metabolite pathways, highlighting among-sample variation for each environment.** These data are shown as a complement to those in Fig. 2b of the main text. We note that as abundance data were not normalized (e.g., by using log-ratios as in Fig. 3a), caution should be used in interpreting differences among environments. Boxplots are in the style of Tukey, where the center line indicates the median, lower and upper hinges the first- and third quartiles, respectively, and each whisker 1.5 x the interquartile range (IQR) from its respective hinge.

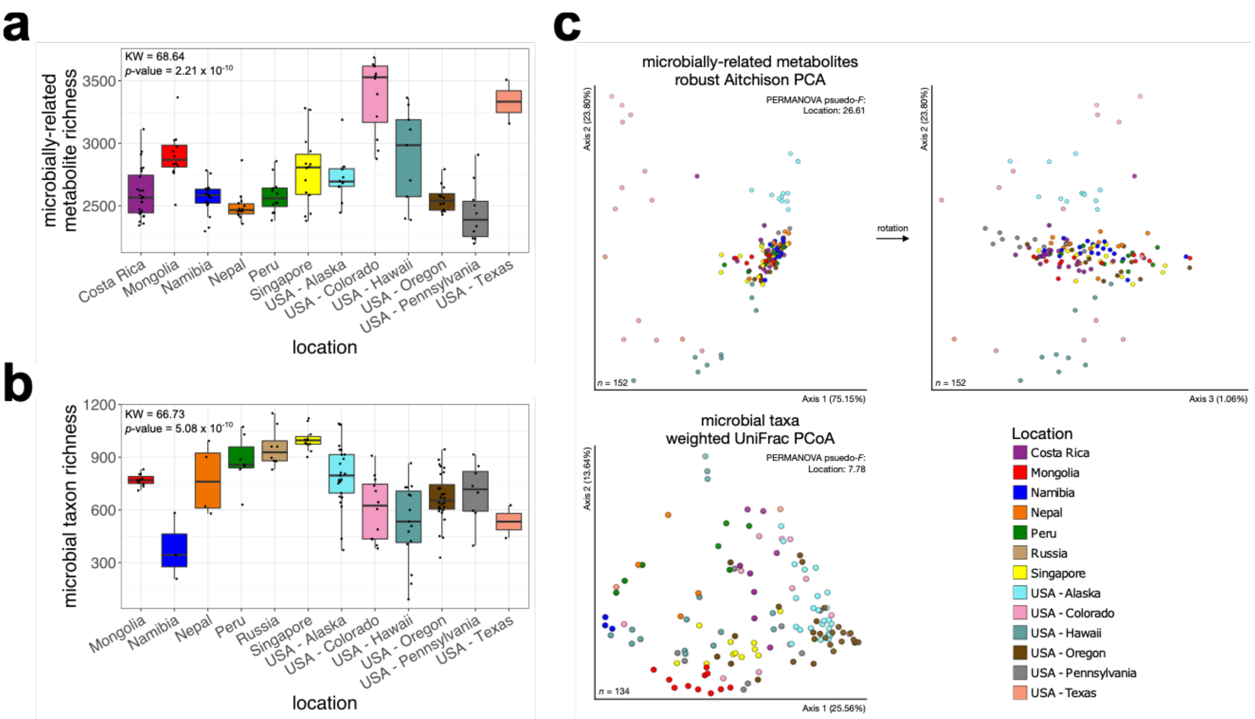


**Figure S3 | Microbially-related metabolite and microbial taxon composition among geographic locations for all non-saline soil samples. a,** Metabolite richness. **b,** Microbe richness. For **a** and **b**, the chi-squared statistic from a Kruskal-Wallis rank sum test for differences in richness across environments is shown (i.e., each test had *p*-value < $2.2 \times 10^{-16}$). **c,** Beta-diversity based on metabolites (upper panel) and microbes (lower panel). Results from PERMANOVA tests (*n* = 999 permutations) for variance explained by salinity as well as each level of EMPO are shown; *p*-value = 0.001 for all tests.

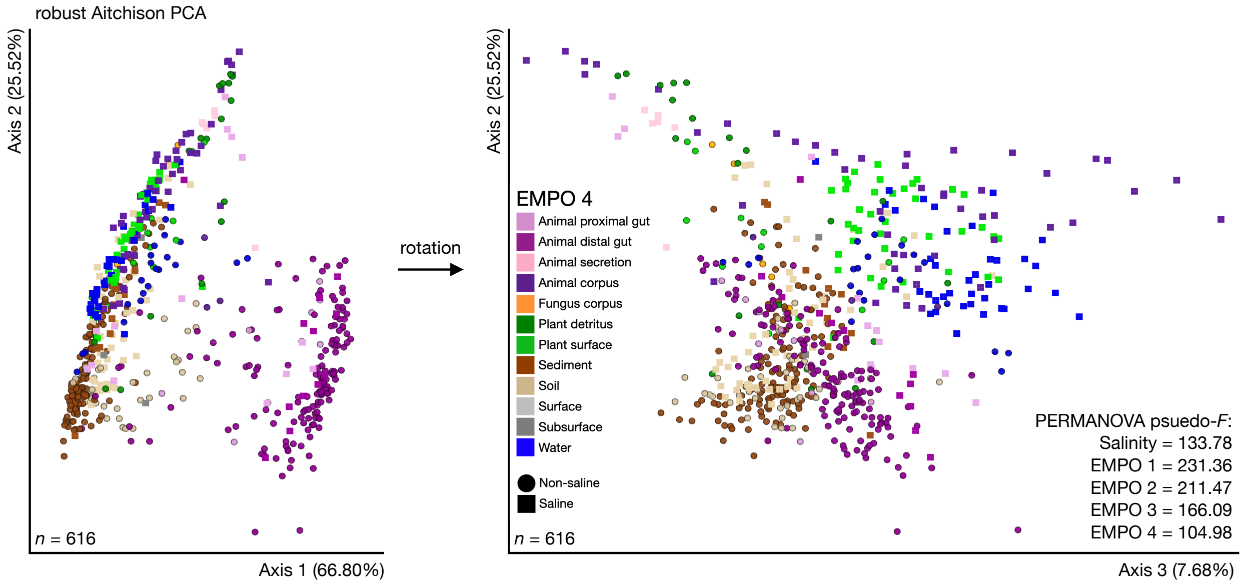


**Figure S4 | Clustering of samples by environments highlighting beta-diversity based on shotgun metagenomics data for microbial functions.** Robust Aitchison PCA with samples colored by EMPO 4 and shaped by salinity. Features are KEGG ECs (i.e., enzymes). Results from PERMANOVA tests ($n$ = 999 permutations) for variance explained by salinity as well as each level of EMPO are shown; $p$-value = 0.001 for all tests.

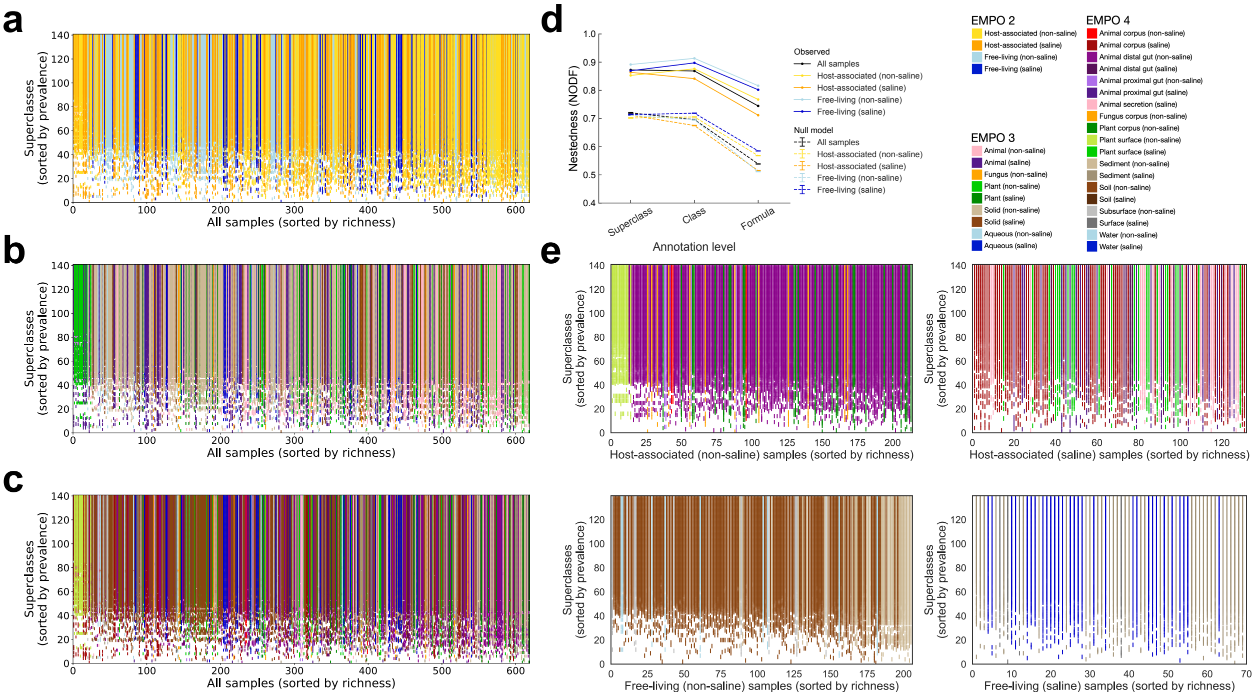


**Figure S5 | Nestedness of community composition based on microbially-related metabolites.** **a,** Presence-absence of superclasses across samples, with superclasses (rows) sorted by prevalence and samples (columns, $n$ = 618) sorted by richness. With increasing sample richness, superclasses tended to be gained but not lost (SES = 108.61, $p$-value < 0.0001 vs. a null model; nestedness measure based on overlap and decreasing fills [NODF] statistic = 0.87). Samples are colored by EMPO 2. **b,** As in **a** but with samples colored by EMPO 3. **c,** As in **a** but with samples colored by EMPO 4. **d,** Nestedness as a function of annotation level, from superclass to molecular formula, across all samples and within environments based on EMPO 2. Also shown are median null model NODF scores (± s.d.) for all samples, as well as samples at each level of EMPO 2. NODF measures the average fraction of metabolites from less diverse communities that occur in more diverse communities. All environments at all annotation levels examined were more nested than expected randomly, with nestedness higher at higher annotation levels ($p$-value < 0.0001 for all comparisons). **e,** As in **c** but with each environment at EMPO 2 shown separately, with samples colored by EMPO 4.

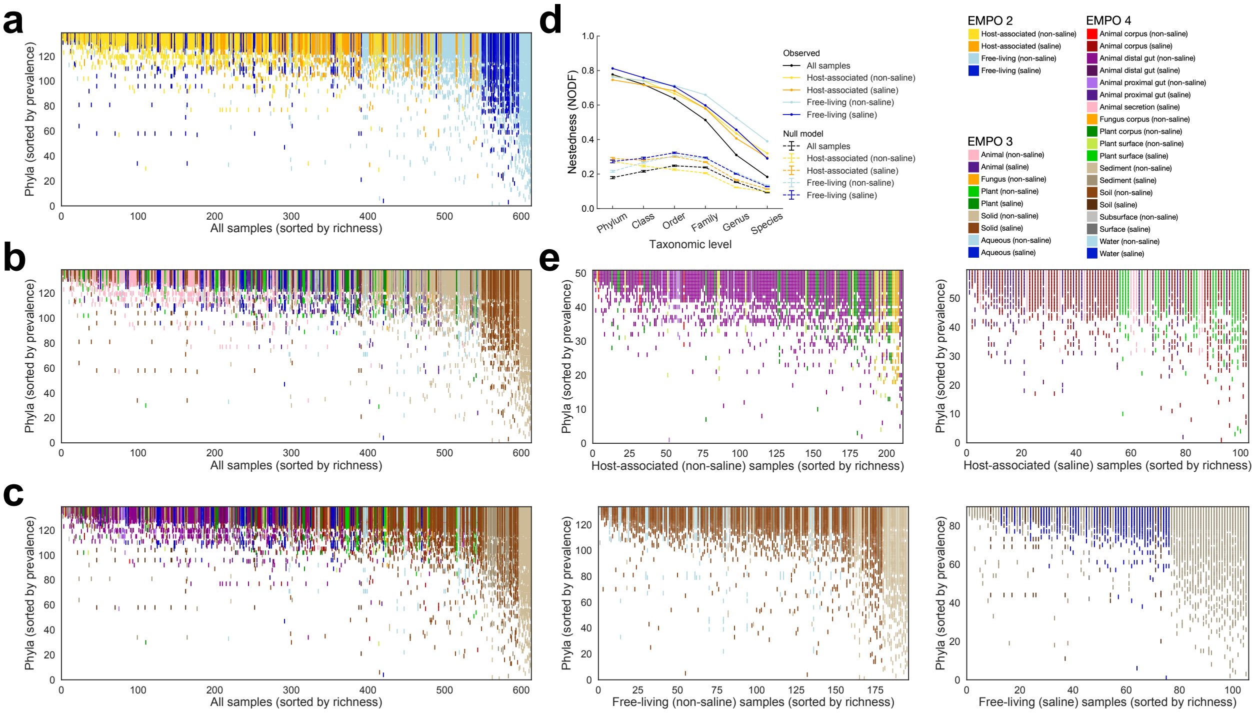


**Figure S6 | Nestedness of community composition based on microbial taxa. a,** Presence-absence of phyla across samples, with phyla (rows) sorted by prevalence and samples (columns, $n$ = 612) sorted by richness. With increasing sample richness, phyla tended to be gained but not lost (SES = 91.86, $p$-value < 0.0001 vs. a null model; nestedness measure based on overlap and decreasing fills [NODF] statistic = 0.78). Samples are colored by EMPO 2. **b,** As in **a** but with samples colored by EMPO 3. **c,** As in **a** but with samples colored by EMPO 4. **d,** Nestedness as a function of taxonomic level, from phylum to species, across all samples and within environments based on EMPO 2. Also shown are median null model NODF scores (± s.d.) for all samples, as well as samples at each level of EMPO 2. NODF measures the average fraction of taxa from less diverse communities that occur in more diverse communities. All environments at all taxonomic levels examined were more nested than expected randomly, with nestedness higher at higher taxonomic levels ($p$-value < 0.0001 for all comparisons). **e,** As in **c** but with each environment at EMPO 2 shown separately, with samples colored by EMPO 4.

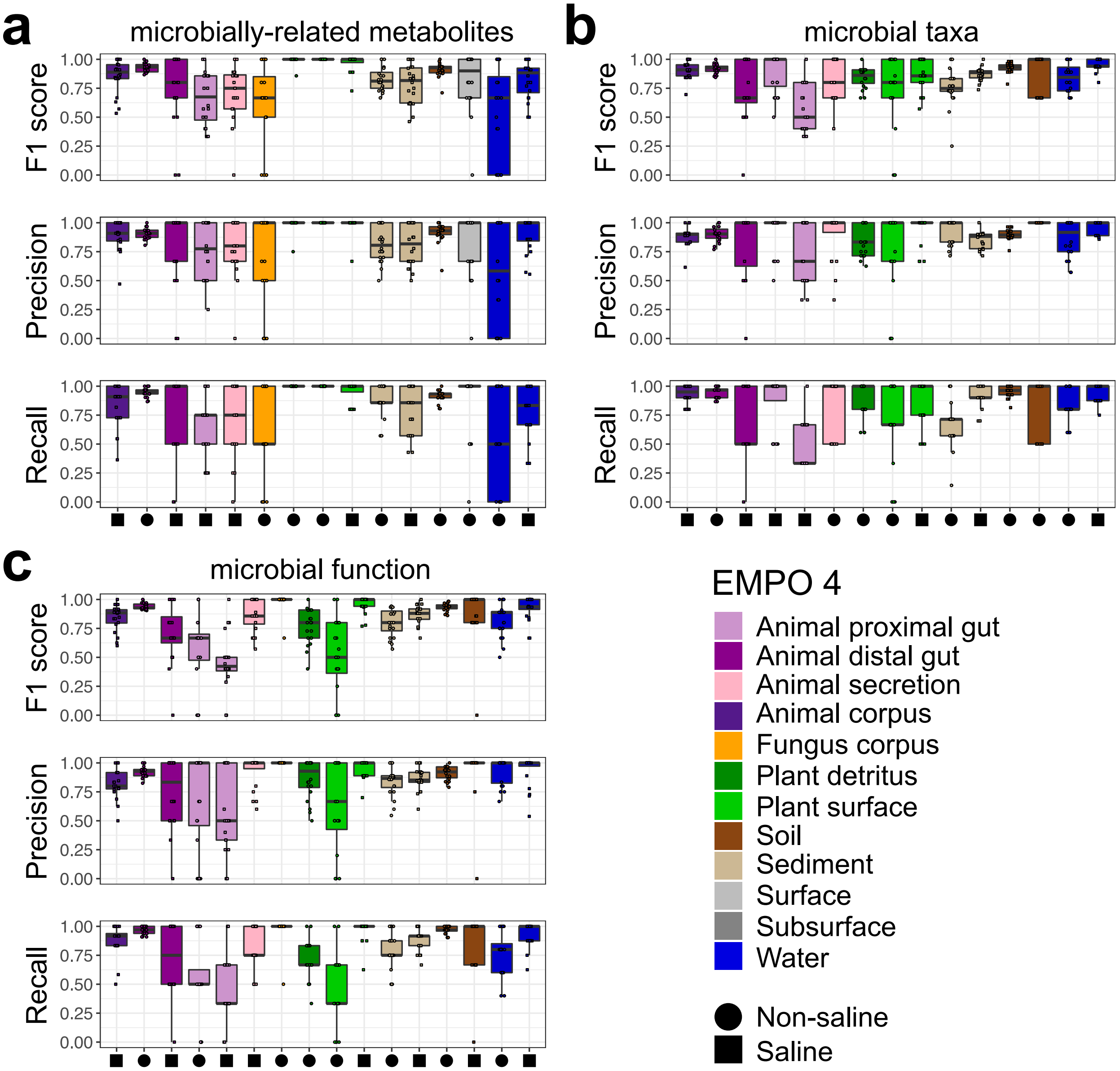


**Figure S7 | Machine-learning performance, highlighting F1 score, precision and recall across environments. a,** Microbially-related metabolites. **b,** Microbial taxa (i.e., OGUs). **c,** Microbial functions (i.e., KEGG ECs). Boxplots are in the style of Tukey, where the center line indicates the median, lower and upper hinges the first- and third quartiles, respectively, and each whisker 1.5 x the interquartile range (IQR) from its respective hinge.

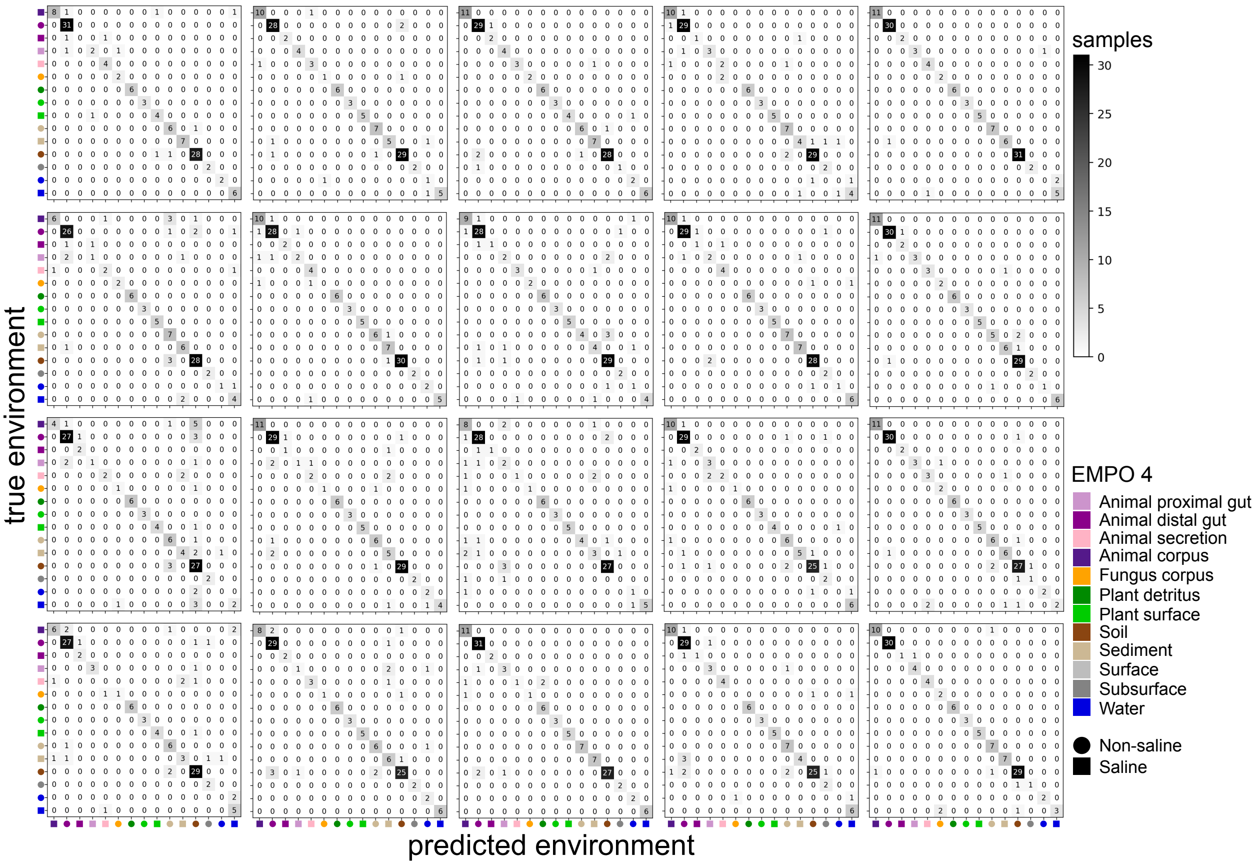


**Figure S8 | Machine learning performance for microbially-related metabolites, highlighting which environments are most often confused.** Data are from 20 iterations. The candidate confusion matrix shown in Fig. 5b of the main text is that in the first row, fifth column.

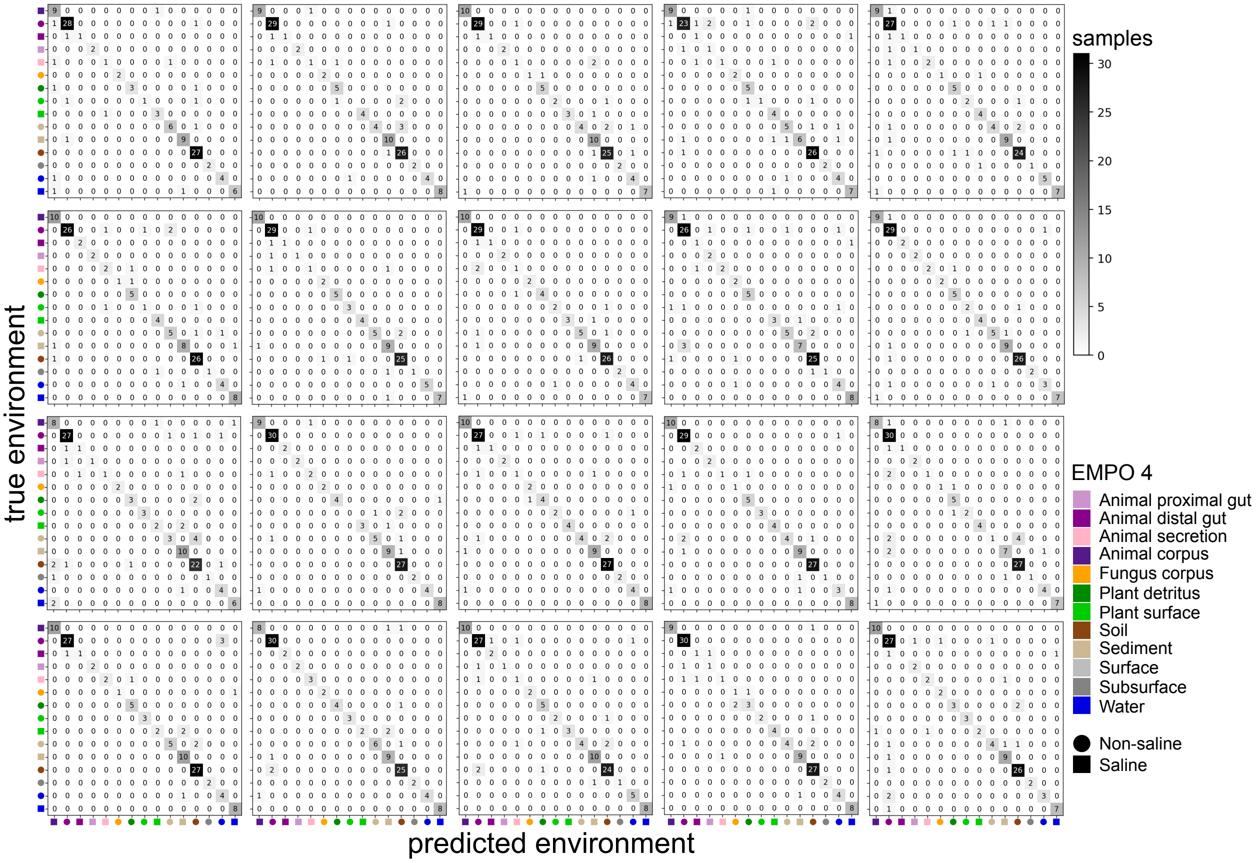


**Figure S9 | Machine learning performance for microbial taxa, highlighting which environments are most often confused.** Data are from 20 iterations. The candidate confusion matrix shown in Fig. 5b of the main text is that in the first row, third column.

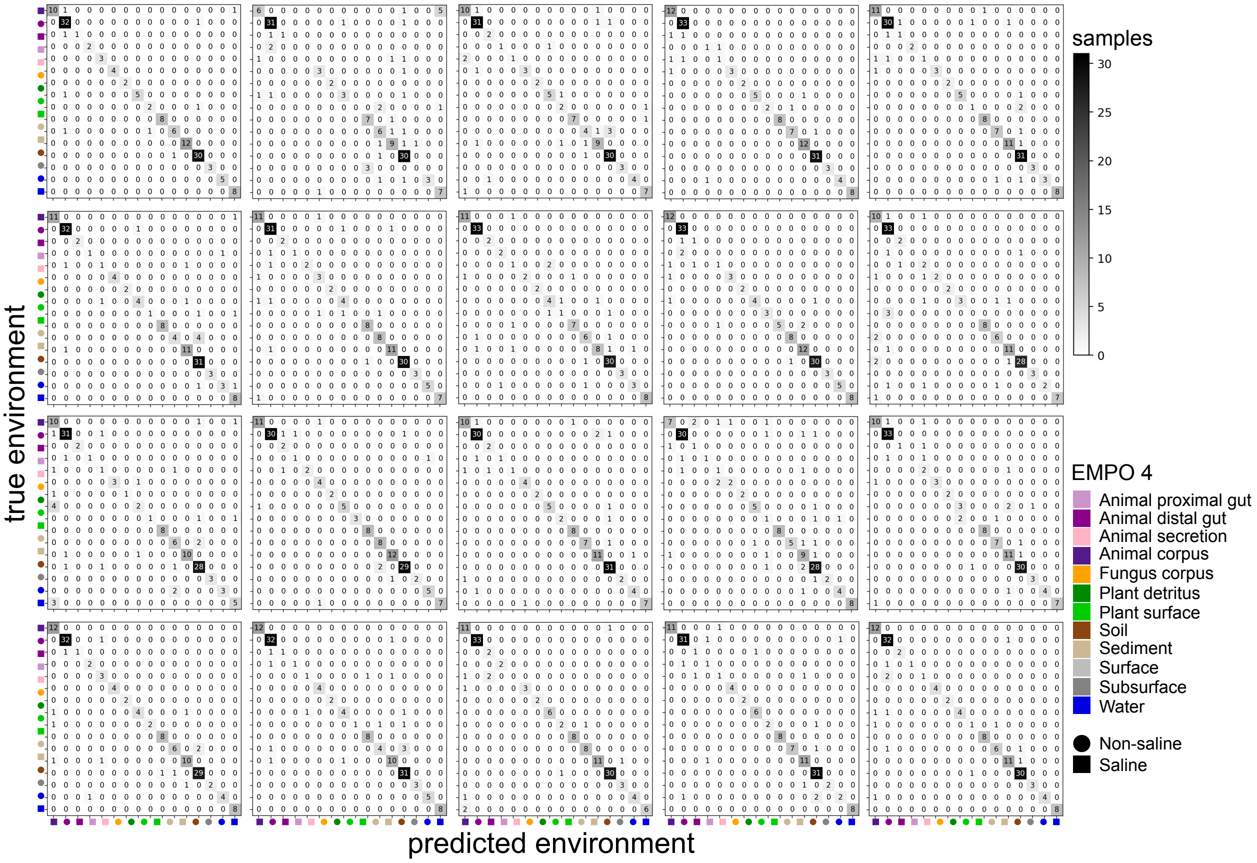


**Figure S10 | Machine learning performance for microbial functions, highlighting which environments are most often confused.** Data are from 20 iterations. The candidate confusion matrix shown in Fig. 5b of the main text is that in the first row, fourth column.

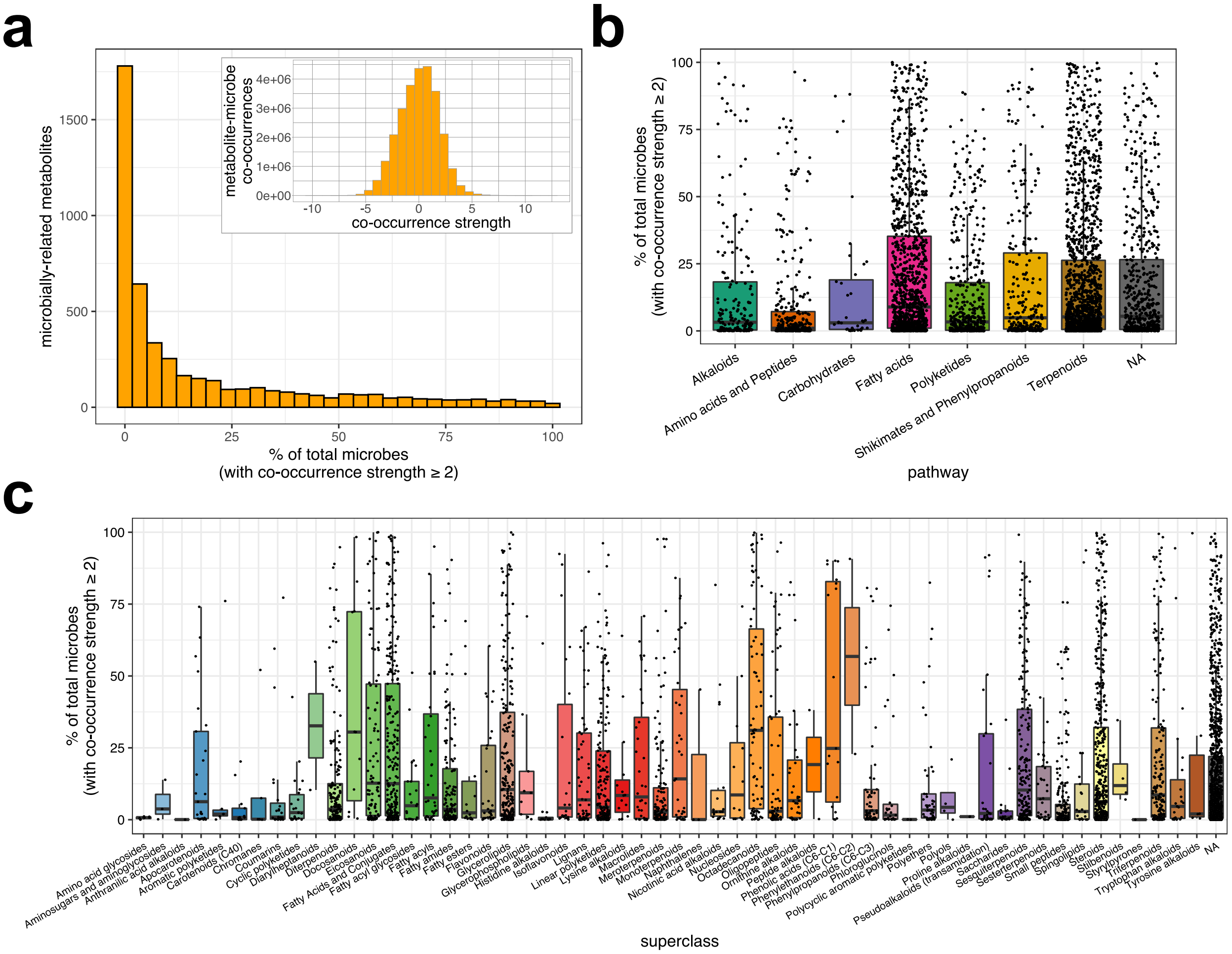


**Figure S11 | Summary of co-occurrence ranks for microbially-related metabolites. a,** Distribution of the percentage of microbial taxa for which co-occurrences were strong. Strong co-occurrence was defined as having a co-occurrence strength (i.e., rank, or log conditional probability) $\geq 2$. The overall distribution of co-occurrence strengths is shown in the inset ($n$ = 26,784,120). For values > 0 ($n$ = 13,851,755), the minimum = –10.17, maximum = 12.69, mean = 2.40 x $10^{-18}$, median = 0.08, and mode = 1.22. For values $\geq 2$ ($n$ = 3,496,639), the minimum = 2.00, maximum = 12.69, mean = 2.87, median = 2.63, and mode = 4.26. **b,** The percentage of microbial taxa for which co-occurrences were strong (i.e., $\geq 2$), across metabolite pathways. **c,** The percentage of microbial taxa for which co-occurrences were strong (i.e., $\geq 2$), across metabolite superclasses. For panels **b** and **c**, points were jittered horizontally for clarity. Boxplots are in the style of Tukey, where the center line indicates the median, lower and upper hinges the first- and third quartiles, respectively, and each whisker 1.5 x the interquartile range (IQR) from its respective hinge.

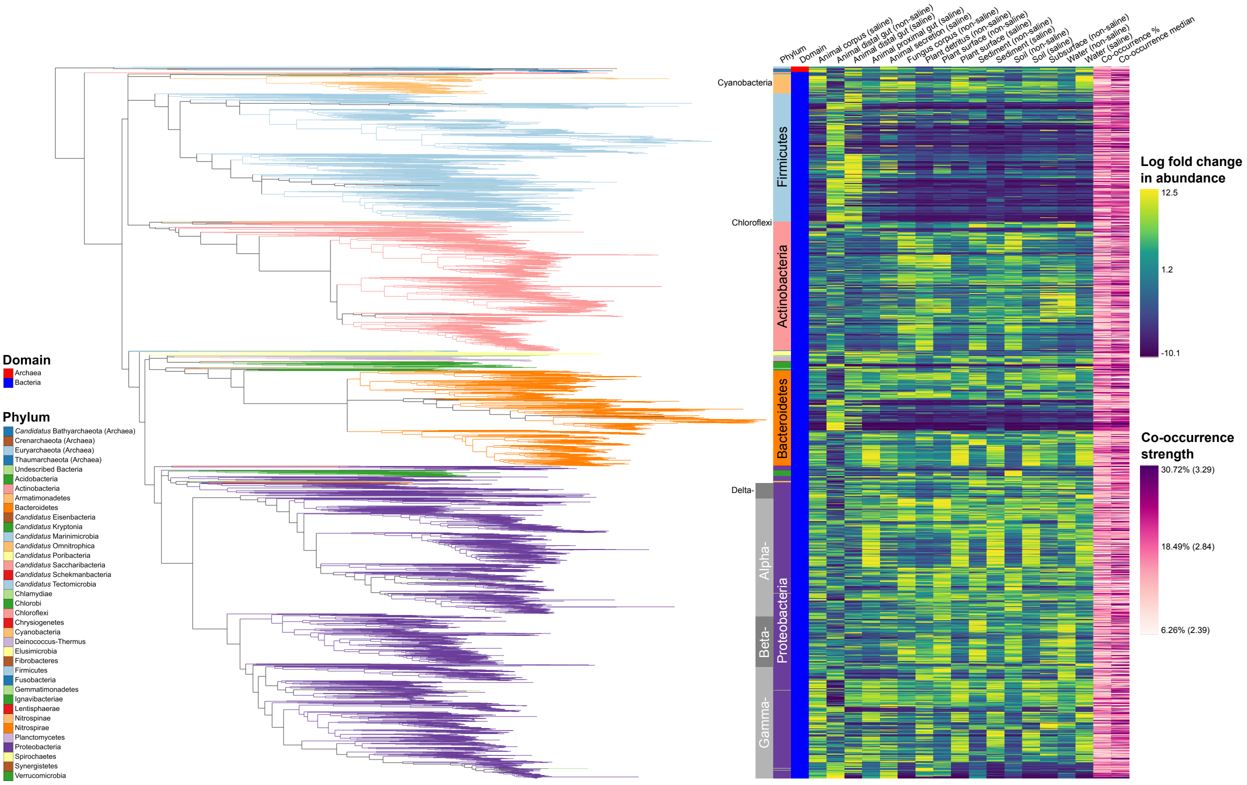


**Figure S12 | Phylogenetic relationships among microbial taxa highlighting log fold changes in abundance relative to environment, and overall co-occurrences with microbially-related metabolites.** Branches are colored by microbial phylum. Annotations include Domain and Phylum level associations (and Class for *Proteobacteria*), heat maps representing log fold changes in relative abundance for each environment (from *songbird*), and heat maps summarizing co-occurrences with microbially-related metabolites (from *mmvec*). Co-occurrence strength indicates (1) the percentage of all microbially-related metabolites for which the co-occurrence rank (i.e., log conditional probability) was ≥ 2 (i.e., strong), and (2) the median co-occurrence rank value, considering only strong values (in parentheses in the legend).

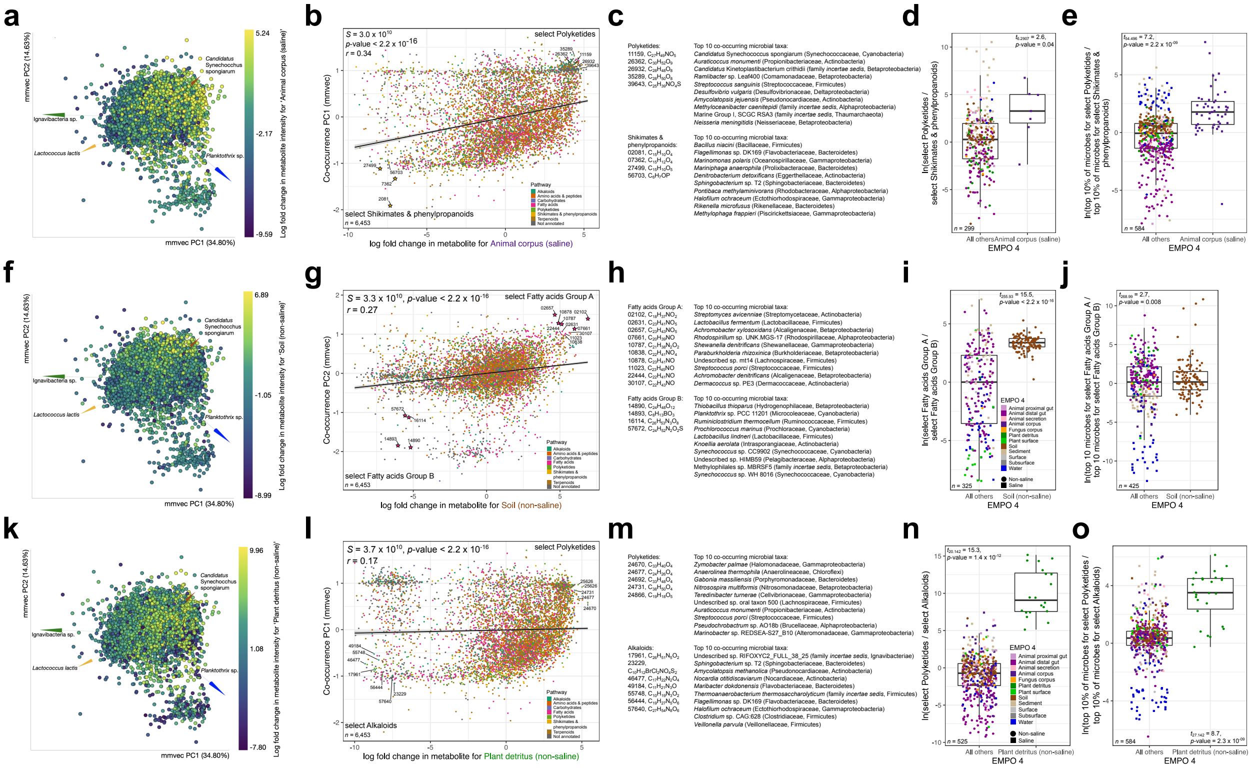

**Figure S13 | Metabolite-microbe co-occurrences reveal exhibit strong turnover across environments.** Further elaboration on Fig. 4 of the main text, including results from three environments in addition to ‘Water (saline)’, and to highlight differences driven by salinity and host-association: ‘Animal corpus (saline)’, ‘Soil (non-saline)’, and ‘Plant detritus (non-saline’). **a, f, k,** The relationship between log fold changes in abundance for metabolites with respect to the focal environment, and the first three *mmvec* PCs shown as a multi-omics biplot of metabolite–microbe co-occurrences learned from microbial profiles. Points represent metabolites, and the distance between metabolites indicates similarity in their co-occurrences with microbial taxa (i.e., two metabolites that are close together have similar co-occurrence probabilities to the same microbes). Metabolites are colored based on their relative log-fold changes with respect to the focal environment. Vectors represent specific microbial taxa, where distances between arrow tips indicate similarity in their co-occurrence with specific metabolites (i.e., two microbes that are close together have similar co-occurrence probabilities to the same metabolites), and the direction of each arrow indicates which metabolites each microbe co-occurs most strongly with. The model predicting metabolite–microbe co-occurrences was more accurate than one representing a random baseline, with a pseudo-$Q^2$ value of 0.18, indicating much reduced error during cross-validation. **b, g, l,** The relationship between log fold changes in metabolite abundances with respect to the focal environment and loadings for metabolites on PC1 of the co-occurrence ordination. The correlation is one example of those summarized in Fig. 6a. Metabolites are colored by pathway. Select features representing the focal group and reference group are highlighted. **c, h, m,** The top 10 co-occurring microbial taxa for all select focal group features and all select reference group features with respect to the focal environment, with a heatmap showing the co-occurrence strength between each metabolite-microbe pair. **d, i, n,** Log-ratio of metabolite intensities for select focal group features and select reference group features with respect to the focal environment. **e, j, o,** Log-ratio of abundances of the top ten microbial taxa associated with focal group features and the top ten microbial taxa associated with reference group features with respect to the focal environment. For panels **d, e, i, j, n, o** samples are colored by environment (based on EMPO level 4), and results from a *t*-test comparing the focal vs. all other environments are shown. Boxplots are in the style of Tukey, where the center line indicates the median, lower and upper hinges the first- and third quartiles, respectively, and each whisker 1.5 x the interquartile range (IQR) from its respective hinge.

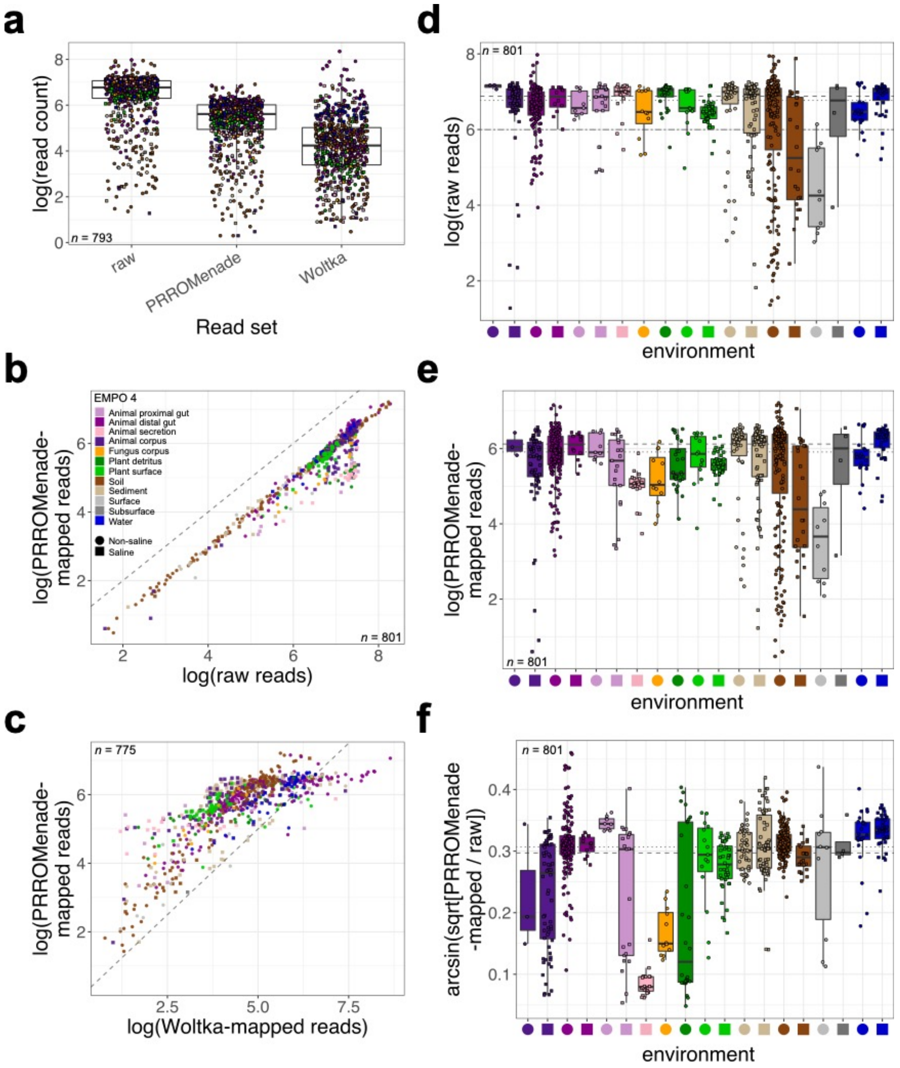


**Figure S14 | Summary of shotgun metagenomics read mapping for microbial functional profiling. a,** Comparison of read counts among raw reads, reads mapped using PRROMenade, and reads mapped using Woltka. **b,** Relationship between counts of raw reads and those mapped to using PRROMenade. The gray, dashed line indicates $y = x$. **c,** Relationship between counts of reads mapped using Woltka and those mapped using PRROMenade. The dashed line indicates $y = x$. **d**, Comparison of counts of raw reads among environments (based on EMPO 4). The two-dashed line indicates our expectation of 1 million reads from gut samples. **e,** Comparison of counts of reads mapped to using PRROMenade among environments. **f**, Comparison of the proportion of raw reads that were mapped using PRROMenade among environments. For panels **d-e**, the dashed line indicates the global mean and the dotted line indicates the global median. For all panels, note the respective *y*-axis transformation used. Colors and shapes are described in the legend in panel **b**. Boxplots are in the style of Tukey, where the center line indicates the median, lower and upper hinges the first- and third quartiles, respectively, and each whisker 1.5 x the interquartile range (IQR) from its respective hinge.

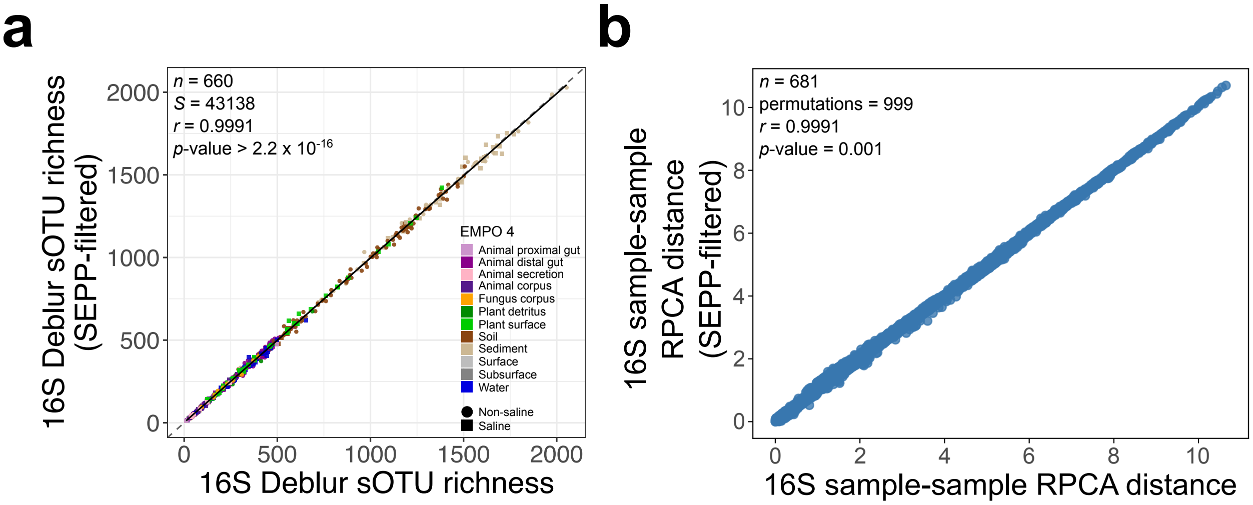


**Figure S15 | Comparison of alpha- and beta-diversity with and without removal of features that were not placed during fragment insertion (SEPP) for 16S data. a,** Spearman correlation in alpha-diversity (sOTU richness) between datasets, showing a high correlation across all environments. **b,** Mantel spearman correlation (999 permutations) in beta-diversity (sample-sample robust Aitchison distances) between datasets, showing a high correlation across all environments.

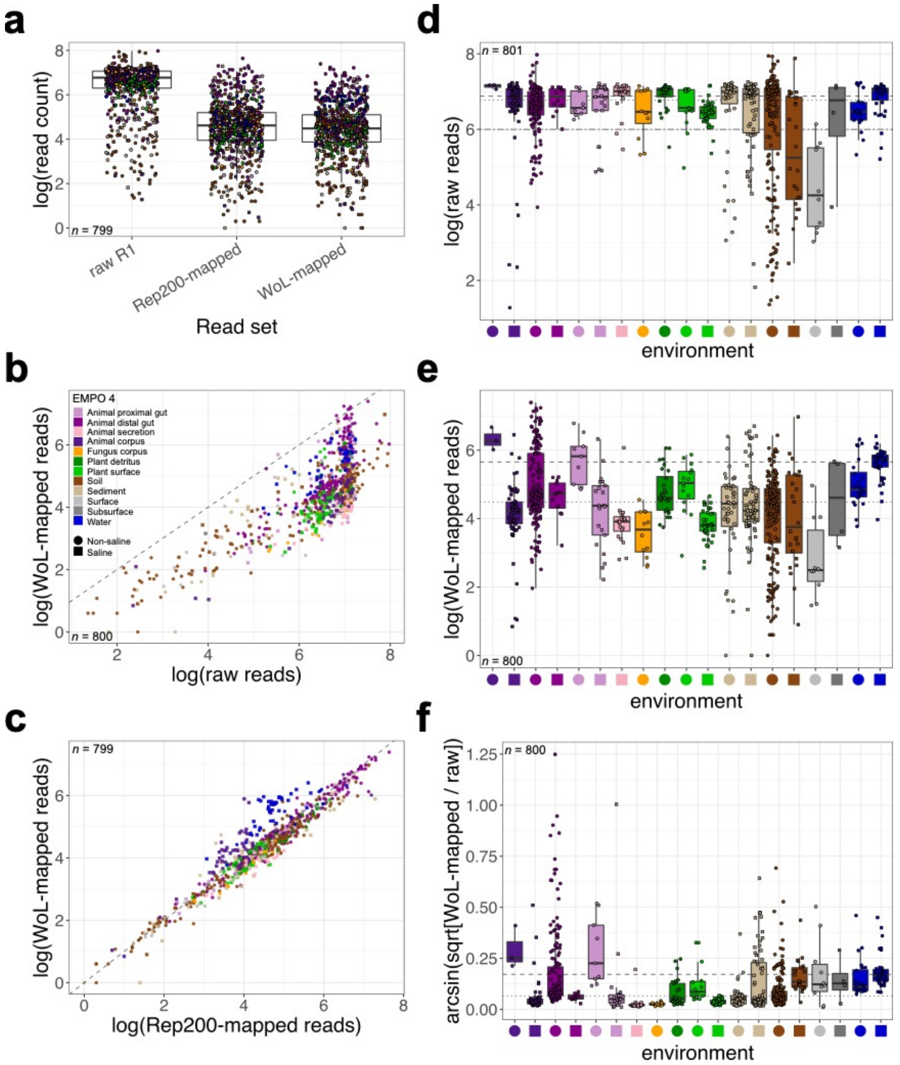


**Figure S16 | Summary of shotgun metagenomics read mapping for microbial taxonomic profiling. a,** Comparison of read counts among raw reads, reads mapped to NCBI's Rep200, and reads mapped to the Web of Life (WoL). **b,** Relationship between counts of raw reads and those mapped to the WoL. The gray, dashed line indicates $y = x$. **c,** Relationship between counts of reads mapped to NCBI's Rep200 and those mapped to the WoL. The dashed line indicates $y = x$. **d**, Comparison of counts of raw reads among environments (based on EMPO 4). The two-dashed line indicates our expectation of 1 million reads from gut samples. **e,** Comparison of counts of reads mapped to the WoL among environments. **f**, Comparison of the proportion of raw reads that were mapped to the WoL among environments. For panels **d-e**, the dashed line indicates the global mean and the dotted line indicates the global median. For all panels, note the respective *y*-axis transformation used. Colors and shapes are described in the legend in panel **b**. Boxplots are in the style of Tukey, where the center line indicates the median, lower and upper hinges the first- and third quartiles, respectively, and each whisker 1.5 x the interquartile range (IQR) from its respective hinge.

### SUPPLEMENTARY NOTES

**Permits for sample collection.**

For all international sample collection, the proper procedures for sampling, exporting, and importing were followed. In accordance with the genetic resource sharing component of the Nagoya Protocol, we have made all sequence data publicly available at NCBI. Here we provide specific statements for sample collection where relevant.

For samples from Costa Rica, permits were granted by the Institutional Biodiversity Commission of the University of Costa Rica (UCR, resolution number 055-2016) and authorized by the Organization for Tropical Studies (OTS) and the Central Pacific Conservation Area (ACOPAC) of the Ministry of Energy and the Environment (MINAE), Costa Rican government, under UCR project B6-656.

For samples from Ukraine, all procedures were performed in accordance with legal requirements and regulations from Ukrainian authorities (957-i/16/05/2016), and the Animal Experiment Board in Finland (ESAVI/7256/04.10.07/2014). The samples were transported to Finland for research purposes based on the import permission from the Evira (3679/0460/2016).

Samples from Namibia were collected under the Republic of Namibia - Ministry of Mines and Energy permit number ES30246 and transported to South Africa for research purpose with import permit P0067933 from the Department of Agriculture, Forestry and Fisheries of the Republic of South Africa.

For samples from Singapore, permit (No:NP/RP18-086) was granted by National Parks Board (NParks) and sampling was conducting according to stipulations of the permit.

All coral samples were collected by AAUS-certified scientific divers, in accordance with local regulations. Relevant permit numbers are: CITES (PWS2014-AU-002155, 12US784243/9), Great Barrier Reef Marine Park Authority (G12/35236.1, G14/36788.1), Lord Howe Island Marine Park (LHIMP/R/2015/005), New South Wales Department of Primary Industries (P15/0072–1.0, OUT 15/11450), US Fish and Wildlife Service (2015LA1632527, 2015LA1703560), and Western Australia Department of Parks and Wildlife (SF010348, CE004874, ES002315).

**How to access and contribute to the EMP500.**

All methods and protocols can be accessed at www.earthmicrobiome.org and GitHub (https://github.com/biocore/emp/). All raw and processed sequence data, as well as feature-tables for all data, and sample metadata can be accessed through Qiita (www.qiita.ucsd.edu; study ID: 13114). All raw metabolomics data can be accessed through MassIVE (MSV000083475, MSV000083743). We note that in parallel to future sample collection efforts directed by the EMP Consortium, all projects adhering to the EMP standardized protocols for sample collection and sample processing can be analyzed using meta-analyses with the data provided here, and all other data generated by following those protocols. Announcements for future sample collection directed by the EMP500 Consortium will be made via https://earthmicrobiome.org.
