## Supplementary material for "Multi-omics profiling of Earth’s biomes reveals patterns of diversity and co-occurrence in microbial and metabolite composition across environments": Table S2

**Table S2 | Sample preparation summary** showing the distribution of samples among data layers generated for the EMP500. Values for sequence data layers represent read counts.

| sample name | 16S | 18S | ITS | shotgun metagenomic | lc-msms | gcms |
|---|---|---|---|---|---|---|
| 13114.angenent.65.s001 | 53525 | 103312 | 302905 | 10443561 | TRUE | TRUE |
| 13114.angenent.65.s002 | 159869 | 67791 | 266339 | 12716323 | TRUE | TRUE |
| 13114.angenent.65.s003 | 31242 | FALSE | FALSE | 1813547 | FALSE | FALSE |
| 13114.angenent.65.s004 | 37541 | FALSE | FALSE | 3711056 | FALSE | FALSE |
| 13114.angenent.65.s005 | 127807 | 135445 | 86347 | 4697336 | TRUE | TRUE |
| 13114.angenent.65.s006 | 39481 | 90884 | 33439 | 3123931 | TRUE | TRUE |
| 13114.angenent.65.s007 | 90201 | 165656 | 125655 | 13106283 | TRUE | TRUE |
| 13114.angenent.65.s008 | 33883 | FALSE | FALSE | 2356117 | FALSE | FALSE |
| 13114.angenent.65.s009 | 36237 | FALSE | FALSE | 2632055 | FALSE | FALSE |
| 13114.berry.2.s001 | 72149 | 121274 | 2003 | 12858750 | TRUE | TRUE |
| 13114.berry.2.s002 | 100353 | 89939 | 4163 | 13034781 | TRUE | TRUE |
| 13114.berry.2.s003 | 46923 | 103613 | 2933 | 13640507 | TRUE | TRUE |
| 13114.berry.2.s004 | 101912 | 247617 | 573 | 6355810 | TRUE | TRUE |
| 13114.berry.2.s005 | 83799 | 44135 | 490 | 3219287 | TRUE | TRUE |
| 13114.berry.2.s006 | 14716 | 28279 | 279 | 9615256 | TRUE | TRUE |
| 13114.berry.2.s007 | 64774 | 205835 | 13893 | 3081491 | TRUE | TRUE |
| 13114.berry.2.s008 | 162908 | 229570 | 46227 | 15823305 | TRUE | TRUE |
| 13114.berry.2.s009 | 138059 | 275862 | 8622 | 14497052 | TRUE | TRUE |
| 13114.berry.2.s010 | 139871 | 173365 | 504 | 9802326 | TRUE | TRUE |
| 13114.berry.2.s011 | 145077 | 243816 | 941 | 9211503 | TRUE | TRUE |
| 13114.berry.2.s012 | 7780 | 119326 | 724 | 3574746 | TRUE | TRUE |
| 13114.berry.2.s013 | 117105 | 125568 | 26894 | 14528843 | TRUE | TRUE |
| 13114.berry.2.s014 | 119943 | 224032 | 6458 | 14249291 | TRUE | TRUE |
| 13114.berry.2.s015 | 25255 | 151408 | 797 | 7711033 | TRUE | TRUE |
| 13114.berry.2.s016 | 104197 | 144340 | 13967 | 11955206 | TRUE | TRUE |
| 13114.berry.2.s017 | 72049 | 134822 | 3968 | 8361807 | TRUE | TRUE |
| 13114.berry.2.s018 | 31271 | 59903 | 866 | 9909138 | TRUE | TRUE |
| 13114.berry.2.s019 | 112954 | 162394 | 18 | 7398499 | TRUE | TRUE |
| 13114.berry.2.s020 | 78734 | 116100 | 147 | 14562487 | TRUE | TRUE |
| 13114.bittleston.68.s001 | 41079 | 104006 | 30635 | 3703541 | TRUE | TRUE |
| 13114.bittleston.68.s002 | 32587 | 243123 | 71092 | 11741353 | TRUE | TRUE |
| 13114.bittleston.68.s003 | 59719 | 105138 | 113078 | 11030244 | TRUE | TRUE |
| 13114.bittleston.68.s004 | 36496 | 69856 | 5757 | 11644282 | FALSE | TRUE |
| 13114.bittleston.68.s005 | 26014 | 69036 | 289 | 11639541 | TRUE | TRUE |
| 13114.bittleston.68.s006 | 35456 | 62849 | 10373 | 10136982 | TRUE | TRUE |
| 13114.bittleston.68.s007 | 32866 | 116115 | 14021 | 9990613 | TRUE | TRUE |
| 13114.bittleston.68.s008 | 42990 | 80266 | 31839 | 11451837 | TRUE | TRUE |
| 13114.bittleston.68.s009 | 23388 | 31257 | 11817 | 3363432 | TRUE | TRUE |
| 13114.bittleston.68.s010 | 88658 | 74589 | 288610 | 3577448 | TRUE | TRUE |
| 13114.bittleston.68.s011 | 21760 | 24223 | 369576 | 774604 | TRUE | TRUE |
| 13114.bittleston.68.s012 | 24581 | 50077 | 120054 | 2895010 | TRUE | TRUE |
| 13114.bittleston.68.s013 | 36854 | 88637 | 85089 | 95415 | FALSE | TRUE |

| sample name | 16S | 18S | ITS | shotgun metagenomic | lc-msms | gcms |
|---|---|---|---|---|---|---|
| 13114.bittleston.68.s014 | 8914 | 42918 | 61532 | 3081432 | TRUE | TRUE |
| 13114.bittleston.68.s015 | 11932 | 21059 | 124863 | 1517911 | TRUE | TRUE |
| 13114.bowen.74.s001 | 120392 | 109282 | 86652 | 15071594 | TRUE | TRUE |
| 13114.bowen.74.s002 | 107631 | 114908 | 34468 | 13544095 | TRUE | TRUE |
| 13114.bowen.74.s003 | 52900 | 106541 | 49602 | 11093244 | TRUE | TRUE |
| 13114.bowen.74.s004 | 98011 | 176066 | 25941 | 11756206 | TRUE | TRUE |
| 13114.bowen.74.s005 | 135568 | 125202 | 27045 | 9744319 | TRUE | TRUE |
| 13114.bowen.74.s006 | 98149 | 163951 | 51221 | 10967182 | TRUE | TRUE |
| 13114.bowen.74.s007 | 29803 | 99140 | 17786 | 10954555 | TRUE | TRUE |
| 13114.bowen.74.s008 | 62944 | 132537 | 3828 | 3308907 | TRUE | TRUE |
| 13114.bowen.74.s009 | 45254 | 129160 | 26198 | 12647235 | TRUE | TRUE |
| 13114.bowen.74.s010 | 50764 | 103706 | 25653 | 749094 | TRUE | TRUE |
| 13114.bowen.74.s011 | 90978 | 153785 | 53075 | 14528729 | TRUE | TRUE |
| 13114.bowen.74.s012 | 126504 | 185951 | 34806 | 13401542 | TRUE | TRUE |
| 13114.bowen.74.s013 | 23174 | 108022 | 12048 | 14044198 | TRUE | TRUE |
| 13114.bowen.74.s014 | 112306 | 53696 | 25190 | 10685668 | TRUE | TRUE |
| 13114.bowen.74.s015 | 81482 | 162245 | 86622 | 11835184 | TRUE | TRUE |
| 13114.bowen.74.s016 | 86368 | 137806 | 38130 | 8331544 | TRUE | TRUE |
| 13114.distel.72.s001 | 44033 | 91271 | 1583 | 11768152 | TRUE | TRUE |
| 13114.distel.72.s002 | 40610 | 74716 | 757 | 7084298 | TRUE | TRUE |
| 13114.distel.72.s003 | 7719 | 58661 | 161 | 13363923 | TRUE | TRUE |
| 13114.distel.72.s004 | 35797 | 94416 | 1099 | 12134239 | TRUE | TRUE |
| 13114.distel.72.s005 | 13613 | 187772 | 241 | 11513261 | TRUE | TRUE |
| 13114.distel.72.s006 | 14541 | 154762 | 1265 | 16879487 | TRUE | TRUE |
| 13114.distel.72.s007 | 43393 | 142448 | 3554 | 5110129 | TRUE | TRUE |
| 13114.distel.72.s008 | 55357 | 131022 | 1391 | 14658430 | TRUE | TRUE |
| 13114.distel.72.s009 | 39350 | 78089 | 1525 | 4160215 | TRUE | FALSE |
| 13114.distel.72.s010 | 17941 | 69251 | 56 | 12630666 | TRUE | TRUE |
| 13114.distel.72.s011 | 99414 | 56063 | 11383 | 13162204 | TRUE | TRUE |
| 13114.distel.72.s012 | 46035 | 123841 | 904 | 7654851 | TRUE | TRUE |
| 13114.distel.72.s013 | 80327 | 107769 | 7004 | 10200588 | TRUE | FALSE |
| 13114.distel.72.s014 | 33660 | 31803 | 2204 | 8784 | TRUE | FALSE |
| 13114.distel.72.s015 | 72122 | 48528 | 12236 | 14357614 | TRUE | FALSE |
| 13114.girguis.50.s001 | 84389 | 223903 | 27166 | 12902608 | TRUE | TRUE |
| 13114.girguis.50.s002 | 84688 | 74171 | 9947 | 13094538 | TRUE | TRUE |
| 13114.girguis.50.s003 | 39396 | 85094 | 16614 | 4742830 | TRUE | TRUE |
| 13114.girguis.50.s004 | 55798 | 105335 | 469 | 3003223 | TRUE | TRUE |
| 13114.girguis.50.s005 | 104624 | 127094 | 4853 | 12979719 | TRUE | TRUE |
| 13114.girguis.50.s006 | 21809 | 96705 | 7 | 1009737 | TRUE | TRUE |
| 13114.girguis.50.s007 | 60747 | 189002 | 23840 | 2192433 | TRUE | TRUE |
| 13114.girguis.50.s008 | 67219 | 50745 | 572 | 11311122 | TRUE | TRUE |
| 13114.girguis.50.s009 | 75337 | 100191 | 7307 | 10950380 | TRUE | TRUE |
| 13114.girguis.50.s010 | 46550 | 27211 | 449 | 5778394 | TRUE | TRUE |
| 13114.girguis.50.s011 | 83706 | 255603 | 1888 | 4302125 | TRUE | TRUE |

| sample name | 16S | 18S | ITS | shotgun metagenomic | lc-msms | gcms |
|---|---|---|---|---|---|---|
| 13114.girguis.50.s012 | 130110 | 152045 | 1280 | 9508681 | TRUE | TRUE |
| 13114.jensen.43.s001 | 85339 | 106856 | 12226 | 8054122 | TRUE | TRUE |
| 13114.jensen.43.s002 | 60073 | 76388 | 53282 | 756249 | TRUE | TRUE |
| 13114.jensen.43.s003 | 38602 | 30900 | 32 | 12173585 | TRUE | TRUE |
| 13114.jensen.43.s004 | 52662 | 189560 | 13924 | 9512844 | TRUE | TRUE |
| 13114.jensen.43.s005 | 128204 | 88957 | 80087 | 1832829 | TRUE | TRUE |
| 13114.jensen.43.s006 | 80863 | 117905 | 75099 | 12091709 | TRUE | TRUE |
| 13114.jensen.43.s007 | 52226 | 109910 | 17148 | 8660370 | TRUE | FALSE |
| 13114.jensen.43.s008 | 87475 | 204446 | 7465 | 1224977 | TRUE | TRUE |
| 13114.jensen.43.s009 | 91792 | 88122 | 5612 | 6555534 | TRUE | TRUE |
| 13114.jensen.43.s010 | 35625 | 61861 | 2165 | 12987794 | TRUE | TRUE |
| 13114.jensen.43.s011 | 75512 | 66040 | 40073 | 13198921 | TRUE | TRUE |
| 13114.jensen.43.s012 | 32084 | 69825 | 1811 | 9363635 | TRUE | TRUE |
| 13114.jensen.43.s013 | 95387 | 131845 | 766 | 2366988 | TRUE | TRUE |
| 13114.jensen.43.s014 | 85937 | 233042 | 7692 | 1385957 | TRUE | FALSE |
| 13114.jensen.43.s015 | 94284 | 236809 | 39774 | 10157405 | TRUE | TRUE |
| 13114.jensen.43.s016 | 93533 | 110935 | 14647 | 10878763 | TRUE | TRUE |
| 13114.jensen.43.s017 | 121802 | 55554 | 895 | 4907376 | TRUE | TRUE |
| 13114.jensen.43.s018 | 52546 | 84895 | 250 | 3101538 | TRUE | TRUE |
| 13114.jensen.43.s019 | 82353 | 109723 | 19807 | 11624620 | TRUE | TRUE |
| 13114.king.27.s001 | 98894 | 18720 | 132433 | 18795908 | TRUE | TRUE |
| 13114.king.27.s002 | 69488 | 70960 | 21794 | 224014 | TRUE | TRUE |
| 13114.king.27.s003 | 85952 | 88973 | 146209 | 1136926 | TRUE | TRUE |
| 13114.king.27.s004 | 103252 | 14946 | 19216 | 2922184 | TRUE | TRUE |
| 13114.king.27.s005 | 61054 | 82586 | 68011 | 4517747 | TRUE | FALSE |
| 13114.king.27.s006 | 37042 | 44850 | 14077 | 144139 | TRUE | FALSE |
| 13114.king.27.s007 | 117024 | 32260 | 57671 | 3282101 | TRUE | TRUE |
| 13114.king.27.s008 | 107105 | 37962 | 29269 | 9167860 | TRUE | TRUE |
| 13114.king.27.s009 | 46498 | 17764 | 40827 | 3180849 | TRUE | FALSE |
| 13114.king.27.s010 | 56054 | 32481 | 81706 | 4736691 | TRUE | TRUE |
| 13114.king.27.s011 | 149356 | 136478 | 101473 | 6755741 | TRUE | TRUE |
| 13114.king.27.s012 | 47159 | 96064 | 114164 | 5557420 | TRUE | TRUE |
| 13114.king.27.s013 | 133418 | 41439 | 83529 | 10892179 | TRUE | TRUE |
| 13114.king.27.s014 | 107293 | 141208 | 127413 | 4384706 | TRUE | FALSE |
| 13114.king.27.s015 | 61502 | 50253 | 18546 | 4214815 | TRUE | FALSE |
| 13114.king.27.s016 | 52568 | 332 | 52406 | 9190349 | TRUE | TRUE |
| 13114.king.27.s017 | 15882 | 87311 | 132 | 564 | TRUE | TRUE |
| 13114.king.27.s018 | 109695 | 71242 | 29618 | 2604718 | TRUE | TRUE |
| 13114.king.27.s019 | 6831 | 1 | 2 | 140 | TRUE | TRUE |
| 13114.king.27.s020 | 64956 | 58739 | 2426 | 10779701 | TRUE | FALSE |
| 13114.king.27.s021 | 9034 | 71 | 164 | 36 | TRUE | FALSE |
| 13114.Kshtrika1.misc.103 | 1824 | 23111 | 7 | FALSE | FALSE | FALSE |
| 13114.Kshtrika1.misc.1977 | 589 | 42804 | 29 | FALSE | FALSE | FALSE |
| 13114.Kshtrika1.misc.1981 | 387 | 6701 | 92 | FALSE | FALSE | FALSE |

| sample name | 16S | 18S | ITS | shotgun metagenomic | lc-msms | gcms |
|---|---|---|---|---|---|---|
| 13114.Kshtrika1.misc.2026 | 280 | FALSE | 50 | FALSE | FALSE | FALSE |
| 13114.Kshtrika1.misc.2033.P1 | 130 | 10014 | 279 | FALSE | FALSE | FALSE |
| 13114.Kshtrika1.misc.2033.P3 | 364 | 18291 | 296 | FALSE | FALSE | FALSE |
| 13114.Kshtrika1.misc.2063 | 1501 | 35746 | 20 | FALSE | FALSE | FALSE |
| 13114.Kshtrika1.misc.2065.P1 | 50 | 30517 | 591 | FALSE | FALSE | FALSE |
| 13114.Kshtrika1.misc.2065.P4 | 160 | 9169 | 9 | FALSE | FALSE | FALSE |
| 13114.Kshtrika1.misc.2069 | 158 | 23039 | 9 | FALSE | FALSE | FALSE |
| 13114.Kshtrika1.misc.360 | 2662 | 16790 | 57 | FALSE | FALSE | FALSE |
| 13114.Kshtrika1.misc.362.P1 | 15 | 66270 | 133 | FALSE | FALSE | FALSE |
| 13114.Kshtrika1.misc.362.P5 | 105 | 45185 | 38 | FALSE | FALSE | FALSE |
| 13114.Kshtrika1.misc.600 | 1951 | 1094 | 7 | FALSE | FALSE | FALSE |
| 13114.macrae.crerar.3.s001 | 13 | FALSE | 1 | FALSE | TRUE | TRUE |
| 13114.macrae.crerar.3.s002 | 17 | FALSE | 1 | FALSE | TRUE | TRUE |
| 13114.macrae.crerar.3.s003 | 17 | FALSE | FALSE | FALSE | TRUE | TRUE |
| 13114.macrae.crerar.3.s004 | 10 | FALSE | 84191 | FALSE | TRUE | TRUE |
| 13114.macrae.crerar.3.s005 | 12 | FALSE | 2 | FALSE | TRUE | TRUE |
| 13114.macrae.crerar.3.s006 | 34138 | FALSE | 1 | FALSE | TRUE | TRUE |
| 13114.macrae.crerar.3.s007 | 17 | FALSE | FALSE | FALSE | TRUE | TRUE |
| 13114.macrae.crerar.3.s008 | 47407 | FALSE | FALSE | FALSE | TRUE | TRUE |
| 13114.macrae.crerar.3.s009 | 18 | FALSE | FALSE | FALSE | TRUE | TRUE |
| 13114.macrae.crerar.3.s010 | 36828 | FALSE | FALSE | FALSE | TRUE | TRUE |
| 13114.macrae.crerar.3.s011 | 28 | FALSE | FALSE | FALSE | TRUE | TRUE |
| 13114.macrae.crerar.3.s012 | 13 | FALSE | 9013 | FALSE | TRUE | TRUE |
| 13114.makhalanyane.46.s001 | 6 | FALSE | FALSE | 348 | TRUE | TRUE |
| 13114.makhalanyane.46.s002 | 33527 | FALSE | FALSE | 484 | TRUE | TRUE |
| 13114.makhalanyane.46.s003 | 35906 | FALSE | 1 | 348 | TRUE | TRUE |
| 13114.makhalanyane.46.s004 | 15 | FALSE | 4 | 24582 | TRUE | TRUE |
| 13114.makhalanyane.46.s005 | 20 | FALSE | FALSE | 120889 | TRUE | TRUE |
| 13114.makhalanyane.46.s006 | 40297 | FALSE | FALSE | 14437 | TRUE | TRUE |
| 13114.makhalanyane.46.s007 | 44986 | FALSE | 914 | 376831 | TRUE | TRUE |
| 13114.makhalanyane.46.s008 | 21 | FALSE | 1 | 33660 | TRUE | TRUE |
| 13114.makhalanyane.46.s009 | 29169 | FALSE | FALSE | 88560 | TRUE | TRUE |
| 13114.makhalanyane.46.s010 | 48189 | FALSE | FALSE | 10114774 | TRUE | TRUE |
| 13114.makhalanyane.47.s001 | FALSE | FALSE | FALSE | 2128 | TRUE | TRUE |
| 13114.makhalanyane.47.s002 | FALSE | FALSE | FALSE | 30 | TRUE | TRUE |
| 13114.makhalanyane.47.s003 | FALSE | FALSE | 2 | 737 | TRUE | TRUE |
| 13114.makhalanyane.47.s004 | FALSE | FALSE | FALSE | 220 | TRUE | TRUE |
| 13114.makhalanyane.47.s005 | FALSE | FALSE | 1 | 156 | TRUE | TRUE |
| 13114.makhalanyane.47.s006 | FALSE | FALSE | FALSE | 23 | TRUE | TRUE |
| 13114.makhalanyane.47.s007 | FALSE | FALSE | 1 | 6705 | TRUE | TRUE |
| 13114.makhalanyane.47.s008 | FALSE | FALSE | 3 | 523 | TRUE | TRUE |
| 13114.makhalanyane.47.s009 | FALSE | FALSE | FALSE | 312 | TRUE | TRUE |
| 13114.mayer.33.s001 | 69863 | 265479 | 50602 | 13444186 | TRUE | TRUE |
| 13114.mayer.33.s002 | 99431 | 223530 | 137453 | 16526773 | TRUE | TRUE |

| sample name | 16S | 18S | ITS | shotgun metagenomic | lc-msms | gcms |
|---|---|---|---|---|---|---|
| 13114.mayer.33.s003 | 151872 | 40819 | 74853 | 6933743 | TRUE | TRUE |
| 13114.mayer.33.s004 | 48503 | 114118 | 41007 | 12830587 | TRUE | TRUE |
| 13114.mayer.33.s005 | 86941 | 212020 | 19334 | 15727225 | TRUE | TRUE |
| 13114.mayer.33.s006 | 38787 | 82526 | 65123 | 14285082 | TRUE | TRUE |
| 13114.mayer.33.s007 | 150019 | 270754 | 118506 | 14255418 | TRUE | TRUE |
| 13114.mayer.34.s001 | 114416 | 179712 | 155137 | 17268977 | TRUE | TRUE |
| 13114.mayer.34.s002 | 153983 | 149495 | 204852 | 12942118 | TRUE | TRUE |
| 13114.mayer.34.s003 | 79864 | 114621 | 97900 | 11729800 | TRUE | TRUE |
| 13114.mayer.34.s004 | 44456 | 140533 | 108408 | 6824775 | TRUE | TRUE |
| 13114.mayer.34.s005 | 35802 | 96404 | 27327 | 12399515 | TRUE | TRUE |
| 13114.mayer.34.s006 | 123774 | 62707 | 120500 | 9190552 | TRUE | TRUE |
| 13114.mayer.34.s007 | 56262 | 89350 | 10910 | 9922509 | TRUE | TRUE |
| 13114.mayer.34.s008 | 47338 | 177001 | 62259 | 13076909 | TRUE | TRUE |
| 13114.mayer.34.s009 | 107653 | 179948 | 79522 | 11803590 | TRUE | TRUE |
| 13114.mayer.34.s010 | 103536 | 45598 | 99761 | 11838031 | TRUE | TRUE |
| 13114.mcmahon.21.s001 | 24058 | 9851 | 1134 | 3025701 | FALSE | FALSE |
| 13114.mcmahon.21.s002 | 115 | 2 | 19419 | 2937668 | FALSE | FALSE |
| 13114.mcmahon.21.s003 | 27744 | 64834 | 2994 | 5099029 | FALSE | FALSE |
| 13114.mcmahon.21.s004 | 24884 | FALSE | 59439 | 2448399 | FALSE | FALSE |
| 13114.mcmahon.21.s005 | 37660 | FALSE | FALSE | 3000269 | FALSE | FALSE |
| 13114.mcmahon.21.s006 | 20218 | FALSE | 4 | 1738357 | FALSE | FALSE |
| 13114.mcmahon.21.s007 | 32920 | FALSE | 5 | 3243588 | FALSE | FALSE |
| 13114.mcmahon.21.s008 | 48574 | 133957 | 140094 | 17204065 | TRUE | FALSE |
| 13114.mcmahon.21.s009 | 114357 | 306386 | 99988 | 16094738 | TRUE | FALSE |
| 13114.mcmahon.21.s010 | 107962 | 189949 | 150322 | 17023050 | TRUE | FALSE |
| 13114.mcmahon.21.s011 | 31196 | 71316 | 1321 | 5331572 | FALSE | FALSE |
| 13114.mcmahon.21.s012 | 19068 | 66904 | 26678 | 1528421 | FALSE | FALSE |
| 13114.mcmahon.21.s013 | 22288 | 50767 | 4309 | 2943889 | FALSE | FALSE |
| 13114.mcmahon.21.s014 | 50322 | FALSE | FALSE | 2566508 | FALSE | FALSE |
| 13114.mcmahon.21.s015 | 61261 | FALSE | 43322 | 3115773 | FALSE | FALSE |
| 13114.mcmahon.21.s016 | 37939 | FALSE | FALSE | 1777102 | FALSE | FALSE |
| 13114.mcmahon.21.s017 | 38144 | FALSE | FALSE | 2703179 | FALSE | FALSE |
| 13114.metcalf.40.s001 | 95935 | 98102 | 95492 | 12550563 | TRUE | FALSE |
| 13114.metcalf.40.s002 | 101698 | 120348 | 55059 | 10218499 | TRUE | TRUE |
| 13114.metcalf.40.s003 | 44513 | 96969 | 112777 | 2623007 | TRUE | TRUE |
| 13114.metcalf.40.s004 | 35331 | 85232 | 95107 | 5864046 | TRUE | FALSE |
| 13114.metcalf.40.s005 | 103233 | 111652 | 108539 | 11267455 | TRUE | FALSE |
| 13114.metcalf.40.s006 | 155838 | 206351 | 163061 | 13031156 | TRUE | TRUE |
| 13114.metcalf.40.s007 | 71240 | 90305 | 61496 | 9777007 | TRUE | TRUE |
| 13114.metcalf.40.s008 | 79391 | 85963 | 38930 | 10516541 | TRUE | FALSE |
| 13114.metcalf.40.s009 | 90101 | 90369 | 88751 | 12846752 | TRUE | FALSE |
| 13114.metcalf.40.s010 | 73814 | 130178 | 86784 | 12159349 | TRUE | TRUE |
| 13114.metcalf.40.s011 | 105590 | 55264 | 50450 | 13379692 | TRUE | TRUE |
| 13114.metcalf.40.s012 | 45862 | 92228 | 32857 | 13627895 | TRUE | FALSE |

| sample name | 16S | 18S | ITS | shotgun metagenomic | lc-msms | gcms |
|---|---|---|---|---|---|---|
| 13114.metcalf.40.s013 | 76283 | 55224 | 204357 | 12287519 | TRUE | TRUE |
| 13114.metcalf.40.s014 | 26371 | 32868 | 40791 | 8290897 | TRUE | TRUE |
| 13114.metcalf.81.s001 | 53979 | 195325 | 194554 | 14764764 | TRUE | TRUE |
| 13114.metcalf.81.s002 | 118741 | 117939 | 107651 | 11608813 | TRUE | TRUE |
| 13114.metcalf.81.s003 | 108099 | 87369 | 38472 | 14987870 | TRUE | TRUE |
| 13114.minich.76.s001 | 102331 | 145844 | 70522 | 12659405 | TRUE | TRUE |
| 13114.minich.76.s002 | 68551 | 218874 | 60593 | 18544183 | TRUE | TRUE |
| 13114.minich.76.s003 | 107670 | 147721 | 103972 | 12733176 | TRUE | TRUE |
| 13114.minich.76.s004 | 80459 | 89473 | 117309 | 3638612 | TRUE | TRUE |
| 13114.minich.76.s005 | 74540 | 193950 | 84561 | 12433199 | TRUE | TRUE |
| 13114.minich.76.s006 | 82410 | 263306 | 192637 | 10254311 | TRUE | TRUE |
| 13114.mousseau.88.s001 | 141011 | 78480 | 44789 | 7417403 | FALSE | TRUE |
| 13114.mousseau.88.s002 | 119364 | 26210 | 33420 | 1983796 | TRUE | FALSE |
| 13114.mousseau.88.s003 | 43821 | 22720 | 2630 | 4135265 | FALSE | TRUE |
| 13114.mousseau.88.s004 | 63374 | 21607 | 44955 | 68990 | TRUE | FALSE |
| 13114.mousseau.88.s005 | 94741 | 42564 | 58787 | 5374869 | TRUE | FALSE |
| 13114.mousseau.88.s006 | 108784 | 5692 | 50 | 3362020 | TRUE | FALSE |
| 13114.mousseau.88.s007 | 99356 | 1909 | 1739 | 2138357 | TRUE | FALSE |
| 13114.mousseau.88.s008 | 25127 | 24397 | 16587 | 6070729 | FALSE | TRUE |
| 13114.mousseau.88.s009 | 70810 | 1154 | 973 | 3882824 | TRUE | FALSE |
| 13114.mousseau.88.s010 | 64173 | 24504 | 5611 | 1388356 | TRUE | FALSE |
| 13114.mousseau.88.s011 | 94707 | 63808 | 40079 | 49876414 | TRUE | FALSE |
| 13114.mousseau.88.s012 | 32262 | FALSE | FALSE | 175768 | FALSE | TRUE |
| 13114.mousseau.88.s013 | 38320 | FALSE | 1 | 133364 | TRUE | FALSE |
| 13114.mousseau.88.s014 | 46165 | FALSE | FALSE | 5681581 | FALSE | TRUE |
| 13114.mousseau.88.s015 | 39052 | FALSE | 1 | 89383 | TRUE | FALSE |
| 13114.mousseau.88.s016 | 60522 | 34084 | 50468 | 58571908 | TRUE | FALSE |
| 13114.mousseau.88.s017 | 65417 | 2429 | 961 | 1795468 | TRUE | FALSE |
| 13114.mousseau.88.s018 | 41438 | 479 | 416 | 8685 | TRUE | FALSE |
| 13114.mousseau.88.s019 | 54552 | FALSE | 20 | 94449628 | FALSE | TRUE |
| 13114.mousseau.88.s020 | 44022 | FALSE | 1 | 195168 | FALSE | TRUE |
| 13114.mousseau.88.s021 | 31048 | FALSE | 1 | 4041393 | TRUE | FALSE |
| 13114.mousseau.88.s022 | 43224 | FALSE | 1 | 63971 | FALSE | TRUE |
| 13114.mousseau.88.s023 | 124242 | 71728 | 32051 | 11685886 | TRUE | FALSE |
| 13114.mousseau.88.s024 | 31881 | FALSE | FALSE | 1352957 | TRUE | FALSE |
| 13114.mousseau.88.s025 | 58504 | FALSE | FALSE | 2029863 | TRUE | FALSE |
| 13114.mousseau.88.s026 | 69757 | 1844 | 1235 | 1769294 | FALSE | TRUE |
| 13114.mousseau.88.s027 | 15073 | FALSE | 2 | 1438875 | FALSE | TRUE |
| 13114.mousseau.88.s028 | 59215 | FALSE | 3 | 875864 | TRUE | FALSE |
| 13114.mousseau.88.s029 | 93088 | 21993 | 14941 | 26531035 | FALSE | TRUE |
| 13114.mousseau.88.s030 | 55998 | FALSE | 1 | 8754403 | TRUE | FALSE |
| 13114.mousseau.88.s031 | 81191 | 57247 | 1679 | 1022989 | FALSE | TRUE |
| 13114.mousseau.88.s032 | 40160 | FALSE | 1 | 8543459 | TRUE | FALSE |
| 13114.mousseau.88.s033 | 72388 | FALSE | 7 | 130535 | TRUE | FALSE |

| sample name | 16S | 18S | ITS | shotgun metagenomic | lc-msms | gcms |
|---|---|---|---|---|---|---|
| 13114.mousseau.88.s034 | 29869 | FALSE | FALSE | 345096 | TRUE | FALSE |
| 13114.mousseau.88.s035 | 64779 | 9206 | 40920 | 4209360 | TRUE | FALSE |
| 13114.mousseau.88.s036 | 81686 | 18326 | 20968 | 1839822 | TRUE | FALSE |
| 13114.mousseau.88.s037 | 166209 | 53255 | 6923 | 5099535 | TRUE | FALSE |
| 13114.mousseau.88.s038 | 39909 | FALSE | 1 | FALSE | FALSE | FALSE |
| 13114.mousseau.88.s039 | 74246 | 17585 | 5954 | 3465024 | TRUE | FALSE |
| 13114.mousseau.88.s040 | 119818 | 41124 | 152 | 2391515 | TRUE | FALSE |
| 13114.mousseau.88.s041 | 113671 | 22344 | 3552 | 4435099 | TRUE | FALSE |
| 13114.mousseau.88.s042 | 117929 | 20594 | 24585 | 1999803 | TRUE | FALSE |
| 13114.mousseau.88.s043 | 83934 | 111 | 4284 | 142227 | TRUE | FALSE |
| 13114.mousseau.88.s044 | 86241 | 55410 | 6809 | 862410 | TRUE | FALSE |
| 13114.mousseau.88.s045 | 70128 | 17871 | 849 | 7339004 | TRUE | FALSE |
| 13114.mousseau.88.s046 | 123930 | 36399 | 21458 | 4946467 | TRUE | FALSE |
| 13114.mousseau.88.s047 | 81582 | 51007 | 55149 | 4860566 | TRUE | FALSE |
| 13114.mousseau.88.s048 | 61242 | 394 | 2977 | 177166 | TRUE | FALSE |
| 13114.mousseau.88.s049 | 81786 | 66443 | 30225 | 4514002 | TRUE | FALSE |
| 13114.mousseau.88.s050 | 57523 | 15686 | 46265 | 6770295 | TRUE | FALSE |
| 13114.mousseau.88.s051 | 102790 | 1875 | 1568 | 740096 | TRUE | FALSE |
| 13114.mousseau.88.s052 | 136301 | 78719 | 47161 | 6312396 | TRUE | FALSE |
| 13114.mousseau.88.s053 | 103574 | 56364 | 70694 | 4712425 | TRUE | FALSE |
| 13114.mousseau.88.s054 | 86029 | 32 | 5178 | 2436821 | TRUE | FALSE |
| 13114.mousseau.88.s055 | 100098 | 54401 | 27063 | 4601219 | TRUE | FALSE |
| 13114.mousseau.88.s056 | 65441 | 33544 | 32921 | 4980586 | TRUE | FALSE |
| 13114.mousseau.88.s057 | 141485 | 50487 | 29945 | 6288063 | TRUE | FALSE |
| 13114.mousseau.88.s058 | 92509 | 22539 | 525 | 5190588 | TRUE | FALSE |
| 13114.mousseau.88.s059 | 29165 | 8775 | 26197 | 1858738 | TRUE | FALSE |
| 13114.mousseau.88.s060 | 36246 | FALSE | 3 | FALSE | FALSE | FALSE |
| 13114.mousseau.88.s061 | 88933 | 648 | 2899 | 1457698 | TRUE | FALSE |
| 13114.mousseau.88.s062 | 76599 | 1025 | 11372 | 4374260 | TRUE | FALSE |
| 13114.mousseau.88.s063 | 56036 | 17616 | 15486 | 4187445 | TRUE | FALSE |
| 13114.mousseau.88.s064 | 94674 | 38216 | 2495 | 3304057 | TRUE | FALSE |
| 13114.mousseau.88.s065 | 65383 | 30412 | 6760 | 9224679 | TRUE | FALSE |
| 13114.mousseau.88.s066 | 103551 | 50030 | 11979 | 2799241 | TRUE | FALSE |
| 13114.mousseau.88.s067 | 62336 | 28062 | 138 | 60019 | TRUE | FALSE |
| 13114.mousseau.88.s068 | 110881 | 57857 | 54664 | 3475541 | TRUE | FALSE |
| 13114.mousseau.88.s069 | 107327 | 53786 | 27743 | 5441086 | TRUE | FALSE |
| 13114.mousseau.88.s070 | 61839 | 1422 | 27 | 3225813 | TRUE | FALSE |
| 13114.mousseau.88.s071 | 77600 | 12640 | 499 | 166240 | TRUE | FALSE |
| 13114.mousseau.88.s072 | 48461 | 15533 | 34576 | 497946 | FALSE | TRUE |
| 13114.mousseau.88.s073 | 107184 | 37709 | 801 | 2935837 | TRUE | FALSE |
| 13114.mousseau.88.s074 | 97787 | 9323 | 1593 | 3196598 | TRUE | FALSE |
| 13114.mousseau.88.s075 | 115527 | 3543 | 6581 | 1415219 | FALSE | TRUE |
| 13114.mousseau.88.s076 | 97641 | 40664 | 23569 | 4408616 | FALSE | TRUE |
| 13114.mousseau.88.s077 | 122279 | 93476 | 35201 | 199040 | FALSE | TRUE |

| sample name | 16S | 18S | ITS | shotgun metagenomic | lc-msms | gcms |
|---|---|---|---|---|---|---|
| 13114.mousseau.88.s078 | 44949 | 597 | 424 | 2710644 | FALSE | TRUE |
| 13114.mousseau.88.s079 | 71544 | 52885 | 36761 | 6247365 | FALSE | TRUE |
| 13114.mousseau.88.s080 | 58759 | 7651 | 6171 | 3635029 | TRUE | FALSE |
| 13114.mousseau.88.s081 | 100251 | 72143 | 20387 | 6367680 | TRUE | FALSE |
| 13114.mousseau.88.s082 | 86825 | 11 | 22 | 64424 | TRUE | FALSE |
| 13114.mousseau.88.s083 | 114752 | 24467 | 33908 | 2968321 | TRUE | FALSE |
| 13114.mousseau.88.s084 | 71103 | 45582 | 3909 | 7406529 | TRUE | FALSE |
| 13114.mousseau.88.s085 | 64196 | 93 | 745 | 1286583 | TRUE | FALSE |
| 13114.mousseau.88.s086 | 123584 | 47644 | 59897 | 4961497 | TRUE | FALSE |
| 13114.mousseau.88.s087 | 96656 | 63454 | 8795 | 5387734 | TRUE | FALSE |
| 13114.mousseau.88.s088 | 124711 | 1827 | 2669 | 2606380 | TRUE | FALSE |
| 13114.mousseau.88.s089 | 94966 | 59 | 1286 | 2975180 | TRUE | FALSE |
| 13114.mousseau.88.s090 | 53253 | 39777 | 602 | 3419726 | TRUE | FALSE |
| 13114.mousseau.88.s091 | 137367 | 8774 | 2164 | 3444713 | TRUE | FALSE |
| 13114.mousseau.88.s092 | 79348 | 27787 | 94 | 2649264 | TRUE | FALSE |
| 13114.mousseau.88.s093 | 34321 | 20702 | 510 | 3554868 | TRUE | FALSE |
| 13114.mousseau.88.s094 | 114657 | 5202 | 7033 | 4172415 | TRUE | FALSE |
| 13114.mousseau.88.s095 | 108612 | 21221 | 11072 | 3043613 | TRUE | FALSE |
| 13114.mousseau.88.s096 | 110032 | 27759 | 26164 | 3708948 | TRUE | FALSE |
| 13114.mousseau.88.s097 | 60415 | 24557 | 1207 | 10491189 | TRUE | FALSE |
| 13114.mousseau.88.s098 | 47141 | 15981 | 411 | 4804844 | TRUE | FALSE |
| 13114.mousseau.88.s099 | 98915 | 7286 | 17014 | 4756004 | TRUE | FALSE |
| 13114.mousseau.88.s100 | 87139 | 3898 | 582 | 3112012 | TRUE | FALSE |
| 13114.mousseau.88.s101 | 77913 | 16905 | 3 | 3863700 | TRUE | FALSE |
| 13114.mousseau.88.s102 | 89450 | 21613 | 1000 | 2482347 | TRUE | FALSE |
| 13114.mousseau.88.s103 | 75392 | 2006 | 1569 | 1130546 | TRUE | FALSE |
| 13114.mousseau.88.s104 | 46821 | 21926 | 58713 | 3286673 | TRUE | FALSE |
| 13114.mousseau.88.s105 | 115672 | 33757 | 55519 | 4222842 | TRUE | FALSE |
| 13114.mousseau.88.s106 | 86855 | 20163 | 1112 | 3001801 | TRUE | FALSE |
| 13114.mousseau.88.s107 | 98414 | 43030 | 85977 | 6825320 | TRUE | FALSE |
| 13114.mousseau.88.s108 | 105031 | 54420 | 42497 | 5733968 | TRUE | FALSE |
| 13114.mousseau.88.s109 | 75797 | 10591 | 1014 | 2581813 | TRUE | FALSE |
| 13114.mousseau.88.s110 | 53711 | 26862 | 55516 | 1878333 | TRUE | FALSE |
| 13114.mousseau.88.s111 | 66561 | 16922 | 78461 | 395914 | TRUE | FALSE |
| 13114.mousseau.88.s112 | 30116 | 39101 | 365 | 1579345 | TRUE | FALSE |
| 13114.myrold.5.s001 | 13 | FALSE | 2 | 12678069 | FALSE | FALSE |
| 13114.myrold.5.s002 | 26 | FALSE | FALSE | 16459513 | FALSE | FALSE |
| 13114.myrold.5.s003 | 21 | FALSE | 3 | 87814387 | FALSE | FALSE |
| 13114.myrold.5.s004 | 59741 | FALSE | 4 | 11815531 | FALSE | FALSE |
| 13114.myrold.5.s005 | 10 | FALSE | FALSE | 20689181 | FALSE | FALSE |
| 13114.myrold.5.s006 | 13 | FALSE | FALSE | 16364918 | FALSE | FALSE |
| 13114.myrold.5.s007 | 55660 | FALSE | 1 | 18470170 | FALSE | FALSE |
| 13114.myrold.5.s008 | 10 | FALSE | FALSE | 53310850 | FALSE | FALSE |
| 13114.myrold.5.s009 | 8 | FALSE | 1 | 12337539 | FALSE | FALSE |

| sample name | 16S | 18S | ITS | shotgun metagenomic | lc-msms | gcms |
|---|---|---|---|---|---|---|
| 13114.myrold.5.s010 | 36333 | FALSE | 47374 | 18714948 | FALSE | FALSE |
| 13114.myrold.5.s011 | 36938 | FALSE | 1 | 11311155 | FALSE | FALSE |
| 13114.myrold.5.s012 | 34272 | FALSE | FALSE | 9289385 | FALSE | FALSE |
| 13114.myrold.59.s001 | 30 | FALSE | 1 | 9023580 | FALSE | FALSE |
| 13114.myrold.59.s002 | 30227 | FALSE | FALSE | 26130165 | TRUE | TRUE |
| 13114.myrold.59.s003 | 12 | FALSE | FALSE | 5749281 | TRUE | TRUE |
| 13114.myrold.59.s004 | 9 | FALSE | FALSE | 20782833 | TRUE | TRUE |
| 13114.myrold.59.s005 | 40882 | FALSE | FALSE | 8249852 | TRUE | TRUE |
| 13114.myrold.59.s006 | 45291 | FALSE | 1 | 12356427 | FALSE | FALSE |
| 13114.myrold.59.s007 | 13 | FALSE | 1 | 13110421 | FALSE | FALSE |
| 13114.myrold.59.s008 | 31212 | FALSE | FALSE | 920711 | TRUE | TRUE |
| 13114.myrold.59.s009 | 41193 | FALSE | 3 | 2170277 | TRUE | TRUE |
| 13114.myrold.59.s010 | 37661 | FALSE | 60526 | 10854069 | FALSE | FALSE |
| 13114.myrold.59.s011 | 42584 | FALSE | FALSE | 19353226 | TRUE | TRUE |
| 13114.myrold.59.s012 | 8 | FALSE | 1 | 1974 | TRUE | TRUE |
| 13114.myrold.59.s013 | 32511 | FALSE | FALSE | 41736 | FALSE | FALSE |
| 13114.myrold.59.s014 | 17 | FALSE | 1 | 15498 | TRUE | TRUE |
| 13114.myrold.59.s015 | 34143 | FALSE | 37810 | 16831721 | FALSE | FALSE |
| 13114.myrold.59.s016 | 25 | FALSE | 1 | 619822 | TRUE | TRUE |
| 13114.myrold.59.s017 | 66115 | FALSE | FALSE | 6480557 | FALSE | FALSE |
| 13114.myrold.59.s018 | 19606 | FALSE | 1 | 100 | TRUE | TRUE |
| 13114.myrold.59.s019 | 23 | FALSE | 30889 | 4572161 | FALSE | FALSE |
| 13114.myrold.59.s020 | 73001 | FALSE | 5 | 3017256 | FALSE | TRUE |
| 13114.myrold.59.s021 | 32916 | FALSE | FALSE | 3759473 | FALSE | FALSE |
| 13114.myrold.59.s022 | 35258 | FALSE | FALSE | 6509543 | TRUE | FALSE |
| 13114.palenik.42.s001 | 45794 | 156591 | 79816 | 13257086 | TRUE | FALSE |
| 13114.palenik.42.s002 | 100785 | 197165 | 60073 | 15578790 | TRUE | FALSE |
| 13114.palenik.42.s003 | 88950 | 179534 | 66166 | 11758795 | TRUE | FALSE |
| 13114.palenik.42.s004 | 89018 | 141126 | 8797 | 163752 | TRUE | FALSE |
| 13114.palenik.42.s005 | 56368 | 126618 | 86580 | 3042936 | TRUE | FALSE |
| 13114.palenik.42.s006 | 61758 | 65354 | 109235 | 10011400 | TRUE | FALSE |
| 13114.palenik.42.s007 | 66305 | 109055 | 55666 | 13224689 | TRUE | FALSE |
| 13114.palenik.42.s008 | 106172 | 140827 | 49247 | 13913856 | TRUE | FALSE |
| 13114.palenik.42.s009 | 80815 | 71431 | 75825 | 13035061 | TRUE | FALSE |
| 13114.palenik.42.s010 | 50660 | 83918 | 130176 | 11527516 | TRUE | FALSE |
| 13114.palenik.42.s011 | 46910 | 132652 | 90084 | 14427981 | TRUE | FALSE |
| 13114.palenik.42.s012 | 101948 | 169882 | 6734 | 9553793 | TRUE | FALSE |
| 13114.palenik.42.s013 | 36090 | 128632 | 33316 | 8975350 | TRUE | FALSE |
| 13114.palenik.42.s014 | 44700 | 74363 | 12049 | 319508 | TRUE | FALSE |
| 13114.palenik.42.s015 | 74260 | 109704 | 69794 | 10787986 | TRUE | FALSE |
| 13114.palenik.42.s016 | 24125 | 80417 | 26121 | 2606336 | FALSE | FALSE |
| 13114.palenik.42.s017 | 86906 | 42727 | 23555 | 9177262 | TRUE | FALSE |
| 13114.palenik.42.s018 | 86256 | 188967 | 39694 | 13682465 | TRUE | FALSE |
| 13114.palenik.42.s019 | 46171 | 184608 | 75173 | 12247040 | TRUE | FALSE |

| sample name | 16S | 18S | ITS | shotgun metagenomic | lc-msms | gcms |
|---|---|---|---|---|---|---|
| 13114.palenik.42.s020 | 84713 | 149513 | 192022 | 12651970 | TRUE | FALSE |
| 13114.pinto.62.s001 | FALSE | FALSE | FALSE | FALSE | TRUE | FALSE |
| 13114.pinto.62.s002 | FALSE | FALSE | FALSE | FALSE | TRUE | FALSE |
| 13114.pinto.62.s003 | FALSE | FALSE | FALSE | FALSE | TRUE | FALSE |
| 13114.pinto.62.s004 | FALSE | FALSE | FALSE | FALSE | TRUE | FALSE |
| 13114.pinto.62.s005 | FALSE | FALSE | FALSE | FALSE | TRUE | FALSE |
| 13114.pinto.62.s006 | FALSE | FALSE | FALSE | FALSE | TRUE | FALSE |
| 13114.pinto.62.s007 | FALSE | FALSE | FALSE | FALSE | TRUE | FALSE |
| 13114.pinto.62.s008 | FALSE | FALSE | FALSE | FALSE | TRUE | FALSE |
| 13114.pinto.62.s009 | FALSE | FALSE | FALSE | FALSE | TRUE | FALSE |
| 13114.pinto.62.s010 | FALSE | FALSE | FALSE | FALSE | TRUE | FALSE |
| 13114.pinto.62.s011 | FALSE | FALSE | FALSE | FALSE | TRUE | FALSE |
| 13114.pinto.62.s012 | FALSE | FALSE | FALSE | FALSE | TRUE | FALSE |
| 13114.pinto.62.s013 | FALSE | FALSE | FALSE | FALSE | TRUE | FALSE |
| 13114.pinto.62.s014 | FALSE | FALSE | FALSE | FALSE | TRUE | FALSE |
| 13114.pinto.62.s015 | FALSE | FALSE | FALSE | FALSE | TRUE | FALSE |
| 13114.pinto.62.s016 | FALSE | FALSE | FALSE | FALSE | TRUE | FALSE |
| 13114.pinto.62.s017 | FALSE | FALSE | FALSE | FALSE | TRUE | FALSE |
| 13114.pinto.62.s018 | FALSE | FALSE | FALSE | FALSE | TRUE | FALSE |
| 13114.pinto.62.s019 | FALSE | FALSE | FALSE | FALSE | TRUE | FALSE |
| 13114.pinto.62.s020 | FALSE | FALSE | FALSE | FALSE | TRUE | FALSE |
| 13114.pinto.62.s021 | FALSE | FALSE | FALSE | FALSE | TRUE | FALSE |
| 13114.pinto.62.s022 | FALSE | FALSE | FALSE | FALSE | TRUE | FALSE |
| 13114.pinto.62.s023 | FALSE | FALSE | FALSE | FALSE | TRUE | FALSE |
| 13114.pinto.62.s024 | FALSE | FALSE | FALSE | FALSE | TRUE | FALSE |
| 13114.pinto.63.s001 | 71961 | 78850 | 103062 | 8144131 | TRUE | TRUE |
| 13114.pinto.63.s002 | 15781 | 26659 | 60364 | 3613239 | TRUE | TRUE |
| 13114.pinto.63.s003 | 38073 | 48119 | 128480 | 5621559 | TRUE | TRUE |
| 13114.pinto.63.s004 | 91724 | 25914 | 63948 | 12018650 | TRUE | TRUE |
| 13114.pinto.63.s005 | 83242 | 57593 | 23898 | 10599870 | TRUE | TRUE |
| 13114.pinto.63.s006 | 87592 | 55333 | 74674 | 12756952 | TRUE | TRUE |
| 13114.pinto.63.s007 | 65689 | 74009 | 102303 | 14978642 | TRUE | FALSE |
| 13114.pinto.63.s008 | 53006 | 39161 | 71329 | 337922 | TRUE | FALSE |
| 13114.pinto.63.s009 | 75788 | 29148 | 28609 | 11140681 | TRUE | TRUE |
| 13114.pinto.63.s010 | 49475 | 27674 | 42024 | 2862929 | TRUE | TRUE |
| 13114.pinto.63.s011 | 106055 | 77779 | 59056 | 12104644 | TRUE | FALSE |
| 13114.pinto.63.s012 | 24590 | 31567 | 49000 | 11668040 | TRUE | FALSE |
| 13114.pinto.63.s013 | 70632 | 119098 | 100804 | 10364570 | TRUE | TRUE |
| 13114.pinto.63.s014 | 89422 | 82839 | 93240 | 7992491 | TRUE | TRUE |
| 13114.pinto.63.s015 | 75422 | 65678 | 79725 | 11333303 | TRUE | FALSE |
| 13114.pinto.63.s016 | 31263 | 22488 | 45001 | 13947350 | TRUE | FALSE |
| 13114.pinto.63.s017 | 83251 | 1 | 55514 | 13747978 | TRUE | FALSE |
| 13114.pinto.63.s018 | 34672 | 1 | 51442 | 13260282 | TRUE | FALSE |
| 13114.pinto.63.s019 | 122261 | 87360 | 67554 | 9613892 | TRUE | TRUE |

| sample name | 16S | 18S | ITS | shotgun metagenomic | lc-msms | gcms |
|---|---|---|---|---|---|---|
| 13114.pinto.63.s020 | 68902 | 131111 | 125128 | 11585217 | TRUE | TRUE |
| 13114.pinto.63.s021 | 104832 | 91839 | 134786 | 2007858 | TRUE | TRUE |
| 13114.pinto.63.s022 | 70027 | 106546 | 99651 | 3168225 | TRUE | TRUE |
| 13114.pinto.63.s023 | 59445 | 89496 | 97046 | 13213579 | TRUE | TRUE |
| 13114.pinto.63.s024 | 98433 | 124887 | 116467 | 11649482 | TRUE | TRUE |
| 13114.pinto.63.s025 | 131879 | 35358 | 84506 | 9003199 | TRUE | TRUE |
| 13114.pinto.63.s026 | 174334 | 79498 | 95387 | 2617674 | TRUE | FALSE |
| 13114.pinto.63.s027 | 91934 | 57511 | 72617 | 13175180 | TRUE | FALSE |
| 13114.pinto.63.s028 | 151677 | 58025 | 145827 | 11227910 | TRUE | FALSE |
| 13114.rohwer.84.s001 | 26629 | 27699 | 28482 | 3925558 | FALSE | FALSE |
| 13114.rohwer.84.s002 | 22725 | 96249 | 70926 | 2260570 | FALSE | FALSE |
| 13114.rohwer.84.s003 | 21773 | 85892 | 13583 | 1216087 | FALSE | FALSE |
| 13114.rohwer.84.s004 | 20404 | 36744 | 1744 | 2838117 | FALSE | FALSE |
| 13114.rohwer.84.s005 | 27 | 130874 | 2 | 13245915 | TRUE | TRUE |
| 13114.rohwer.84.s006 | 40 | 54149 | FALSE | 8573422 | TRUE | TRUE |
| 13114.rohwer.84.s006.rep1 | FALSE | FALSE | FALSE | FALSE | FALSE | FALSE |
| 13114.rohwer.84.s007 | 72140 | 127658 | 172362 | 11785447 | TRUE | TRUE |
| 13114.rohwer.84.s008 | 41635 | 60557 | 34881 | 10317304 | TRUE | TRUE |
| 13114.rohwer.84.s009 | 5122 | 10148 | 192 | FALSE | FALSE | FALSE |
| 13114.rohwer.84.s010 | 2928 | 29169 | 484 | FALSE | FALSE | FALSE |
| 13114.rohwer.84.s011 | 2795 | 13486 | 154 | FALSE | FALSE | FALSE |
| 13114.rohwer.84.s012 | 11247 | 69300 | 16375 | FALSE | FALSE | FALSE |
| 13114.rohwer.84.s013 | 5841 | 79455 | 8913 | FALSE | FALSE | FALSE |
| 13114.rohwer.84.s014 | 1562 | 13201 | 11296 | 834365 | FALSE | FALSE |
| 13114.rohwer.84.s015 | 446 | 42887 | 336 | FALSE | FALSE | FALSE |
| 13114.rohwer.85.s001 | 34903 | 50013 | 1580 | 5266907 | TRUE | TRUE |
| 13114.rohwer.85.s002 | 88108 | 69702 | 68868 | 12773039 | TRUE | TRUE |
| 13114.rohwer.85.s003 | 33028 | 1585 | 785 | 13413460 | TRUE | TRUE |
| 13114.rohwer.85.s004 | 59273 | 23211 | 5265 | 11479819 | TRUE | TRUE |
| 13114.rohwer.85.s005 | 22103 | 34169 | 26545 | 703368 | TRUE | FALSE |
| 13114.rohwer.85.s006 | 86238 | 58191 | 55941 | 7166381 | TRUE | FALSE |
| 13114.rohwer.85.s007 | 57967 | 97412 | 2652 | 7028354 | TRUE | FALSE |
| 13114.rohwer.85.s008 | 48334 | 44069 | 41232 | 345784 | TRUE | FALSE |
| 13114.rohwer.85.s009 | 8568 | 39743 | 974 | 66 | TRUE | FALSE |
| 13114.rohwer.85.s010 | 62017 | 15493 | 12812 | 75758 | TRUE | FALSE |
| 13114.rohwer.85.s011 | 26288 | 62668 | 7589 | 6740632 | TRUE | FALSE |
| 13114.rohwer.85.s012 | 12935 | 62495 | 19752 | 74294 | TRUE | FALSE |
| 13114.rohwer.85.s013 | 18106 | 26791 | 5789 | 43464 | TRUE | FALSE |
| 13114.rohwer.85.s014 | 2479 | 17 | 404 | 32217 | TRUE | FALSE |
| 13114.rohwer.85.s015 | 6110 | 28811 | 23877 | 272 | TRUE | FALSE |
| 13114.rohwer.85.s016 | 39278 | 37452 | 8109 | 12559778 | TRUE | FALSE |
| 13114.rohwer.85.s017 | 12110 | 23201 | 25 | 19713 | TRUE | FALSE |
| 13114.rohwer.85.s018 | 15116 | 18311 | 271 | 88194 | TRUE | FALSE |
| 13114.rohwer.85.s019 | 83909 | 47921 | 35048 | 12981532 | TRUE | FALSE |

| sample name | 16S | 18S | ITS | shotgun metagenomic | lc-msms | gcms |
|---|---|---|---|---|---|---|
| 13114.rohwer.85.s020 | 59835 | 62334 | 9382 | 7555617 | TRUE | FALSE |
| 13114.rohwer.85.s021 | 61772 | 84715 | 45527 | 697893 | TRUE | TRUE |
| 13114.rohwer.85.s022 | 51218 | 42634 | 100358 | 10442543 | TRUE | TRUE |
| 13114.rohwer.85.s023 | 23540 | 49161 | 30528 | 55263 | TRUE | TRUE |
| 13114.rohwer.85.s024 | 22118 | 55146 | 27170 | 1001880 | TRUE | TRUE |
| 13114.rohwer.85.s025 | 36473 | 50497 | 87330 | 56838 | TRUE | FALSE |
| 13114.rohwer.85.s026 | 2974 | 77152 | 36073 | 189915 | TRUE | FALSE |
| 13114.rohwer.85.s027 | 23060 | 56424 | 18020 | 489722 | TRUE | FALSE |
| 13114.rohwer.86.s001 | 91972 | 130993 | 1083 | 12447947 | TRUE | TRUE |
| 13114.rohwer.86.s002 | 38247 | 154023 | 694 | 9428241 | TRUE | TRUE |
| 13114.rohwer.86.s003 | 99645 | 104308 | 666 | 2768071 | TRUE | TRUE |
| 13114.roy.chowdhury.45.s001 | 38911 | FALSE | 20223 | FALSE | FALSE | FALSE |
| 13114.roy.chowdhury.45.s002 | 43554 | FALSE | 1 | FALSE | FALSE | FALSE |
| 13114.roy.chowdhury.45.s003 | 43288 | FALSE | 37958 | FALSE | FALSE | FALSE |
| 13114.roy.chowdhury.45.s004 | 41720 | FALSE | FALSE | FALSE | FALSE | FALSE |
| 13114.roy.chowdhury.45.s005 | 6 | FALSE | FALSE | FALSE | FALSE | FALSE |
| 13114.roy.chowdhury.45.s006 | 87 | FALSE | 2 | FALSE | FALSE | FALSE |
| 13114.roy.chowdhury.45.s007 | 53 | FALSE | FALSE | FALSE | FALSE | FALSE |
| 13114.roy.chowdhury.45.s008 | 32645 | FALSE | 3 | FALSE | FALSE | FALSE |
| 13114.roy.chowdhury.45.s009 | 66 | FALSE | 1 | FALSE | FALSE | FALSE |
| 13114.roy.chowdhury.45.s010 | 18 | FALSE | 1 | FALSE | FALSE | FALSE |
| 13114.roy.chowdhury.45.s011 | 25 | FALSE | 52410 | FALSE | FALSE | FALSE |
| 13114.roy.chowdhury.45.s012 | 39301 | FALSE | 2 | FALSE | FALSE | FALSE |
| 13114.roy.chowdhury.45.s013 | 11 | FALSE | FALSE | FALSE | FALSE | FALSE |
| 13114.roy.chowdhury.45.s014 | 11 | FALSE | 28696 | FALSE | FALSE | FALSE |
| 13114.roy.chowdhury.45.s015 | 24 | FALSE | 5 | FALSE | FALSE | FALSE |
| 13114.roy.chowdhury.45.s016 | 27200 | FALSE | 4 | FALSE | FALSE | FALSE |
| 13114.roy.chowdhury.45.s017 | FALSE | FALSE | 3 | FALSE | FALSE | FALSE |
| 13114.roy.chowdhury.45.s018 | 45379 | FALSE | FALSE | FALSE | FALSE | FALSE |
| 13114.roy.chowdhury.45.s019 | 37136 | FALSE | 1 | FALSE | FALSE | FALSE |
| 13114.roy.chowdhury.45.s020 | 11 | FALSE | 1 | FALSE | FALSE | FALSE |
| 13114.roy.chowdhury.45.s021 | 36126 | FALSE | 52026 | FALSE | FALSE | FALSE |
| 13114.roy.chowdhury.45.s022 | 47504 | FALSE | FALSE | FALSE | FALSE | FALSE |
| 13114.roy.chowdhury.45.s023 | 18 | FALSE | 1 | FALSE | FALSE | FALSE |
| 13114.roy.chowdhury.45.s024 | 54661 | FALSE | 169 | FALSE | FALSE | FALSE |
| 13114.sandin.54.s001 | 3590 | 152956 | 368 | 75003 | TRUE | TRUE |
| 13114.sandin.54.s002 | 3780 | 58079 | 43 | 81363 | TRUE | TRUE |
| 13114.sandin.54.s003 | 46545 | 56774 | 38316 | 87031 | TRUE | TRUE |
| 13114.sandin.54.s004 | 38861 | 28233 | 275 | 16412519 | TRUE | TRUE |
| 13114.sandin.54.s005 | 2045 | 35596 | 34 | 341885 | TRUE | TRUE |
| 13114.sandin.54.s006 | 23390 | 92112 | 7999 | 341534 | TRUE | TRUE |
| 13114.sandin.54.s007 | 26079 | 46491 | 266 | 3155732 | TRUE | TRUE |
| 13114.sandin.54.s008 | 31025 | 71882 | 2073 | 6575599 | TRUE | TRUE |
| 13114.sandin.54.s009 | 12220 | 2772 | 3 | 11528780 | TRUE | TRUE |

| sample name | 16S | 18S | ITS | shotgun metagenomic | lc-msms | gcms |
|---|---|---|---|---|---|---|
| 13114.sandin.54.s010 | 53625 | 109765 | 476 | 11669043 | TRUE | TRUE |
| 13114.sandin.54.s011 | 16694 | 67801 | 70 | 11728272 | TRUE | TRUE |
| 13114.sandin.54.s012 | 26033 | 91888 | 13 | 7288214 | TRUE | TRUE |
| 13114.sandin.54.s013 | 61046 | 17408 | 43 | 3137915 | TRUE | TRUE |
| 13114.sandin.54.s014 | 52799 | 141004 | 30 | 13416431 | TRUE | TRUE |
| 13114.sandin.54.s015 | 65368 | 37861 | 226 | 13146364 | FALSE | TRUE |
| 13114.schmidt.56.s001 | 52349 | FALSE | 1 | 94361 | TRUE | TRUE |
| 13114.schmidt.56.s002 | 18 | FALSE | FALSE | 636447 | TRUE | TRUE |
| 13114.schmidt.56.s003 | 26089 | FALSE | FALSE | 2168909 | TRUE | TRUE |
| 13114.schmidt.56.s004 | 28347 | FALSE | 1 | 912 | TRUE | TRUE |
| 13114.schmidt.56.s005 | 9 | FALSE | 583 | 13132 | TRUE | TRUE |
| 13114.schmidt.56.s006 | 21 | FALSE | FALSE | 8765 | TRUE | TRUE |
| 13114.schmidt.56.s007 | 33325 | FALSE | 4 | 1481351 | TRUE | TRUE |
| 13114.schmidt.56.s008 | 36 | FALSE | 1 | 285 | TRUE | TRUE |
| 13114.schmidt.56.s009 | 2 | FALSE | 4 | 239 | TRUE | TRUE |
| 13114.schmidt.56.s010 | 1688 | FALSE | FALSE | 1720 | TRUE | TRUE |
| 13114.schmidt.56.s011 | 21 | FALSE | 7 | 8598 | TRUE | TRUE |
| 13114.schmidt.56.s012 | 19 | FALSE | FALSE | 9125336 | TRUE | TRUE |
| 13114.schmidt.56.s013 | 16 | FALSE | FALSE | 53861 | TRUE | TRUE |
| 13114.schmidt.56.s014 | 46729 | FALSE | FALSE | 10667877 | TRUE | TRUE |
| 13114.schmidt.56.s015 | 35982 | FALSE | FALSE | 2604 | TRUE | TRUE |
| 13114.schmidt.56.s016 | 35917 | FALSE | FALSE | 4311 | TRUE | TRUE |
| 13114.schmidt.56.s017 | 16 | FALSE | FALSE | 2772855 | TRUE | TRUE |
| 13114.schmidt.56.s018 | 15 | FALSE | 30108 | 644 | TRUE | TRUE |
| 13114.schmidt.56.s019 | 24 | FALSE | FALSE | 13491 | TRUE | TRUE |
| 13114.schmidt.56.s020 | 33805 | FALSE | 3 | 5100716 | TRUE | TRUE |
| 13114.schmidt.56.s021 | 4 | FALSE | 1 | 2788 | TRUE | TRUE |
| 13114.schmidt.56.s022 | 44433 | FALSE | FALSE | 14826450 | TRUE | TRUE |
| 13114.schmidt.56.s023 | 21 | FALSE | 8 | 14166965 | TRUE | TRUE |
| 13114.schmidt.56.s024 | 45271 | FALSE | FALSE | 10658566 | TRUE | TRUE |
| 13114.schmidt.56.s025 | 45429 | FALSE | 26370 | 2446447 | TRUE | TRUE |
| 13114.schmidt.56.s026 | 43139 | FALSE | FALSE | 10306744 | TRUE | TRUE |
| 13114.seedorf.71.s001 | 15 | FALSE | FALSE | 8184473 | TRUE | TRUE |
| 13114.seedorf.71.s002 | 42182 | FALSE | FALSE | 8094726 | TRUE | TRUE |
| 13114.seedorf.71.s003 | 14 | FALSE | FALSE | 9825419 | TRUE | TRUE |
| 13114.seedorf.71.s004 | 48389 | FALSE | 1 | 13799760 | TRUE | TRUE |
| 13114.seedorf.71.s005 | 28235 | FALSE | 20 | 3807 | TRUE | TRUE |
| 13114.seedorf.71.s006 | 36633 | FALSE | FALSE | 10941480 | TRUE | TRUE |
| 13114.seedorf.71.s007 | 34132 | FALSE | 1 | 7979280 | TRUE | TRUE |
| 13114.seedorf.71.s008 | 35820 | FALSE | 1 | 129675 | TRUE | TRUE |
| 13114.seedorf.71.s009 | 14 | FALSE | FALSE | 25615623 | TRUE | TRUE |
| 13114.seedorf.71.s010 | 18 | FALSE | 1 | 24382359 | TRUE | TRUE |
| 13114.seedorf.71.s011 | 37567 | FALSE | 3 | 12425924 | TRUE | TRUE |
| 13114.seedorf.71.s012 | 14 | FALSE | FALSE | 12101419 | TRUE | TRUE |

| sample name | 16S | 18S | ITS | shotgun metagenomic | lc-msms | gcms |
|---|---|---|---|---|---|---|
| 13114.seedorf.71.s013 | 14 | FALSE | FALSE | 1335048 | TRUE | TRUE |
| 13114.seedorf.71.s014 | 18 | FALSE | FALSE | 11959291 | TRUE | TRUE |
| 13114.shade.23.s001 | 43750 | FALSE | 2 | 192787 | TRUE | TRUE |
| 13114.shade.23.s002 | 20 | FALSE | 19753 | 51918861 | TRUE | TRUE |
| 13114.shade.23.s003 | 37528 | FALSE | 1 | 10068612 | TRUE | TRUE |
| 13114.shade.23.s004 | 31770 | FALSE | FALSE | 12531432 | TRUE | TRUE |
| 13114.shade.23.s005 | 46546 | FALSE | FALSE | 87049 | TRUE | TRUE |
| 13114.shade.23.s006 | 33410 | FALSE | 3 | 3179851 | TRUE | TRUE |
| 13114.shade.23.s007 | 40448 | FALSE | 1 | 10148232 | TRUE | TRUE |
| 13114.shade.23.s008 | 44811 | FALSE | FALSE | 34836236 | TRUE | TRUE |
| 13114.shade.23.s009 | 24 | FALSE | 1 | 11401230 | TRUE | TRUE |
| 13114.shade.23.s010 | 32653 | FALSE | FALSE | 13513034 | TRUE | TRUE |
| 13114.smith.39.s001 | 44302 | 57160 | 113659 | 3315790 | FALSE | TRUE |
| 13114.smith.39.s002 | 90558 | 155336 | 22657 | 3110036 | FALSE | TRUE |
| 13114.smith.39.s003 | 38990 | 47616 | 1605 | 14384004 | FALSE | TRUE |
| 13114.smith.39.s004 | 62742 | 147967 | 71568 | 489307 | FALSE | TRUE |
| 13114.smith.39.s005 | 121068 | 132329 | 17180 | 14064295 | FALSE | TRUE |
| 13114.smith.39.s006 | 45432 | 112562 | 11078 | 1144642 | FALSE | TRUE |
| 13114.smith.39.s007 | 87514 | 50995 | 6797 | 8007683 | FALSE | TRUE |
| 13114.smith.39.s008 | 45967 | 154343 | 7897 | 230718 | FALSE | TRUE |
| 13114.song.51.s001 | 51754 | 129795 | 41327 | 45791 | TRUE | TRUE |
| 13114.song.51.s002 | 63014 | 183927 | 7828 | 9599567 | TRUE | TRUE |
| 13114.song.51.s003 | 173609 | 106045 | 180417 | 11282720 | TRUE | TRUE |
| 13114.song.51.s004 | 143397 | 134737 | 6193 | 9256687 | TRUE | TRUE |
| 13114.song.51.s005 | 91310 | 55832 | 115034 | 16701488 | TRUE | TRUE |
| 13114.song.51.s006 | 98848 | 101895 | 50925 | 11926204 | TRUE | TRUE |
| 13114.song.51.s007 | 29238 | 112072 | 83503 | 11830826 | TRUE | TRUE |
| 13114.song.51.s008 | 164738 | 117460 | 134397 | 10615995 | TRUE | TRUE |
| 13114.song.51.s009 | 76572 | 132926 | 75511 | 9911406 | TRUE | TRUE |
| 13114.song.51.s010 | 77850 | 14557 | 886 | 9965473 | TRUE | TRUE |
| 13114.song.51.s011 | 112121 | 133119 | 123308 | 13405644 | TRUE | TRUE |
| 13114.song.51.s012 | 65034 | 91810 | 101941 | 11586658 | TRUE | TRUE |
| 13114.song.51.s013 | 72319 | 46149 | 3984 | 7878000 | TRUE | TRUE |
| 13114.song.51.s014 | 68247 | 66181 | 193540 | 9917866 | TRUE | TRUE |
| 13114.song.51.s015 | 105442 | 139328 | 1986 | 9689005 | TRUE | TRUE |
| 13114.song.51.s016 | 80199 | 104199 | 244405 | 12416801 | TRUE | TRUE |
| 13114.song.51.s017 | 95174 | 66005 | 11049 | 2321800 | TRUE | TRUE |
| 13114.song.51.s018 | 90986 | 119132 | 278953 | 13271710 | TRUE | TRUE |
| 13114.song.51.s019 | 120336 | 165799 | 60746 | 13545111 | TRUE | TRUE |
| 13114.song.51.s020 | 44900 | 60811 | 3118 | 7186537 | TRUE | TRUE |
| 13114.song.52.s001 | 117456 | 196782 | 508 | 11916185 | TRUE | TRUE |
| 13114.song.52.s002 | 72470 | 168922 | 194851 | 11371498 | TRUE | TRUE |
| 13114.song.52.s003 | 41452 | 78995 | 860 | 12877454 | TRUE | TRUE |
| 13114.song.52.s004 | 98206 | 133270 | 219659 | 3528590 | TRUE | TRUE |

| sample name | 16S | 18S | ITS | shotgun metagenomic | lc-msms | gcms |
|---|---|---|---|---|---|---|
| 13114.song.52.s005 | 47930 | 60121 | 22102 | 10699551 | TRUE | TRUE |
| 13114.song.52.s006 | 79926 | 63673 | 36454 | 9904615 | TRUE | TRUE |
| 13114.song.52.s007 | 39789 | 109710 | 514 | 12807221 | TRUE | TRUE |
| 13114.song.52.s008 | 112389 | 135865 | 37818 | 4169899 | TRUE | TRUE |
| 13114.song.52.s009 | 123455 | 205111 | 15854 | 13219237 | TRUE | TRUE |
| 13114.song.52.s010 | 45870 | 73536 | 215596 | 11857247 | TRUE | TRUE |
| 13114.song.52.s011 | 36193 | 304824 | 140581 | 11512609 | TRUE | TRUE |
| 13114.song.52.s012 | 75450 | 37263 | 197 | 11322381 | TRUE | TRUE |
| 13114.song.52.s013 | 94467 | 139501 | 108795 | 3568636 | TRUE | TRUE |
| 13114.song.52.s014 | 131411 | 223557 | 265130 | 11619418 | TRUE | TRUE |
| 13114.song.52.s015 | 75201 | 95865 | 198249 | 4591062 | TRUE | TRUE |
| 13114.song.53.s001 | 129960 | 170848 | 289920 | 11823139 | TRUE | TRUE |
| 13114.song.53.s002 | 85118 | 73519 | 94922 | 9588159 | TRUE | TRUE |
| 13114.song.53.s003 | 66134 | 37806 | 22443 | 7613039 | TRUE | TRUE |
| 13114.song.53.s004 | 156677 | 158334 | 298933 | 9965098 | TRUE | TRUE |
| 13114.song.53.s005 | 43278 | 111685 | 1677 | 13385858 | TRUE | TRUE |
| 13114.song.53.s006 | 128548 | 158834 | 172922 | 10931310 | TRUE | TRUE |
| 13114.song.53.s007 | 36515 | 103580 | 30035 | 3140674 | TRUE | TRUE |
| 13114.song.53.s008 | 79530 | 57708 | 4541 | 14933707 | TRUE | TRUE |
| 13114.song.53.s009 | 74092 | 68284 | 12472 | 8101105 | TRUE | TRUE |
| 13114.song.53.s010 | 84318 | 114255 | 19581 | 9170795 | TRUE | TRUE |
| 13114.song.53.s011 | 60361 | 100609 | 13865 | 12845930 | TRUE | TRUE |
| 13114.song.53.s012 | 95206 | 131118 | 34770 | 6578925 | TRUE | TRUE |
| 13114.song.53.s013 | 50032 | 100255 | 151649 | 5044015 | TRUE | TRUE |
| 13114.song.53.s014 | 66448 | 72111 | 69733 | 16098952 | TRUE | TRUE |
| 13114.song.53.s015 | 57860 | 100412 | 62303 | 32439670 | TRUE | TRUE |
| 13114.song.75.s001 | 26733 | 46072 | 154 | 37456 | TRUE | TRUE |
| 13114.song.75.s002 | 92325 | 146535 | 15947 | 190275 | TRUE | TRUE |
| 13114.song.75.s003 | 36180 | 162495 | 1366 | 14285879 | TRUE | TRUE |
| 13114.song.75.s004 | 170595 | 93158 | 2679 | 13703711 | TRUE | TRUE |
| 13114.stegen.36.s001 | 33846 | FALSE | 2 | 5915529 | FALSE | FALSE |
| 13114.stegen.36.s002 | 18 | FALSE | 4 | 13750380 | TRUE | TRUE |
| 13114.stegen.36.s003 | 16 | FALSE | 2 | 2060716 | TRUE | TRUE |
| 13114.stegen.36.s004 | 41252 | FALSE | 2 | 14778049 | TRUE | TRUE |
| 13114.stegen.36.s005 | 11 | FALSE | FALSE | 27569 | TRUE | TRUE |
| 13114.stegen.36.s006 | 14 | FALSE | FALSE | 13582462 | TRUE | TRUE |
| 13114.stegen.36.s007 | 43371 | FALSE | 1 | 7860422 | FALSE | FALSE |
| 13114.stegen.36.s008 | 29296 | FALSE | 2 | 10367303 | TRUE | TRUE |
| 13114.stegen.36.s009 | 17 | FALSE | 1 | 8814690 | TRUE | TRUE |
| 13114.stegen.36.s010 | 10 | FALSE | 8 | 85408880 | FALSE | FALSE |
| 13114.stegen.36.s011 | 4 | FALSE | FALSE | 19593515 | TRUE | TRUE |
| 13114.stegen.36.s012 | 54307 | FALSE | FALSE | 12966856 | FALSE | FALSE |
| 13114.stegen.36.s013 | 37523 | FALSE | FALSE | 8416577 | TRUE | TRUE |
| 13114.stegen.36.s014 | 16 | FALSE | 1 | 1441453 | TRUE | TRUE |

| sample name | 16S | 18S | ITS | shotgun metagenomic | lc-msms | gcms |
|---|---|---|---|---|---|---|
| 13114.stegen.36.s015 | 20 | FALSE | FALSE | 6608514 | TRUE | TRUE |
| 13114.stegen.37.s001 | 30010 | FALSE | FALSE | 1152 | FALSE | FALSE |
| 13114.stegen.37.s002 | 7 | FALSE | 4 | 7089 | FALSE | FALSE |
| 13114.stegen.37.s003 | 24700 | FALSE | 5 | 12741 | FALSE | FALSE |
| 13114.stegen.37.s004 | 22 | FALSE | 25 | 9519449 | FALSE | FALSE |
| 13114.stegen.37.s005 | 16 | FALSE | FALSE | 1894 | FALSE | FALSE |
| 13114.stegen.37.s006 | 26 | FALSE | FALSE | 4395 | FALSE | FALSE |
| 13114.stegen.37.s007 | 8 | FALSE | 3 | 12855 | FALSE | FALSE |
| 13114.stegen.37.s008 | 42667 | FALSE | 27375 | 252276 | FALSE | FALSE |
| 13114.stegen.37.s009 | 16 | FALSE | 3 | 31123 | FALSE | FALSE |
| 13114.stegen.37.s010 | 37754 | FALSE | 11 | 73521 | FALSE | FALSE |
| 13114.stegen.38.s001 | 33375 | FALSE | FALSE | 14304267 | FALSE | FALSE |
| 13114.stegen.38.s002 | 12 | FALSE | FALSE | 18943476 | FALSE | FALSE |
| 13114.stegen.38.s003 | 19 | FALSE | FALSE | 2529579 | FALSE | FALSE |
| 13114.stegen.38.s004 | 18 | FALSE | FALSE | 14974 | FALSE | FALSE |
| 13114.stegen.38.s005 | 44701 | FALSE | 29767 | 10617546 | FALSE | FALSE |
| 13114.stegen.38.s006 | 53485 | FALSE | FALSE | 15111856 | TRUE | TRUE |
| 13114.stegen.38.s007 | 35 | FALSE | FALSE | 14184897 | FALSE | FALSE |
| 13114.stegen.38.s008 | 30377 | FALSE | 1 | 216830 | FALSE | FALSE |
| 13114.stegen.38.s009 | 44604 | FALSE | FALSE | 9147986 | FALSE | FALSE |
| 13114.stegen.38.s010 | 21 | FALSE | 1 | 731147 | FALSE | FALSE |
| 13114.stegen.38.s011 | 39636 | FALSE | 1 | 10614553 | FALSE | FALSE |
| 13114.stegen.38.s012 | 19 | FALSE | FALSE | 17163593 | FALSE | FALSE |
| 13114.stegen.38.s013 | 47859 | FALSE | FALSE | 10568420 | FALSE | FALSE |
| 13114.stegen.38.s014 | 39665 | FALSE | FALSE | 9971737 | FALSE | FALSE |
| 13114.stegen.38.s015 | 37986 | FALSE | 2 | 42113721 | FALSE | FALSE |
| 13114.stegen.38.s016 | 28 | FALSE | 1 | 7354435 | FALSE | FALSE |
| 13114.stegen.38.s017 | 69367 | FALSE | FALSE | 13389374 | FALSE | FALSE |
| 13114.stegen.38.s018 | 22 | FALSE | FALSE | 13106504 | FALSE | FALSE |
| 13114.stewart.26.s001 | 58946 | 126419 | 551 | 6763812 | FALSE | FALSE |
| 13114.stewart.26.s002 | 31515 | 76356 | 445 | 4859667 | FALSE | FALSE |
| 13114.stewart.26.s003 | 77615 | 109418 | 819 | 784753 | FALSE | FALSE |
| 13114.stewart.26.s004 | 44959 | 84331 | 397 | 2345198 | FALSE | FALSE |
| 13114.stewart.26.s005 | 68423 | 77337 | 47 | 2239498 | FALSE | FALSE |
| 13114.stewart.26.s006 | 49247 | 84600 | 1454 | 2450444 | FALSE | FALSE |
| 13114.stewart.26.s007 | 85742 | 125996 | 1055 | 10715669 | FALSE | FALSE |
| 13114.stewart.26.s008 | 31020 | 30491 | 222 | 5150799 | FALSE | FALSE |
| 13114.tait.77.s001 | 176285 | 195820 | 131525 | 10113059 | TRUE | FALSE |
| 13114.tait.77.s002 | 135368 | 305251 | 56110 | 11632919 | TRUE | FALSE |
| 13114.tait.77.s003 | 97309 | 225635 | 97496 | 11394842 | TRUE | FALSE |
| 13114.tait.77.s004 | 63314 | 68485 | 76206 | 10239927 | TRUE | FALSE |
| 13114.tait.77.s005 | 37861 | 45718 | 26972 | 10932371 | TRUE | FALSE |
| 13114.tait.77.s006 | 59755 | 130983 | 771 | 11299283 | TRUE | FALSE |
| 13114.tait.77.s007 | 50859 | 29411 | 60498 | 10431596 | TRUE | FALSE |

| sample name | 16S | 18S | ITS | shotgun metagenomic | lc-msms | gcms |
|---|---|---|---|---|---|---|
| 13114.tait.77.s008 | 69483 | 44962 | 55088 | 9103992 | TRUE | FALSE |
| 13114.tait.77.s009 | 148271 | 130703 | 142898 | 12402315 | TRUE | FALSE |
| 13114.tait.78.s001 | 30707 | 201087 | 28944 | 15684407 | FALSE | TRUE |
| 13114.tait.78.s002 | 85120 | 275512 | 95607 | 17933885 | FALSE | TRUE |
| 13114.tait.78.s003 | 107807 | 113703 | 73053 | 16242836 | FALSE | TRUE |
| 13114.thomas.18.s001 | 73504 | 138468 | 4246 | 14486043 | TRUE | TRUE |
| 13114.thomas.18.s002 | 84287 | 92855 | 5093 | 17225613 | TRUE | TRUE |
| 13114.thomas.18.s003 | 76818 | 92410 | 4016 | 8446173 | TRUE | FALSE |
| 13114.thomas.18.s004 | 47274 | FALSE | 1 | 2993463 | FALSE | FALSE |
| 13114.thomas.18.s005 | 203 | 5798 | 20 | 3733093 | FALSE | FALSE |
| 13114.thomas.18.s006 | 9784 | 88513 | FALSE | 2698950 | FALSE | FALSE |
| 13114.thomas.18.s007 | 20892 | 96245 | 3 | 3922758 | FALSE | FALSE |
| 13114.thomas.18.s008 | 40710 | 9264 | 27971 | 6949643 | TRUE | TRUE |
| 13114.thomas.18.s009 | 30796 | 89971 | 13942 | 14048989 | TRUE | TRUE |
| 13114.thomas.18.s010 | 45999 | 64112 | 24700 | 15131275 | TRUE | FALSE |
| 13114.thomas.18.s011 | 68739 | 305 | 2888 | 3078771 | TRUE | FALSE |
| 13114.thomas.18.s012 | 65825 | 86032 | 1078 | 15290474 | TRUE | TRUE |
| 13114.thomas.18.s013 | 58193 | 67415 | 475 | 15072946 | TRUE | TRUE |
| 13114.thomas.18.s014 | 70213 | 75430 | 18 | 11284387 | TRUE | FALSE |
| 13114.thomas.18.s015 | 96659 | 14529 | 49952 | 11911035 | TRUE | FALSE |
| 13114.thomas.18.s016 | 75212 | 130565 | 5 | 13362656 | TRUE | FALSE |
| 13114.thomas.18.s017 | 12728 | 42108 | 101 | 4373058 | FALSE | FALSE |
| 13114.thomas.18.s018 | 13543 | 9395 | 168 | 4987656 | FALSE | FALSE |
| 13114.thomas.18.s019 | 83976 | 72733 | 583 | 9668093 | TRUE | TRUE |
| 13114.thomas.18.s020 | 37436 | FALSE | 3 | 5679816 | TRUE | FALSE |
| 13114.thomas.18.s021 | 31457 | 59126 | 42338 | 11869910 | TRUE | FALSE |
| 13114.thomas.18.s022 | 11713 | 99368 | 35658 | 3327080 | FALSE | FALSE |
| 13114.thomas.18.s023 | 51995 | 126422 | 72446 | 8831301 | TRUE | TRUE |
| 13114.thomas.18.s024 | 35124 | 65358 | 89225 | 12386578 | TRUE | TRUE |
| 13114.thomas.18.s025 | 85524 | 26736 | 32302 | 10546734 | TRUE | FALSE |
| 13114.thomas.18.s026 | 90852 | 143022 | 131398 | 8301723 | TRUE | FALSE |
| 13114.thomas.18.s027 | 16472 | 62946 | 122202 | 1611278 | FALSE | FALSE |
| 13114.thomas.18.s028 | 65058 | 130588 | 119235 | 14612934 | TRUE | TRUE |
| 13114.thomas.18.s029 | 93009 | 54961 | 136958 | 13101656 | TRUE | TRUE |
| 13114.thomas.18.s030 | 81264 | 80409 | 45176 | 12044207 | TRUE | FALSE |
| 13114.thomas.18.s031 | 46628 | 23401 | 3128 | 4442336 | TRUE | TRUE |
| 13114.thomas.18.s032 | 55611 | 8044 | 257 | 260 | TRUE | TRUE |
| 13114.thomas.18.s033 | 53285 | 3047 | 4832 | 7736895 | TRUE | FALSE |
| 13114.thomas.18.s034 | 1376 | 982 | 2 | 224 | TRUE | FALSE |
| 13114.thomas.18.s035 | 47671 | 24960 | 1860 | 7410489 | TRUE | TRUE |
| 13114.thomas.18.s036 | 24005 | 40198 | 879 | 6320958 | TRUE | TRUE |
| 13114.thomas.18.s037 | 54181 | 18216 | 7664 | 6973009 | TRUE | FALSE |
| 13114.thomas.18.s038 | 18852 | 58360 | 48794 | 2884821 | TRUE | TRUE |
| 13114.thomas.18.s039 | 23378 | 48339 | 42707 | 4267765 | TRUE | TRUE |

| sample name | 16S | 18S | ITS | shotgun metagenomic | lc-msms | gcms |
|---|---|---|---|---|---|---|
| 13114.thomas.18.s040 | 931 | 49020 | 12569 | 8671300 | TRUE | FALSE |
| 13114.thomas.18.s041 | 12317 | 50335 | 1035 | 2947568 | TRUE | FALSE |
| 13114.thomas.18.s042 | 16 | FALSE | 6 | 19 | TRUE | TRUE |
| 13114.thomas.18.s043 | 63840 | 7404 | 1355 | 6658083 | TRUE | TRUE |
| 13114.thomas.18.s044 | 11100 | 1 | 1 | 10276 | TRUE | FALSE |
| 13114.thomas.18.s045 | 23146 | 7274 | 47 | 5080832 | TRUE | FALSE |
| 13114.thomas.18.s046 | 72569 | 772 | 6 | 8978813 | TRUE | TRUE |
| 13114.thomas.18.s047 | 69521 | 16 | FALSE | 5288 | TRUE | TRUE |
| 13114.thomas.18.s048 | 78342 | 2908 | 24 | 8137000 | TRUE | FALSE |
| 13114.thomas.18.s049 | 75629 | 3797 | 8 | 606509 | TRUE | FALSE |
| 13114.thomas.18.s050 | 21502 | 55703 | 41033 | 5071194 | TRUE | TRUE |
| 13114.thomas.18.s051 | 33731 | 47087 | 69006 | 3220634 | TRUE | TRUE |
| 13114.thomas.18.s052 | 36662 | 48171 | 44766 | 4466524 | TRUE | FALSE |
| 13114.thomas.18.s053 | 25867 | 15484 | 31688 | 3388854 | TRUE | FALSE |
| 13114.thomas.18.s054 | 41864 | 56200 | 88153 | 4868185 | TRUE | TRUE |
| 13114.thomas.18.s055 | 48632 | 45752 | 38179 | 5814168 | TRUE | TRUE |
| 13114.thomas.18.s056 | 62011 | 72534 | 33003 | 5680949 | TRUE | FALSE |
| 13114.thomas.18.s057 | 46595 | 59227 | 71767 | 4742369 | TRUE | FALSE |
| 13114.thomas.19.s001 | 53719 | 213310 | 81 | FALSE | FALSE | FALSE |
| 13114.thomas.19.s002 | 17725 | 92728 | 1133 | 4064788 | FALSE | FALSE |
| 13114.thomas.19.s003 | 18869 | 69785 | 10525 | 2305290 | FALSE | FALSE |
| 13114.thomas.19.s004 | 19659 | 75501 | 790 | 2535132 | FALSE | FALSE |
| 13114.thomas.19.s005 | 20492 | 136104 | 24494 | FALSE | FALSE | FALSE |
| 13114.thomas.19.s006 | 39117 | 123015 | 183 | FALSE | FALSE | FALSE |
| 13114.thomas.19.s007 | 28035 | 118113 | 1296 | 2719807 | FALSE | FALSE |
| 13114.thomas.19.s008 | 12760 | 75840 | 10644 | 4005850 | FALSE | FALSE |
| 13114.thomas.19.s009 | 17304 | 72965 | 3805 | 2028345 | FALSE | FALSE |
| 13114.thomas.19.s010 | 21344 | 38446 | 11136 | 3076991 | FALSE | FALSE |
| 13114.thomas.19.s011 | 14898 | 52389 | 3587 | FALSE | FALSE | FALSE |
| 13114.thomas.19.s012 | 6398 | 102310 | 95 | FALSE | FALSE | FALSE |
| 13114.thomas.19.s013 | 18385 | 101940 | 1261 | FALSE | FALSE | FALSE |
| 13114.thomas.19.s014 | 46606 | 89826 | 2681 | 2436782 | FALSE | FALSE |
| 13114.thomas.19.s015 | 13074 | 41413 | 2832 | 4860070 | FALSE | FALSE |
| 13114.thomas.19.s016 | 18111 | 34562 | 7 | FALSE | FALSE | FALSE |
| 13114.thomas.19.s017 | 12675 | 15988 | 12865 | 2583439 | FALSE | FALSE |
| 13114.thomas.19.s018 | 18589 | 88879 | 20654 | 3019676 | FALSE | FALSE |
| 13114.thomas.19.s019 | 110 | 113833 | 1221 | 3902807 | FALSE | FALSE |
| 13114.thomas.19.s020 | 34628 | 130955 | 5194 | 2979740 | FALSE | FALSE |
| 13114.thomas.19.s021 | 14555 | 98914 | 3870 | 1943099 | FALSE | FALSE |
| 13114.thomas.19.s022 | 21712 | 28158 | 17809 | 4019987 | FALSE | FALSE |
| 13114.thomas.19.s023 | 10200 | 72289 | 2572 | 2760870 | FALSE | FALSE |
| 13114.thomas.19.s024 | 20933 | 116774 | 20963 | 3589303 | FALSE | FALSE |
| 13114.thomas.19.s025 | 19179 | 11113 | 631 | 2364314 | FALSE | FALSE |
| 13114.thomas.19.s026 | 19822 | 61462 | 11136 | 2809753 | FALSE | FALSE |

| sample name | 16S | 18S | ITS | shotgun metagenomic | lc-msms | gcms |
|---|---|---|---|---|---|---|
| 13114.thomas.19.s027 | FALSE | FALSE | FALSE | FALSE | TRUE | TRUE |
| 13114.thomas.19.s028 | FALSE | FALSE | FALSE | FALSE | TRUE | TRUE |
| 13114.thomas.19.s029 | FALSE | FALSE | FALSE | FALSE | TRUE | TRUE |
| 13114.thomas.19.s030 | FALSE | FALSE | FALSE | FALSE | TRUE | TRUE |
| 13114.thomas.19.s031 | FALSE | FALSE | FALSE | FALSE | TRUE | TRUE |
| 13114.thomas.19.s032 | FALSE | FALSE | FALSE | FALSE | TRUE | TRUE |
| 13114.thomas.19.s033 | FALSE | FALSE | FALSE | FALSE | TRUE | TRUE |
| 13114.thomas.19.s034 | FALSE | FALSE | FALSE | FALSE | TRUE | TRUE |
| 13114.thomas.19.s035 | FALSE | FALSE | FALSE | FALSE | TRUE | TRUE |
| 13114.thomas.19.s036 | FALSE | FALSE | FALSE | FALSE | TRUE | TRUE |
| 13114.thomas.19.s037 | FALSE | FALSE | FALSE | FALSE | TRUE | TRUE |
| 13114.thomas.19.s038 | FALSE | FALSE | FALSE | FALSE | TRUE | TRUE |
| 13114.thomas.19.s039 | FALSE | FALSE | FALSE | FALSE | TRUE | TRUE |
| 13114.thomas.19.s040 | FALSE | FALSE | FALSE | FALSE | TRUE | TRUE |
| 13114.thomas.19.s041 | FALSE | FALSE | FALSE | FALSE | TRUE | TRUE |
| 13114.thomas.19.s042 | FALSE | FALSE | FALSE | FALSE | TRUE | TRUE |
| 13114.thomas.19.s043 | FALSE | FALSE | FALSE | FALSE | TRUE | TRUE |
| 13114.thomas.19.s044 | FALSE | FALSE | FALSE | FALSE | TRUE | TRUE |
| 13114.thomas.19.s045 | FALSE | FALSE | FALSE | FALSE | TRUE | TRUE |
| 13114.thomas.19.s046 | FALSE | FALSE | FALSE | FALSE | TRUE | TRUE |
| 13114.thomas.19.s047 | FALSE | FALSE | FALSE | FALSE | TRUE | TRUE |
| 13114.thomas.19.s048 | FALSE | FALSE | FALSE | FALSE | TRUE | TRUE |
| 13114.thomas.19.s049 | FALSE | FALSE | FALSE | FALSE | TRUE | TRUE |
| 13114.thomas.19.s050 | 40665 | 68147 | 874 | 1849651 | FALSE | FALSE |
| 13114.thomas.19.s051 | 44391 | 35136 | 3047 | 1550900 | FALSE | FALSE |
| 13114.thomas.19.s052 | 36922 | 72697 | 13645 | 5572587 | FALSE | FALSE |
| 13114.thomas.19.s053 | 23445 | 46673 | 12532 | 5315000 | FALSE | FALSE |
| 13114.thomas.19.s054 | 19635 | 47038 | 373 | 1926094 | FALSE | FALSE |
| 13114.thomas.19.s055 | 18263 | 28380 | 2199 | 3476377 | FALSE | FALSE |
| 13114.thomas.19.s056 | 39193 | 50963 | 12344 | 2832719 | FALSE | FALSE |
| 13114.thomas.19.s057 | 34571 | 49414 | 11598 | 2743245 | FALSE | FALSE |
| 13114.thomas.19.s058 | 27036 | 42139 | 47 | 1155664 | FALSE | FALSE |
| 13114.thomas.19.s059 | 53472 | 40307 | 1638 | 1290832 | FALSE | FALSE |
| 13114.thomas.19.s060 | 28208 | 10681 | 22862 | 5085123 | FALSE | FALSE |
| 13114.thomas.19.s061 | 27523 | 32374 | 506 | 1115238 | FALSE | FALSE |
| 13114.thomas.19.s062 | 26323 | 44438 | 694 | 1637588 | FALSE | FALSE |
| 13114.thomas.19.s063 | 20720 | 36982 | 52 | 1614656 | FALSE | FALSE |
| 13114.thomas.19.s064 | 60239 | 53022 | 5336 | 4254636 | FALSE | FALSE |
| 13114.tucker.58.s001 | 518 | 8359 | 18 | 1382040 | TRUE | TRUE |
| 13114.tucker.58.s002 | 3849 | 32650 | 28 | 13903 | TRUE | TRUE |
| 13114.tucker.58.s003 | 2711 | 10859 | 79 | 431853 | TRUE | TRUE |
| 13114.tucker.58.s004 | 2227 | 40580 | 750 | 3344 | TRUE | FALSE |
| 13114.tucker.58.s005 | 3362 | 105244 | 10 | 1810 | TRUE | TRUE |
| 13114.tucker.58.s006 | 20252 | 17228 | 289 | 23164 | TRUE | FALSE |

| sample name | 16S | 18S | ITS | shotgun metagenomic | lc-msms | gcms |
|---|---|---|---|---|---|---|
| 13114.tucker.58.s007 | 5911 | 19248 | 2328 | 2507 | TRUE | TRUE |
| 13114.tucker.58.s008 | 11989 | 6223 | 860 | 288948 | TRUE | FALSE |
| 13114.tucker.58.s009 | 3673 | 117619 | 19 | 1064 | TRUE | TRUE |
| 13114.tucker.58.s010 | 18817 | 40252 | 2822 | 338013 | TRUE | FALSE |
| 13114.tucker.58.s011 | 22255 | 27326 | 1751 | 212818 | TRUE | FALSE |
| 13114.tucker.58.s012 | 38010 | 3458 | 128 | 12869244 | TRUE | FALSE |
| 13114.tucker.58.s013 | 58453 | 61467 | 683 | 1766941 | TRUE | FALSE |
| 13114.tucker.58.s014 | 28598 | 72213 | 1298 | 1685615 | TRUE | FALSE |
| 13114.tucker.58.s015 | 2018 | 46363 | 80 | 518096 | TRUE | FALSE |
| 13114.tucker.58.s016 | 47790 | 64016 | 134 | 14634036 | TRUE | FALSE |
| 13114.uren.82.s001 | 85493 | 104485 | 158544 | 10716982 | TRUE | TRUE |
| 13114.uren.82.s002 | 75057 | 74545 | 96685 | 1952239 | TRUE | TRUE |
| 13114.uren.82.s003 | 43489 | 73691 | 136403 | 566665 | TRUE | TRUE |
| 13114.uren.82.s004 | 55914 | 57241 | 80241 | 11826730 | TRUE | TRUE |
| 13114.uren.82.s005 | 46984 | 57537 | 114876 | 10363798 | TRUE | TRUE |
| 13114.uren.82.s006 | 40977 | 107455 | 118865 | 3026021 | TRUE | TRUE |
| 13114.uren.82.s007 | 3718 | 28130 | 15688 | 227402 | TRUE | TRUE |
| 13114.uren.82.s008 | 31428 | 79556 | 46438 | 2678905 | TRUE | TRUE |
| 13114.uren.82.s009 | 27482 | 75607 | 96394 | 2803936 | TRUE | TRUE |
| 13114.uren.82.s010 | 87652 | 106606 | 114694 | 10637686 | TRUE | TRUE |
| 13114.uren.82.s011 | 144361 | 133867 | 279003 | 213548 | TRUE | TRUE |
| 13114.uren.82.s012 | 90990 | 169327 | 158161 | 14029443 | TRUE | TRUE |
| 13114.zaneveld.9.s001 | 11198 | 200846 | 9 | 13822439 | TRUE | TRUE |
| 13114.zaneveld.9.s002 | 20293 | 59153 | 438 | 776290 | TRUE | TRUE |
| 13114.zaneveld.9.s003 | 14076 | 90608 | 630 | 11373645 | TRUE | FALSE |
| 13114.zaneveld.9.s004 | 39400 | 121537 | 346251 | 14374017 | TRUE | TRUE |
| 13114.zaneveld.9.s005 | 85621 | 96521 | 408984 | 13905083 | TRUE | TRUE |
| 13114.zaneveld.9.s006 | 92216 | 103546 | 364987 | 11461433 | TRUE | FALSE |
| 13114.zaneveld.9.s007 | 94689 | 142108 | 391996 | 9344994 | TRUE | TRUE |
| 13114.zaneveld.9.s008 | 32179 | 101954 | 204394 | 9868060 | TRUE | TRUE |
| 13114.zaneveld.9.s009 | 48000 | 94154 | 230427 | 13543100 | TRUE | FALSE |
| 13114.zaneveld.9.s010 | 15117 | 82230 | 26 | 4876791 | TRUE | TRUE |
| 13114.zaneveld.9.s011 | 49097 | 194183 | 286 | 292797 | TRUE | TRUE |
| 13114.zaneveld.9.s012 | 50503 | 150122 | 63711 | 8787332 | TRUE | FALSE |
| 13114.zaneveld.9.s013 | 20204 | 105052 | 76084 | 11493363 | TRUE | TRUE |
| 13114.zaneveld.9.s014 | 27559 | 36828 | 15169 | 8277244 | TRUE | TRUE |
| 13114.zaneveld.9.s015 | 13945 | 69045 | 26216 | 15576799 | TRUE | TRUE |
| 13114.zaneveld.9.s016 | 9557 | 48930 | 1270 | 6598171 | TRUE | TRUE |
| 13114.zaneveld.9.s017 | 27620 | 75101 | 95242 | 10541895 | TRUE | TRUE |
| 13114.zaneveld.9.s018 | 30242 | 32998 | 290121 | 9865574 | TRUE | TRUE |
| 13114.zaneveld.9.s019 | 31098 | 159503 | 159998 | 1415449 | TRUE | TRUE |
| 13114.zaneveld.9.s020 | 41944 | 118074 | 155281 | 14546140 | TRUE | TRUE |
| 13114.zaneveld.9.s021 | 62443 | 101597 | 22765 | 13241043 | TRUE | FALSE |
| 13114.zaneveld.9.s022 | 124632 | 254091 | 82301 | 8113820 | TRUE | FALSE |
